# Toward Essential Oil Stewardship: Strain-Resolved Evaluation of Thyme Oil Activity Against *Pseudomonas aeruginosa*

**DOI:** 10.1101/2025.06.13.659493

**Authors:** Malwina Brożyna, Zuzanna Stępnicka, Natalia Tymińska, Bartłomiej Dudek, Katarzyna Kapczyńska, Adam Matkowski, Yanfang Sun, Adam Junka

## Abstract

The rising interest in essential oils (EOs) as antimicrobial agents demands evaluation frameworks that move beyond anecdotal efficacy and toward structured, reproducible assessment. In this study, we examined the strain-dependent response of *Pseudomonas aeruginosa* to Pharmacopoeia-grade Thyme Essential Oil (TEO) or polyhexamethylene biguanide antiseptic (PHMB), using a panel of ten genetically diversified strains in planktonic and biofilm form and by complementary *in vitro* models. Despite uniform test conditions, we observed striking inter-strain variability: TEO Minimal Inhibitory Concentrations (MICs) differed by up to 1000-fold, and biofilm susceptibility profiles ranged from full tolerance to near-complete eradication. Notably, strains with low metabolic activity and sparse cell populations—but high matrix biomass—exhibited reduced responsiveness to TEO, while susceptibility to PHMB was more consistent, though not absolute. These findings highlight the critical influence of both microbial phenotype and agent formulation on antimicrobial outcomes. Rather than framing EOs as superior or inferior alternatives, our results advocate for their integration into a stewardship paradigm—one that values standardization, model-based evaluation, and informed formulation. In this context, we position essential oil stewardship not as a constraint, but as a necessary evolution for their credible inclusion in antimicrobial strategies.

## Introduction

*Pseudomonas aeruginosa* is a genomically complex and phenotypically diverse Gram-negative pathogen, notorious for its intrinsic resistance, environmental resilience, and clinical adaptability. Its remarkable success across ecological niches — from soil and water to medical devices and chronically infected tissues — is rooted in a large and dynamic genome composed of a conserved core and an expansive accessory repertoire (1,2). Horizontal gene transfer, genomic rearrangements, and microevolutionary processes contribute to a non-clonal population structure, in which even closely related isolates may differ markedly in virulence, metabolic capacity, biofilm behavior, and antimicrobial susceptibility (2–4).

These features pose formidable challenges to infection control. *P. aeruginosa* is listed among the World Health Organization’s highest-priority pathogens due to its pervasive multidrug resistance and its ability to thrive in biofilms — structured, matrix-embedded communities that protect the bacteria from antimicrobials and host immune responses (5,6).While most experimental studies focus on reference strains such as PAO1 or PA14, clinical outcomes are ultimately shaped by the specific traits of the infecting isolate (2). Yet, strain-level variability is often overlooked in antimicrobial research, where either a single strain is profiled using advanced techniques, or a broader panel of strains is tested with lesser methodological depth (7).

This gap arises from both practical and conceptual constraints. In-depth phenotypic analyses — including biofilm imaging, viability assays, and molecular profiling — are time- and resource-intensive, making comprehensive evaluation of multiple strains logistically difficult (8). As a result, studies frequently default to using a well-characterized reference strain for comparability and genetic tractability. Conversely, larger isolate panels are commonly assessed using only limited spectrum of readouts such as Minimal Inhibitory Concentrations (MIC) or biomass measurements, which fail to capture the functional heterogeneity within species. Compounding this is the prevailing assumption that findings from a model strain can be generalized across the species — an assumption increasingly at odds with accumulating evidence. This methodological bottleneck has impeded progress in understanding intra-species variability, particularly in response to complex, multicomponent agents such as essential oils (EOs) (9). Much like *P. aeruginosa* itself — with its multifactorial resistance mechanisms and strain-specific phenotypes — EOs are not uniform chemical entities but complex, multicomponent mixtures (10). Even when pharmacopeial standards define their minimum content of major constituents, such as thymol in Thyme EO, these oils typically contain between several and over a dozen bioactive compounds that interact through synergistic, additive, or even antagonistic mechanisms (11,12).

*Thymus vulgaris* L. – common thyme – the source material for the pharmacopeial material is not only a medicinal aromatic plant but is also a commonly used culinary spice. Cultivated worldwide, this plant is genetically diverse with numerous races, genotypes and cultivars that, along with environmental and agronomic factors may significantly modulate essential oil profile, including minor compounds (13). These may also add some degree of uncertainty in desired activity, despite conforming to quality standards (14). Moreover, European Pharmacopeia and EMA HMPC (European Medicines Agency, Herbal Medicinal Products Committee) monograph accept also *T.zygis* L. or a mixture of those as valid herbal material (15). *T. zygis* (red thyme) is less widespread in cultivation but also versatile in use and variable in composition (16). *Thymi aetheroleum* (Thyme Essential Oil) according to European Pharmacopoeia shall contain between 37-55 % thymol. The typically found minor (non-thymol) compounds of T-EO include: carvacrol 0.5% to 5.5%, carvacrol methyl ether 0.05% to 1.5%, p-cymene 14.0% to 28.0%, linalool 1.5% to 6.5%, β-myrcene 1.0% to 3.0%, α-terpinene 0.9% to 2.6%, γ-terpinene 4.0% to 12.0%, terpinen-4-ol 0.1% to 2.5%, α-thujene 0.2% to 1.5%, with an admixture of a sesquiterpene β-caryophyllene (15).

It makes the prediction of a real-world pharmacological activity even more challenging if only based on the major compound and strengthens the importance of EO stewardship.

Moreover, their antimicrobial activity is not limited to the liquid phase: volatile fractions can exert effects on planktonic and biofilm-forming bacteria at a distance, further complicating mechanistic interpretation and reproducibility (17). The dual-phase nature of EO activity — airborne and contact-dependent — challenges classical testing paradigms developed for water-soluble antibiotics and necessitates more nuanced, multi-layered assessment strategies (18).

Further complicating the landscape is the methodological heterogeneity inherent to *in vitro* biofilm research (19). Unlike planktonic assays, where standardized protocols exist for determining MICs, or zones of growth inhibition, biofilm studies are subject to considerable variability in experimental design (20,21). Factors such as the duration of biofilm maturation, the nature of the surface (e.g., polystyrene, glass, biomaterials), the composition of the culture medium, and even the strain-specific propensity for biofilm formation can profoundly influence outcomes (19,22). Importantly, the choice of medium should reflect the chemical and nutritional composition of the infection site, as nutrient-rich laboratory broths (e.g., Luria Broth or Tryptic Soya Broth) poorly mimic the physiological conditions found in wound exudate, respiratory mucus, or urinary secretions (23–25). Another critical but often underestimated variable in biofilm research is the nature of the surface on which the biofilm *in vitro* develops. While most standard assays employ inert, aforementioned flat surfaces such as polystyrene or glass, these materials poorly replicate the structural and biochemical properties of host tissues (26). In contrast, porous and biologically relevant matrices — such as collagen-coated surfaces or bio-nanocellulose — are considered to better mimic the extracellular environment encountered during infection, providing more accurate insights into biofilm architecture, antimicrobial penetration, and host–pathogen interactions (27,28). Biofilms formed on such tissue-like substrates often exhibit greater density, deeper stratification, and altered metabolic gradients, which can significantly affect susceptibility profiles. Therefore, the choice of surface is not merely a technical detail but a major determinant of experimental outcome, particularly when evaluating agents intended for clinical application in wound, mucosal, or implant-associated infections (29).

Additionally, antimicrobial agents may be introduced at different stages — either during biofilm development (preventive models) or after its establishment (therapeutic models) and their activity assessed *via* a wide range of readouts: biomass staining (e.g., crystal violet), metabolic activity (e.g., resazurin, XTT), viability (CFU (Colony-Forming Unit) counts, live/dead staining), or imaging (confocal microscopy, scanning electron microscope) (30). Each of these endpoints captures distinct dimensions of biofilm integrity and susceptibility, making cross-study comparisons challenging. Thus, the complexity of the biological target and the therapeutic agent is further compounded by the methodological plasticity of the models themselves — underscoring the need for integrative, multi-parametric approaches when evaluating biofilm-active compounds (8).

Taken together, the evaluation of antimicrobial EOs’ strategies against *P. aeruginosa* biofilms *in vitro* is confounded by at least three major overlapping layers of complexity: (i) the extensive intra-species variability in biofilm formation and resistance phenotypes; (ii) the multicomponent and phase-diverse nature of EOs, whose activity depends not only on chemical composition but also on volatility, solubility, and interactions between constituents; and (iii) the methodological heterogeneity of *in vitro* biofilm models, influenced by medium composition, surface properties, exposure protocols, and endpoint selection (4,17,24,27). These challenges have led to fragmented literature and hindered the translational potential of plant-based antimicrobials. Therefore, in this study, we address these limitations by integrating a genetically diverse pseudomonal strains’ panel, chemically specified TEO and a suite of complementary biofilm assessment techniques, all embedded within a rigorous statistical framework. Our approach enables a multidimensional comparison of strain-specific responses and provides a robust template for the rational preclinical evaluation of phytotherapeutic agents.

## Materials and methods

### Microorganisms

Ten strains of *Pseudomonas aeruginosa* were selected for research purposes. Eight strains included clinical isolates from chronic wounds of various etiology (later referred to as P4, P20, P30E, P34, P44E, P68E, P92E, P856) and two reference ones (marked ATCC 15442, ATCC 9027, American Type and Culture Collection). The tested strains were part of the Strain and Line Collection of the Platform for Unique Models Application, Department of the Pharmaceutical Microbiology and Parasitology, Medical University of Wroclaw. The bioethical approval was granted with the following number: Bioethical Committee of Wroclaw Medical University, protocol # 8/2016.

### Essential oils

The antimicrobial and antibiofilm activity of a commercial Thyme Essential Oil, thyme chemotype (TEO, obtained from the leaves of *Tymus vulgaris* L., Instytut Aromaterapii), was evaluated. Moreover, the antiseptic Prontosan Wound Irrigation Solution (marked PHMB, polyhexanide, B. Braun Medical AG), composed of 0.1% polyhexamethylene biguanide, 0.1% undecylenamidopropyl betaine, and purified water, was used as a comparator substance.

### Assessment of EOs chemical composition using gas chromatography-mass spectrometry (GC– MS)

The research was conducted to assess the percentage content of the TEO constituents. Firstly, the oil was diluted 50 times using hexane, mixed, and submitted for analysis with an Agilent 7890B GC system coupled with 7000GC/TQ, equipped with a PAL RSI85 autosampler (Agilent Technologies) and an HP-5 MS column (30 m × 0.25 mm × 0.25 μm). Helium was used as a carrier gas with a total flow of 1 mL/min. The ratio of injection splitting was 1:100. The initial temperature of the analysis was 50 °C maintained for 1 min. Next, the temperature was set to reach 170 °C (4 °C/min) and then to 280 °C (10 °C/min), which was kept for 2 min. The MS detector was set as follows: temperature of transfer line, source, and quadrupole – 320, 230, and 150 °C, respectively, and 70 eV voltage of ionization. Detection was performed in total scan mode at 30–400 m/z. The NIST 17.1 library and literature data were applied to compare the acquired mass spectra and the retention index (RI). Indexes of linear retention were evaluated under the conditions applied for the TEO analysis using a mixture of C8–C20 saturated alkanes. The relative abundance of each constituent was presented as a percentage content based on peak area normalization (according to the MassHunter Workstation Sofware version B.09.00). The analysis was carried out in triplicate.

### Determination of strains phylogentic relationship using ERIC-PCR

In the present study, the Enterobacterial Repetitive Intergenic Consensus – Polymerase Chain Reaction (ERIC-PCR) method was employed to amplify repetitive genetic sequences in the genomes of the analyzed microorganisms (31). The reaction was carried out using two primers – ERIC1 (5’ CACTTAGGGGTCCTCGAATGTA 3’) and ERIC2 (5’ AAGTAAGTAGTGGGGTGAGCG 3’) (Genomed). Genomic DNA isolated from the investigated strains served as the template, while nuclease-free water (A&A Biotechnology), a commercial PCR Mix Plus Green kit (A&A Technology), and distilled water were used to prepare the reaction mixture.

For each ERIC-PCR assay, 25 µl of reaction mixture was prepared, comprising 0.3 µl of primer ERIC1, 0.3 µl of primer ERIC2, 12.5 µl of PCR Mix Plus Green, 1.9 µl of nuclease-free water, and 10 µl of genomic DNA (final volume: 25 µl). Amplification was performed in a thermal cycler (TaKaRa PCR Thermal Cycler) for 30 cycles under the following conditions: initial denaturation at 95 °C for 10 min, denaturation at 90 °C for 30 s, primer annealing at 52 °C for 1 min, and elongation at 65 °C for 8 min, followed by a final elongation step at 65 °C for 16 min.

The resulting PCR products were analyzed by 2% agarose gel electrophoresis (Cleaver PowerPro 300). For this purpose, 2.8 g of agarose (Bio-Rad) was dissolved in a mixture of 112 ml distilled water and 28 ml 5× TBE buffer (BioShop), and the solution was brought to a boil. Subsequently, 6 µl Midori Green (NIPPON Genetics) dye was added, and the gel was poured into a tray once the solution had cooled slightly. Prior to loading, 4 µl of 6× loading buffer (A&A Biotechnology) was added to each 20 µl PCR product. DNA markers (5 µl of DNA Marker 3, A&A Technology, and 100 bp DNA Ladder H3 RTU, NIPPON Genetics) were applied to the first and last wells. Electrophoresis proceeded for approximately 60 min, and the gels were visualized using the Gel Doc XR+ Gel Documentation System (Bio-Rad). ERIC-PCR patterns were analyzed via visual assessment, and the dendrograms were generated with the UPGMA method using online software http://insilico.ehu.es/.

The strains were divided into three groups (marked *group1*, *group2*, *group3*) according to their genetic relationship. Subsequently, all parameters characterizing biofilm features and *P. aeruginosa* tolerance to the tested compounds were analyzed across the groups. For this purpose, mean values for each strain were calculated first.

### Culture conditions

The experiments were performed in a Tryptic Soy Broth medium (TSB, Biomaxima). The bacteria were incubated in the medium overnight at 37 °C, then suspended in saline for each test and 1.5× 10^8^ CFU/mL (Colony-Forming Unit) was adjusted with a densitometer DEN-1B (Biosan). Such suspensions were diluted x1000 in the medium and used in further analysis.

### Evaluation of biofilm biomass and biofilm metabolic activity using crystal violet and tetrazolium staining

The total biofilm mass and metabolic activity were assessed using crystal violet and tetrazolium staining, respectively, to characterize formed pseudomonal biofilms. Three independent experiments in four replicates were performed, and a mean was calculated for each strain. For both tests, biofilms were cultured as follows. The diluted pseudomonal suspensions prepared as described in the section Culture conditions were added to the wells of 96-well polystyrene plates (Wuxi Nest Biotechnology) at the volume of 200 µL and incubated under static conditions for 24 h at 37 °C. After the incubation, the medium was gently removed. For the biofilm biomass evaluation, the 20% (v/v) crystal violet (Chempur) water solution was poured into the wells at 200 µL, and the setting was incubated at room temperature for 10 min. Subsequently, the solution was pipetted out, the biofilm cells were washed with 200 µL of 0.9% (w/v) sodium chloride (Chempur), and the plates were kept at 37 °C for 10 min. Then, 200 µL of 30% (v/v) acetic acid (Chempur) water solution was added, and the plates were subjected to shaking at 450 rpm (Mini-shaker PSU-2T, Biosan SIA) at room temperature until the crystals dissolved. Finally, the absorbance of the solution was measured at 550 nm using a spectrophotometer (MultiScan Go, Thermo Fischer Scientific). For the biofilm metabolic activity assessment, the formed biofilms were stained with 200 µL of 0.1% (w/v) tetrazolium chloride solution (2,3,5-triphenyl-tetrazolium chloride) in TSB medium for 2 h. Subsequently, the solution was removed, and the stained biofilm cells were dried for 10 min at 37 °C. To dissolve formazan crystals, solution of methanol (Chempur) and acetic acid (9:1 ratio) was poured into the wells (200 µL), and the plates were shaken for 30 min at room temperature at 450 rpm. Next, the absorbance of the colour solution was measured with the spectrophotometer at 490 nm.

## Assessment of the antimicrobial activity

### Modified Disk Diffusion method

To apply the tested compounds for the experiment, biocellulose discs (BC) 15 mm in diameter were prepared by culturing a *Komagataeibacter xylinus* ATCC 53524 strain in the Herstin–Schramm (H-S) medium. The medium was comprised of 2% (w/v) glucose (Chempur), 0.05% (w/v) MgSO4·7H2O (POCH), 0.5% (w/v) bacto-peptone (VWR, Radnor), 0.115% (w/v) citric acid monohydric (POCH), 0.5% (w/v) yeast extract (VWR, Radnor), 0.27% (w/v) Na2HPO4 (POCH), and 1% (v/v) ethanol (Chempur). Firstly, the H-S medium was added to the wells of a 24-well plate (Wuxi Nest Biotechnology) at the volume of 1 mL, and inoculated with *K. xylinus*. The plates were incubated under static conditions for 7 days at 28 °C. The BC discs were transferred to a bottle and washed with 0.1 M NaOH (Chempur) at 80 °C to remove unadhered cells and cell debris, then washed with double-distilled water until neutral pH. Next, the BC discs were sterilized in an autoclave. For research purposes, the weight of BC discs ranged from 0.6 g to 0.7 g. The sterile BC discs were placed into fresh 24-well plates and 1 mL of undiluted TEO or PHMB or PBS (Phosphate-Buffered Saline, Sigma-Aldrich, positive control) and kept for 24 h at 4 °C to soak the discs. To evaluate the concentration of absorbed compounds, fresh six BC discs were dried for 24 h at 37 °C and then weighed. The average difference between wet and dry BC discs was approximately 0.67 g. Therefore, the concentration of TEO/ PHMB after the soaking was calculated as follows:

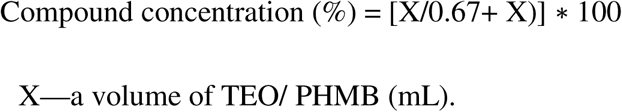

The disc diffusion test was performed using the undiluted 1.5× 10^8^ CFU/mL pseudomonal suspensions in saline prepared as described in the section Culture conditions. The bacteria were seeded onto TSA (Tryptic Soy Agar, Biomaxima, 90 mm diameter, 14.2 mm height, 5 mm agar thickness) Petri dishes (Noex). Next, the soaked BC discs were put onto the agar, and the plates were incubated for 24 h/ 37 °C. After the incubation, zones of bacterial growth inhibition were measured (in mm) with a ruler. When the measured zone was irregular, a shorter diameter was chosen. One independent experiment in three repetitions was performed.

### Microdilution method

The tested compounds’ Minimal Inhibitory Concentration (MIC) was evaluated in 96-well plates. TEO was applied as an emulsion in the medium and Tween 20 (Zielony Klub). Firstly, the stock emulsion was prepared by combining TEO with Tween 20 and vigorous vortexing. Next, the medium was added, and the entire mixture was vortexed. T-EO constituted 40% (v/v), and Tween 20 constituted 2% (v/v) of the stock emulsion volume. Subsequently, the stock emulsion was geometrically diluted in falcon tubes with TSB, and 100 µL of each concentration was added to a separate well. In the case of PHMB, the geometrical dilutions were prepared in the medium, and 100 µL was also added to each well. Subsequently, the diluted pseudomonal suspensions prepared as described in the section Culture conditions were added at the volume of 100 µL to the wells with the tested compounds. Therefore, the actual concentration applied to the bacterial cells ranged from 20 to 0.01% (v/v) for TEO and from 50 to 0.1% (v/v) for PHMB (concerning 100% as an undiluted PHMB solution). The plates were incubated for 24 h at 37 °C with continuous shaking at 450 rpm. Moreover, the following controls were prepared: controls of pseudomonal growth (bacteria in TSB), medium sterility (TSB only), control of Tween’s 20 antipseudomonal activity (Tween 20 at geometric concentrations ranged from 1 to 0.002% (v/v) in TSB). Once the incubation was finished, 1% (w/v) tetrazolium chloride solution in the medium was added to each well at 20 µL, and all plates were returned to the incubator for 2 h. The MIC value was found in the first well, where no formazan red colour was detected. The experiment was performed once with six replicates. In the case of differentiated MIC between the replicates, the higher value was chosen if it repeated at least three times. The MIC values were presented as percentage values, concerning 100% as undiluted compounds.

### Assessment of antibiofilm activity and number of biofilm-forming cells using quantitative culturing

The antibiofilm activity of the studied compounds was assessed in 48-well plates (Wuxi Nest Biotechnology). The compounds were applied at the highest possible concentrations. TEO was applied at the concentration of 5% (v/v) (as an emulsion in TSB with 2% (v/v) Tween 20), and PHMB was used as 50% (v/v) solution in TSB (concerning 100% as undiluted PHMB solution). At higher TEO concentrations, fragile pseudomonal biofilm was destroyed during TEO removal from above the biofilm structure due to the high density of TEO emulsion. In the first step, the diluted pseudomonal suspensions prepared as described in the section Culture conditions were added to the wells at the volume of 500 µL and incubated under static conditions for 24 h at 37 °C. Subsequently, the medium was removed from the wells, and 500 µL of the TEO, PHMB, or TSB was added. Moreover, the influence of 2% (v/v) Tween 20 on the biofilm of the P4 strain was examined. After the incubation, quantitative culturing was carried out. For this purpose, the medium was removed from the wells and biofilm cells were unattached from the surface by six-time pipetting and shaking at 600 rpm twice for 30 s with 2x 500 µL of 0.1% (w/v) saponin (VWR, Radnor) solution in water. Finally, the samples were serially diluted in sodium chloride. Next, 10 µL bacterial spots were seeded onto TSA plates, and the plates were incubated for 24 h/ 37 °C. The bacterial colonies were counted, and each sample’s CFU/mL (Colony-Forming Unit) was calculated. The experiment was performed in one biological repetition with three technical repeats. The results of the growth control were additionally used to evaluate the number of CFU/mL as a parameter for biofilm characterization. The volume of seeded bacteria was 10 µL; therefore, a minimal CFU/mL number detected according to the methodology was 100. However, a 100 CFU/mL value was assumed if no colony was observed in this dilution. For each sample treated with TEO or PHMB, the percentage reduction of biofilm cells was calculated in relation to the mean of the growth control. The results for the separate strains were previously changed to a log10 scale to present them on a graph.

### Visualization of live and dead biofilm-forming cells using fluorescent dyes and fluorescence microscopy

The antibiofilm effect of the examined compounds against *P. aeruginosa* was confirmed microscopically using a fluorescent microscope. For this research, P857 and ATCC 15442 strains were selected. The biofilm culturing and their treatment with TEO or PHMB were performed according to the protocol described in the Antibiofilm activity section. However, the experiment was conducted on 24-well plates, and the volume of applied reagents was 1 mL. After the biofilm’s incubation with the tested compounds, the staining with a fluorescent dye and microscopic visualization were performed as follows. Filmtracer™ LIVE/DEAD™ Biofilm Viability Kit (Thermo Fischer Scientific) prepared according to manufacturer’s instruction was applied as a dye to assess membrane integrity. A total of 10 µL of the reagent was added to each well of a 24-well plate for 15 min (RT, darkness). Next, the cells were washed once with 200 µL of double-distilled water and then fixed with 4% formaldehyde (Chempur) for 2hs. After fixation, formaldehyde was removed, and the biofilms were left to dry at room temperature in darkness. After drying, samples were analyzed using a fluorescence microscope Etaluma 600 (object lens with magnification4×). The whole surface (100%) of wells were recorded and combined (tilted) to form image panel showing changes in biofilm after exposure to T-EO. The clinically applied antiseptic referred to as the Prontosan containing 0.1% of active substance polihexanide was applied as experiment’s usability control setting.

### Statistical analysis

Calculations were performed using R (Version 4.4.3; Rstudio software (2025-02-13)). The distribution and variance homogeneity were calculated using Shapiro-Wilk and Levene test, respectively. The distribution was also inspected visually. Welch-ANOVA followed by post-hoc Games-Howell test was performed to compare differences between biofilm mass and biofilm metabolic activity across *Pseudomonas* strains. The correlation between biofilm mass and metabolic activity was fitted to the linear and exponential function, with the estimated R^2^ value. Kruskal-Wallis followed by Dunn’s test was performed to compare differences between inhibition zone values, MIC values and number of colony-forming units across *Pseudomonas* strains, as well as between conditions (TEO and PHMB) within each strain. The same test was performed to compare differences in investigated parameters across groups of genetically distinct strains of *Pseudomonas.* The results of statistical analyses with a significance level of p < 0.05 were considered significant.

## Results

### Evaluation of TEO chemical composition

In the first step of the research, the percentage content of the TEO’s components was assessed using GC-MS methodology (Fig. 1). Thymol prevailed over other components and constituted half of the percentage composition. Except for carvacrol (which exceeded the standard range at 0.15%) and carvacrol methyl ether (which was not present), all components aligned with the European Pharmacopoeia XI standards.

**Fig 1:**
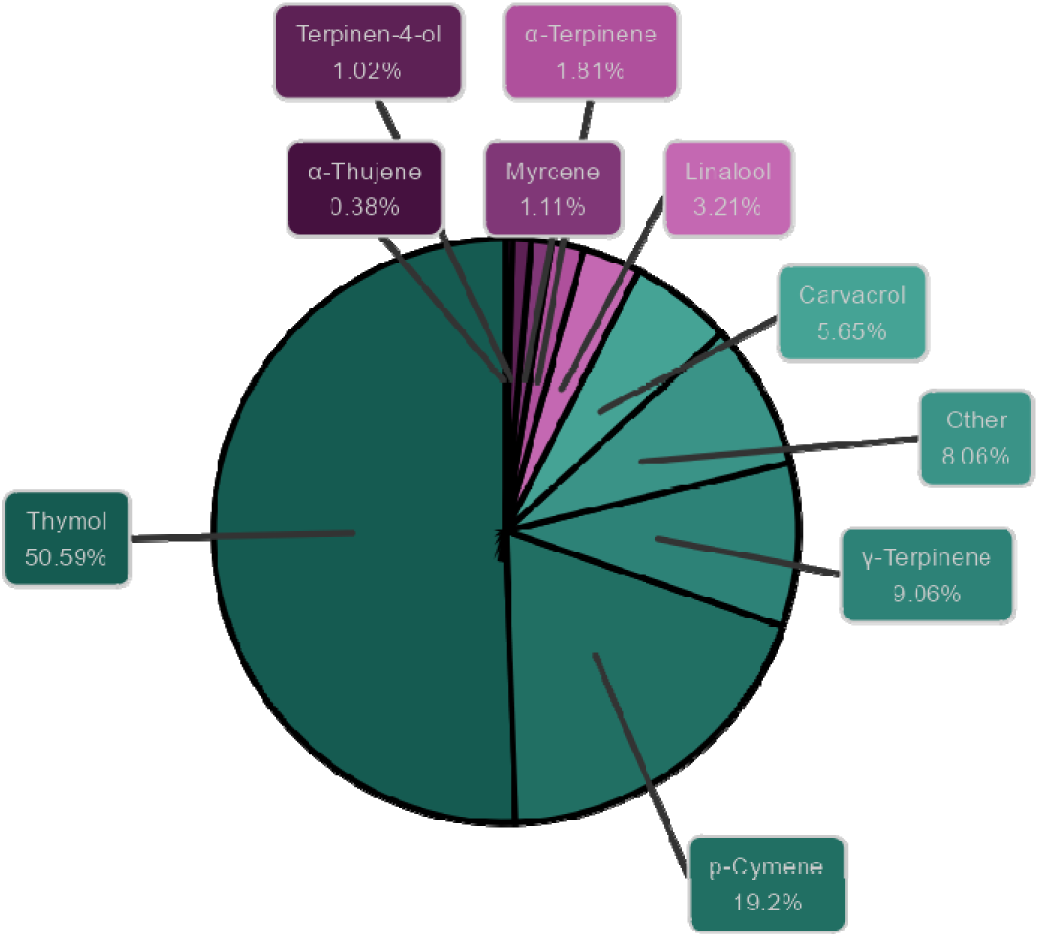
Composition of Thyme Essential Oil (TEO) by GC–MS. Relative percentage of constituents listed in the European Pharmacopoeia XI.

### Strains phylogentic relationship

To explore the genetic diversity among the studied *P. aeruginosa* strains, ERIC-PCR fingerprinting was performed, and dendrograms were generated using the UPGMA method with Dice similarity coefficients. A ≥90% similarity threshold was used to define clonally related strains, while clusters sharing <80% similarity were considered genetically distinct. Based on banding pattern analysis, three groups (*group1*, *group2*, and *group3*) were delineated (Fig. 2). Notably, strains P20 and P68 (*group3*) exhibited the lowest similarity to all other isolates, suggesting a divergent genetic background.

**Fig. 2:**
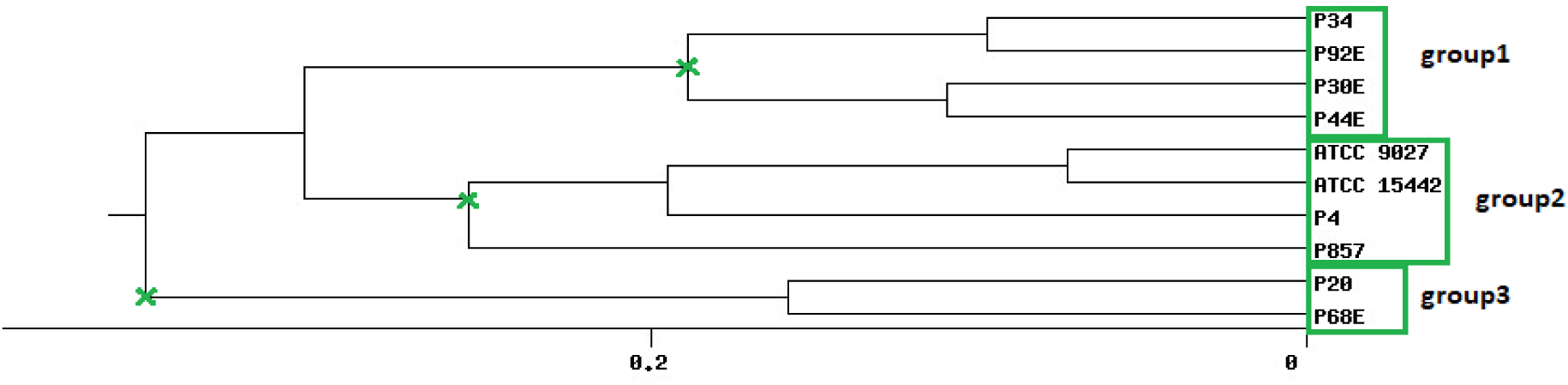
Phylogenetic clustering of *P. aeruginosa* strains based on ERIC-PCR fingerprinting. The dendrogram was constructed using UPGMA and Dice similarity index. A ≥90% similarity threshold was applied to define genetically related groups (green boxes and crosses). Group3 strains (P20, P68) formed a separate cluster with <80% similarity to the remaining isolates.

This stratification enabled the delineation of three distinct genetic groups, providing a framework for downstream analyses of intra-species variability in biofilm formation and antimicrobial susceptibility.

### Biofilm features

Phenotypic characterization of *P. aeruginosa* biofilms included quantification of biomass, metabolic activity, and viable cell counts (Fig. 3). The distribution of these features across individual strains is detailed in Fig. 4, Tables 1–2, and Supplementary Tables S1–S4. All strains formed biofilm under the applied conditions, though the extent varied markedly. Metabolic activity exhibited greater inter-strain variability than biomass, with statistically significant differences observed among 70% of strains (Welch’s ANOVA with Games-Howell post hoc, *p* < 0.05). Notably, strains of *group3*: P20 and P68 showed the lowest values for both biomass and metabolic activity, differing significantly from *group1* and *group2* strains. In contrast, strain P34 demonstrated exceptionally high metabolic activity. No correlation was observed between biomass and metabolic activity levels (Supplementary Fig. S1). Differences in CFU/mL across strains were less pronounced and not consistently statistically significant, likely due to limited sample size and variability, which reduced the sensitivity of non-parametric tests (Kruskal-Wallis with Dunn’s post hoc, Supplementary Table S4).

**Fig. 3:**
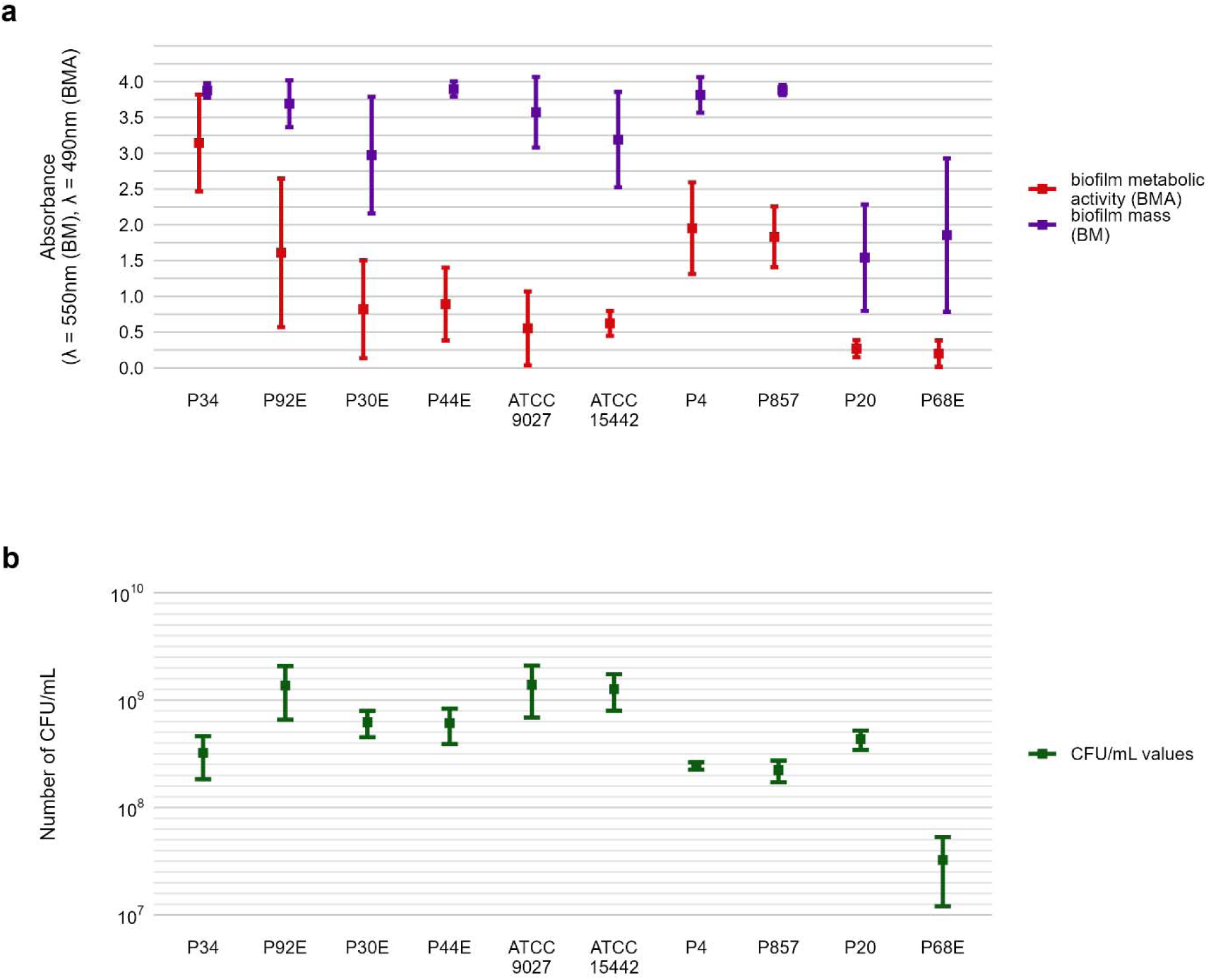
Characterization of *P. aeruginosa* strains (n= 10) biofilms. **a.** The average biofilm biomass and metabolic activity. **b.** The mean number of biofilm Colony-Forming Unit/mL (CFU/mL). The error lines represent the standard deviation.

**Fig. 4:**
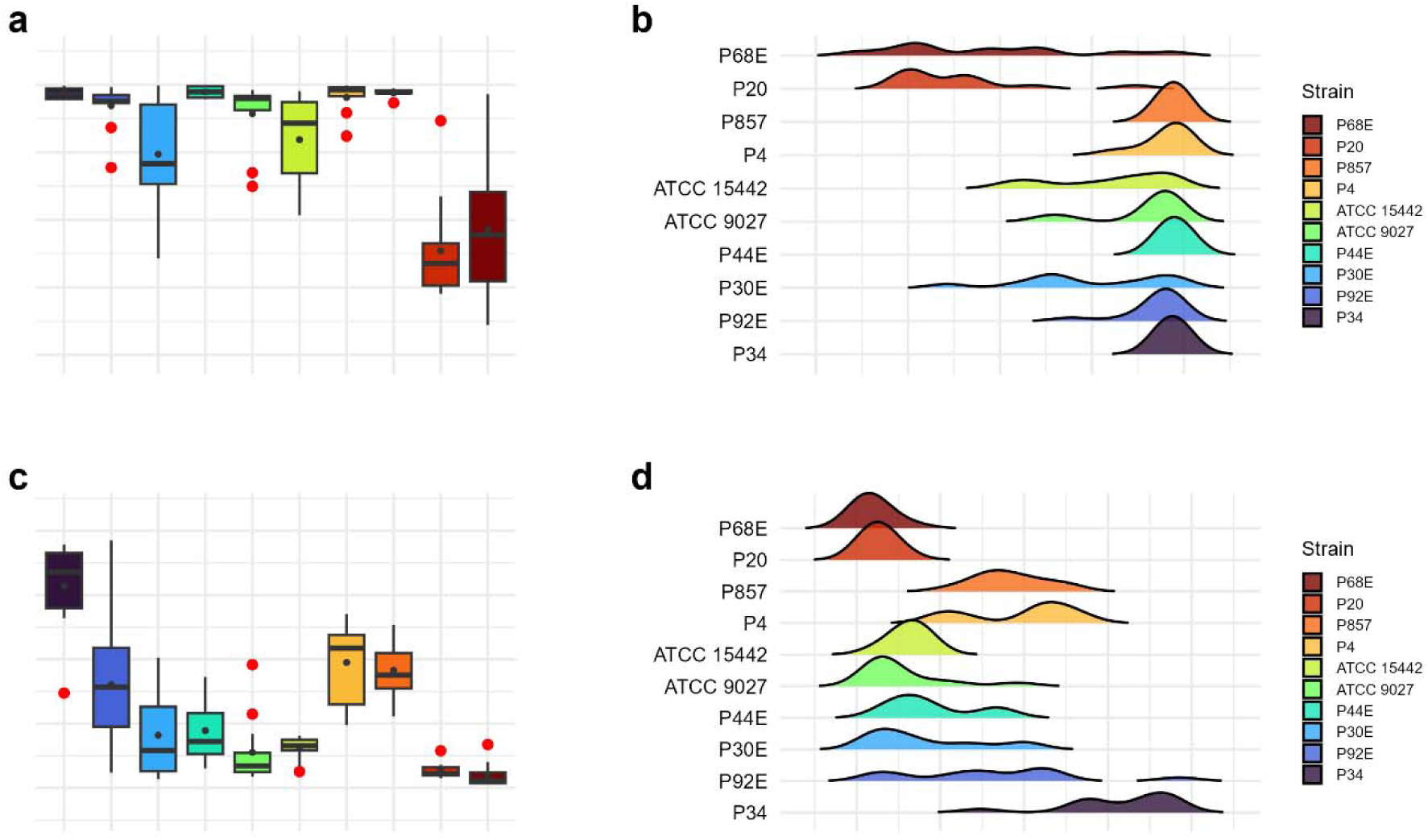
Visual representation of the data distribution for biofilm features of separate *P. aeruginosa* strains. **a-b.** Distribution of biofilm mass. **c-d.** Distribution of biofilm metabolic activity. The horizontal lines indicate a median value, the points indicate a mean value, the error lines indicate a standard deviation, the box indicates interquartile range.

**Table 1:** Summarized statistical differences in biofilm features between particular *P. aeruginosa* strains. **a.** Biological mass. **b.** Metabolic activity. Welch’s ANOVA, followed by the Games-Howell test, was performed. Values of p<0.05 were considered significant, p<=0.05 was marked with one asterisk, p<=0.01 was marked with two asterisks, p<=0.001 was marked with three asterisks, and p<=0.0001 was marked with four asterisks. Ns-no significant differences.

| **a** |  |  |  |  |  |  |  |  |  |  |
| --- | --- | --- | --- | --- | --- | --- | --- | --- | --- | --- |
| Strain | P34 | P92E | P30E | P44E | ATCC 9027 | ATCC 15442 | P4 | P857 | P20 | P68E |
| P34 | - | ns | ns | ns | ns | ns | ns | ns | **** | ** |
| P92E | ns | - | ns | ns | ns | ns | ns | ns | **** | ** |
| P30E | ns | ns | - | ns | ns | ns | ns | ns | ** | ns |
| P44E | ns | ns | ns | - | ns | ns | ns | ns | **** | ** |
| ATCC 9027 | ns | ns | ns | ns | - | ns | ns | ns | **** | ** |
| ATCC 15442 | ns | ns | ns | ns | ns | - | ns | ns | *** | ns |
| P4 | ns | ns | ns | ns | ns | ns | - | ns | **** | ** |
| P857 | ns | ns | ns | ns | ns | ns | ns | - | **** | ** |
| P20 | **** | **** | ** | **** | **** | *** | **** | **** | - | ns |
| P68E | ** | ** | ns | ** | ** | ns | ** | ** | ns | - |
| **b** |  |  |  |  |  |  |  |  |  |  |
| Strain | P34 | P92E | P30E | P44E | ATCC 9027 | ATCC 15442 | P4 | P857 | P20 | P68E |
| P34 | - | * | **** | **** | **** | **** | ** | *** | **** | **** |
| P92E | * | - | ns | ns | ns | ns | ns | ns | * | * |
| P30E | **** | ns | - | ns | ns | ns | * | ** | ns | ns |
| P44E | **** | ns | ns | - | ns | ns | ** | ** | * | * |
| ATCC 9027 | **** | ns | ns | ns | - | ns | *** | **** | ns | ns |
| ATCC 15442 | **** | ns | ns | ns | ns | - | *** | **** | *** | *** |
| P4 | ** | ns | * | ** | *** | *** | - | ns | **** | **** |
| P857 | *** | ns | ** | ** | **** | **** | ns | - | **** | **** |
| P20 | **** | * | ns | * | ns | *** | **** | **** | - | ns |
| P68E | **** | * | ns | * | ns | *** | **** | **** | ns | - |

**Table 2:** Summarized statistical differences in biofilm CFU/mL (Colony-Forming Unit) number between particular *P. aeruginosa* strains. Dunn’s test, followed by the Kruskal-Wallis test, was performed. Values of p<0.05 were considered significant, p<=0.05 was marked with one asterisk, p<=0.01 was marked with two asterisks. Ns-no significant differences.

**a**
| Strain | P34 | P92E | P30E | P44E | ATCC 9027 | ATCC 15442 | P4 | P857 | P20 | P68E |
| --- | --- | --- | --- | --- | --- | --- | --- | --- | --- | --- |
| P34 | - | * | ns | ns | * | * | ns | ns | ns | ns |
| P92E | * | - | ns | ns | ns | ns | * | * | ns | ** |
| P30E | ns | ns | - | ns | ns | ns | ns | ns | ns | * |
| P44E | ns | ns | ns | - | ns | ns | ns | ns | ns | * |
| ATCC 9027 | * | ns | ns | ns | - | ns | * | * | ns | ** |
| ATCC 15442 | * | ns | ns | ns | ns | - | * | * | ns | ** |
| P4 | ns | * | ns | ns | * | * | - | ns | ns | ns |
| P857 | ns | * | ns | ns | * | * | ns | - | ns | ns |
| P20 | ns | ns | ns | ns | ns | ns | ns | ns | - | ns |
| P68E | ns | ** | * | * | ** | ** | ns | ns | ns | - |

These findings highlight a pronounced intra-species variability in biofilm-forming capacity among *P. aeruginosa strains*, with metabolic activity emerging as a particularly discriminative trait. The genetically divergent *group3* strains (P20 and P68) consistently exhibited the weakest biofilm-forming phenotypes, aligning with their distinct ERIC-PCR profiles. Conversely, other strains displayed a wide spectrum of biomass and metabolic activity levels, independent of viable cell count. This heterogeneity underscores the limitations of single-strain or single-parameter models in evaluating biofilm-active compounds. Importantly, the observed decoupling of biomass, metabolic activity, and CFU counts supports the need for multi-parametric assessment strategies. This data provided a rationale for testing Thyme Essential Oil (TEO) across diverse genetic backgrounds and biofilm architectures, using *in vitro* models that capture both structural and functional aspects of microbial communities.

Next, we assessed the susceptibility of *P. aeruginosa* strains to TEO and PHMB using two complementary approaches: a modified disk diffusion assay and a microdilution method (Figs. 5 and 6; Tables 3–4; Supplementary Tables S5–S6). In the disk diffusion setup, bacterial cellulose (BC) discs were soaked with undiluted compounds, delivering an estimated final concentration of 60%. Both methods revealed marked strain-specific variability in response to TEO. The MIC values spanned a 1000-fold range, and inhibition zone diameters ranged from 0 to 7.7 mm, reflecting substantial differences in planktonic tolerance. In general, susceptibility profiles were consistent across methods, with most TEO-tolerant strains in microdilution also displaying reduced inhibition zones, except for strain P857.

**Fig. 5:**
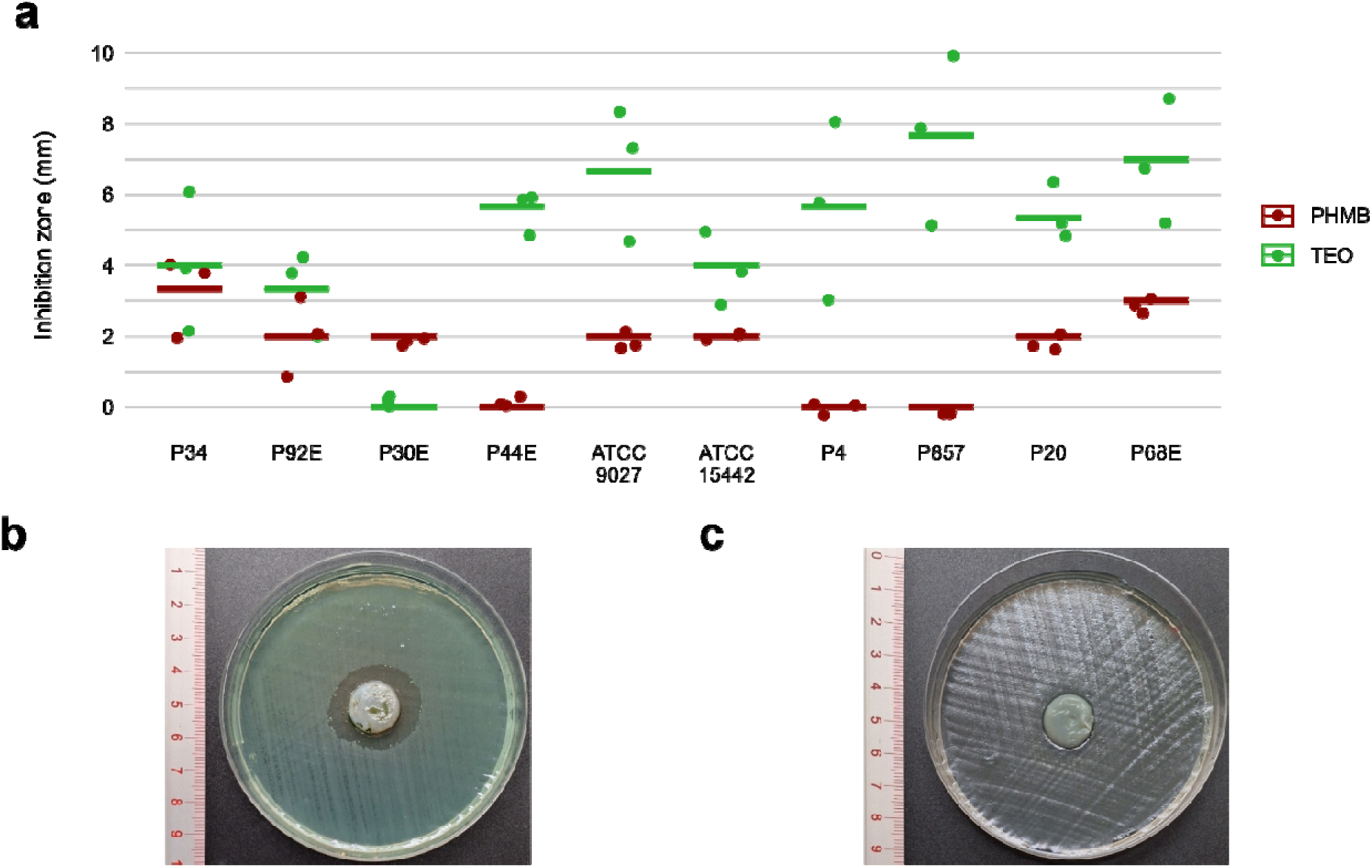
Antimicrobial activity of the tested compounds against *P. aeruginosa* strains assessed using modified disk diffusion method. **a.** Results diameters of growth inhibition zones (mm), (n=10). **b.** Representative zones of P857 strain growth inhibition after the treatment with Thyme Essential Oil (TEO). **c.** Representative zones of P857 strain growth inhibition after the treatment with polyhexanide (PHMB). The lines represent a mean value.

**Fig. 6:**
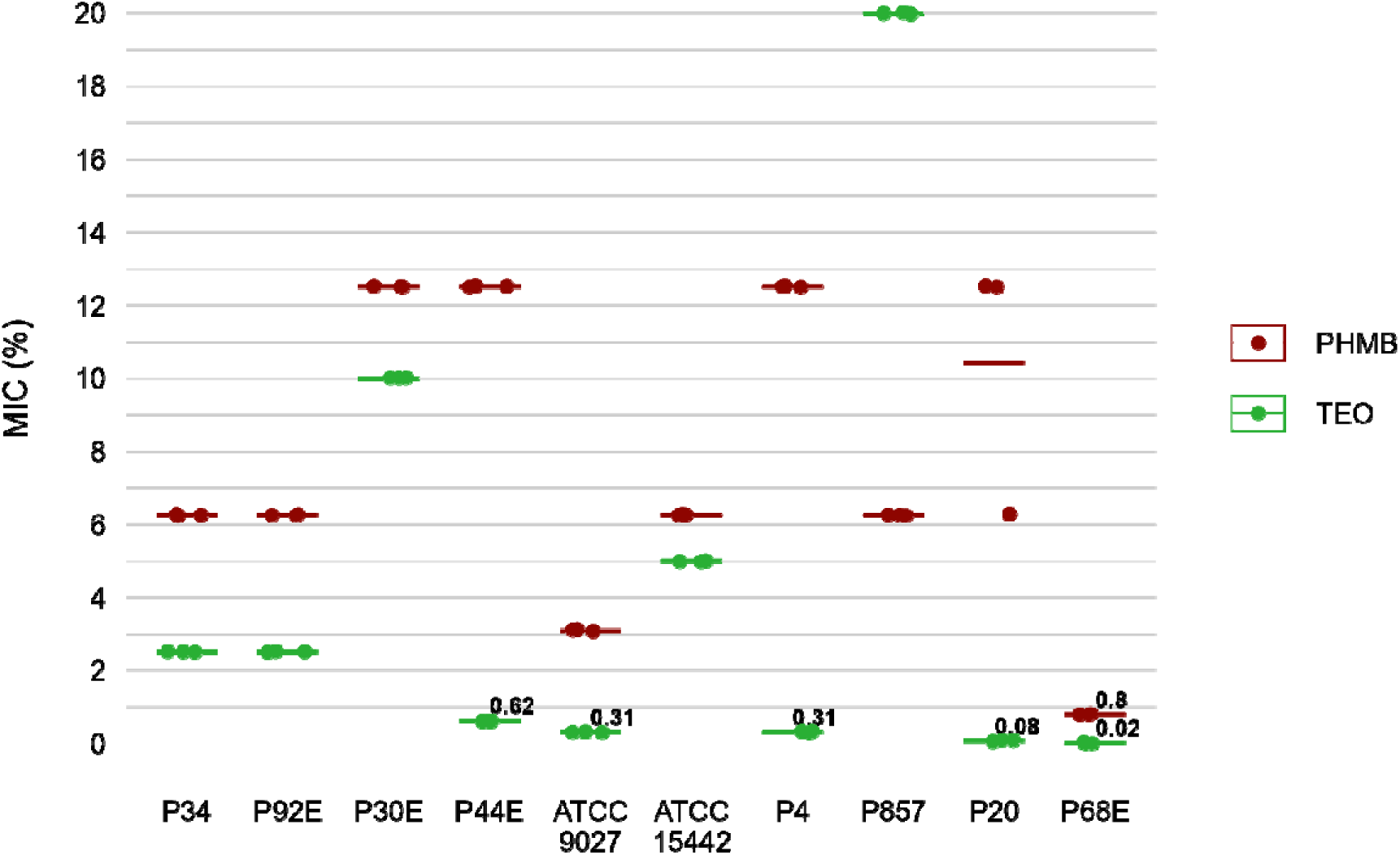
Antimicrobial activity of tested compounds against *P. aeruginosa* (n=10) strains assessed with a microdilution method. MIC-Minimal Inhibitory Concentration, TEO-Thyme Essential Oil, PHMB-polyhexanide. The lines represent a mean value.

**Table 3:**
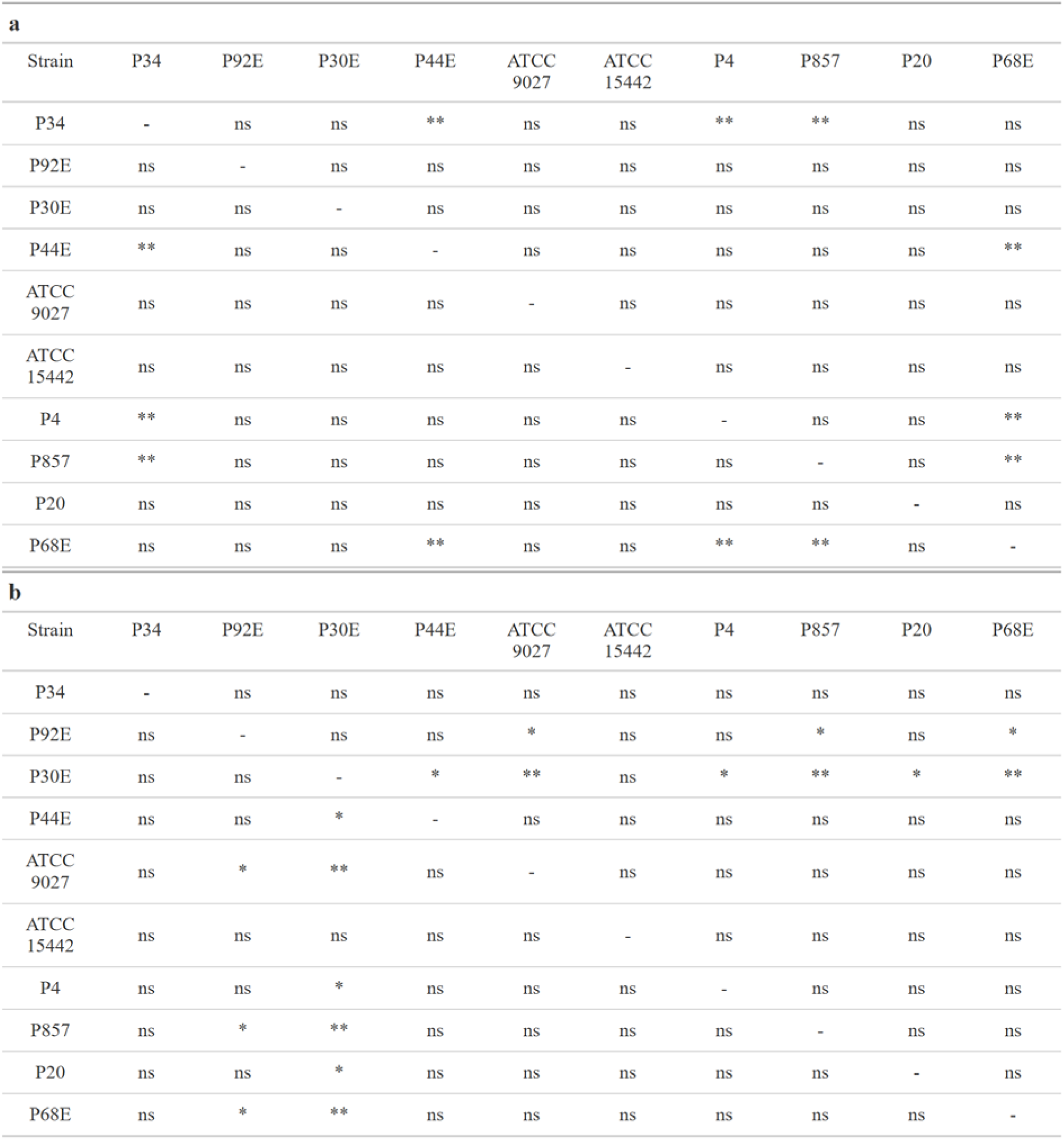
Summarized statistical differences in susceptibility to the tested compounds between particular *P. aeruginosa* strains expressed as growth inhibition zones (mm). **a.** Thyme Essential Oil (TEO). **b.** Polyhexanide (PHMB). Dunn’s test, followed by the Kruskal-Wallis test, was performed. Values of p<0.05 were considered significant, p<=0.05 was marked with one aster sk, and p<=0.01 was marked with two asterisk. Ns-no significant differences.

**Table 4:** Summarized statistical differences in susceptibility to the tested compounds between particular *P. aeruginosa* strains expressed as MIC (%) (v/v) values (Minimal Inhibitory Concentration). **a.** Thyme Essential Oil (TEO). **b.** Polyhexanide (PHMB). Dunn’s test, followed by the Kruskal-Wallis test, was performed. Values of p<0.05 were considered significant, p<=0.05 was marked with one asterisk, p<=0.01 was marked with two asterisks, and p<=0.001 was marked with three asterisks. Ns-no significant differences.

| **a** |  |  |  |  |  |  |  |  |  |  |
| --- | --- | --- | --- | --- | --- | --- | --- | --- | --- | --- |
| Strain | P34 | P92E | P30E | P44E | ATCC 9027 | ATCC 15442 | P4 | P857 | P20 | P68E |
| P34 | - | ns | ns | ns | ns | ns | ns | ns | ns | ns |
| P92E | ns | - | ns | ns | ns | ns | ns | ns | ns | ns |
| P30E | ns | ns | - | ns | ** | ns | ns | ns | ns | *** |
| P44E | ns | ns | ns | - | ** | ns | ns | ns | ns | *** |
| ATCC 9027 | ns | ns | ** | ** | - | ns | ** | ns | * | ns |
| ATCC 15442 | ns | ns | ns | ns | ns | - | ns | ns | ns | ns |
| P4 | ns | ns | ns | ns | ** | ns | - | ns | ns | *** |
| P857 | ns | ns | ns | ns | ns | ns | ns | - | ns | ns |
| P20 | ns | ns | ns | ns | * | ns | ns | ns | - | ** |
| P68E | ns | ns | *** | *** | ns | ns | *** | ns | ** | - |
| **b** |  |  |  |  |  |  |  |  |  |  |
| Strain | P34 | P92E | P30E | P44E | ATCC 9027 | ATCC 15442 | P4 | P857 | P20 | P68E |
| P34 | - | ns | ns | ns | ns | ns | ns | ns | ns | * |
| P92E | ns | - | ns | ns | ns | ns | ns | ns | ns | * |
| P30E | ns | ns | - | ns | * | ns | * | ns | ** | *** |
| P44E | ns | ns | ns | - | ns | ns | ns | * | ns | ns |
| ATCC 9027 | ns | ns | * | ns | - | ns | ns | ** | ns | ns |
| ATCC 15442 | ns | ns | ns | ns | ns | - | ns | ns | * | ** |
| P4 | ns | ns | * | ns | ns | ns | - | ** | ns | ns |
| P857 | ns | ns | ns | * | ** | ns | ** | - | *** | *** |
| P20 | ns | ns | ** | ns | ns | * | ns | *** | - | ns |
| P68E | * | * | *** | ns | ns | ** | ns | *** | ns | - |

Notably, strains P20 and P68—previously shown to form the weakest biofilms—were among the most susceptible to TEO. By contrast, PHMB exhibited more uniform activity across the strain panel, with MIC values clustering between 6.3% and 12.5% (v/v) in 80% of isolates, and inhibition zones ranging narrowly between 2.0 and 3.3 mm in 70% of cases. Overall, TEO showed superior antipseudomonal activity compared to PHMB in 90% of strains, with statistically significant differences observed in both assays (p < 0.05, Pairwise Dunn’s Test). Interestingly, some strains (e.g., P4, P44) demonstrated high susceptibility to TEO while showing relative resistance to PHMB, highlighting divergent mechanisms of action and potential strain-specific vulnerabilities.

In the subsequent step of the experiment, the antibiofilm activity of tested compounds was evaluated using quantitative culturing (Fig. 7, Table 5, Supplementary Tables S7 and S8). The antibiofilm effect of PHMB was significantly (p<0.05, Pairwise Dunn’s Test) higher than the one displayed by TEO against six of ten strains (Supplementary Table S11). However, higher variability in strain’s tolerance to PHMB was observed (Supplementary Table S8). The mean log reduction in biofilm cells after the treatment with PHMB ranged from approx. 2 to 7. In the case of TEO, the log reduction between 2 to 3.3 was determined for all strains except for P68.

**Fig. 7:**
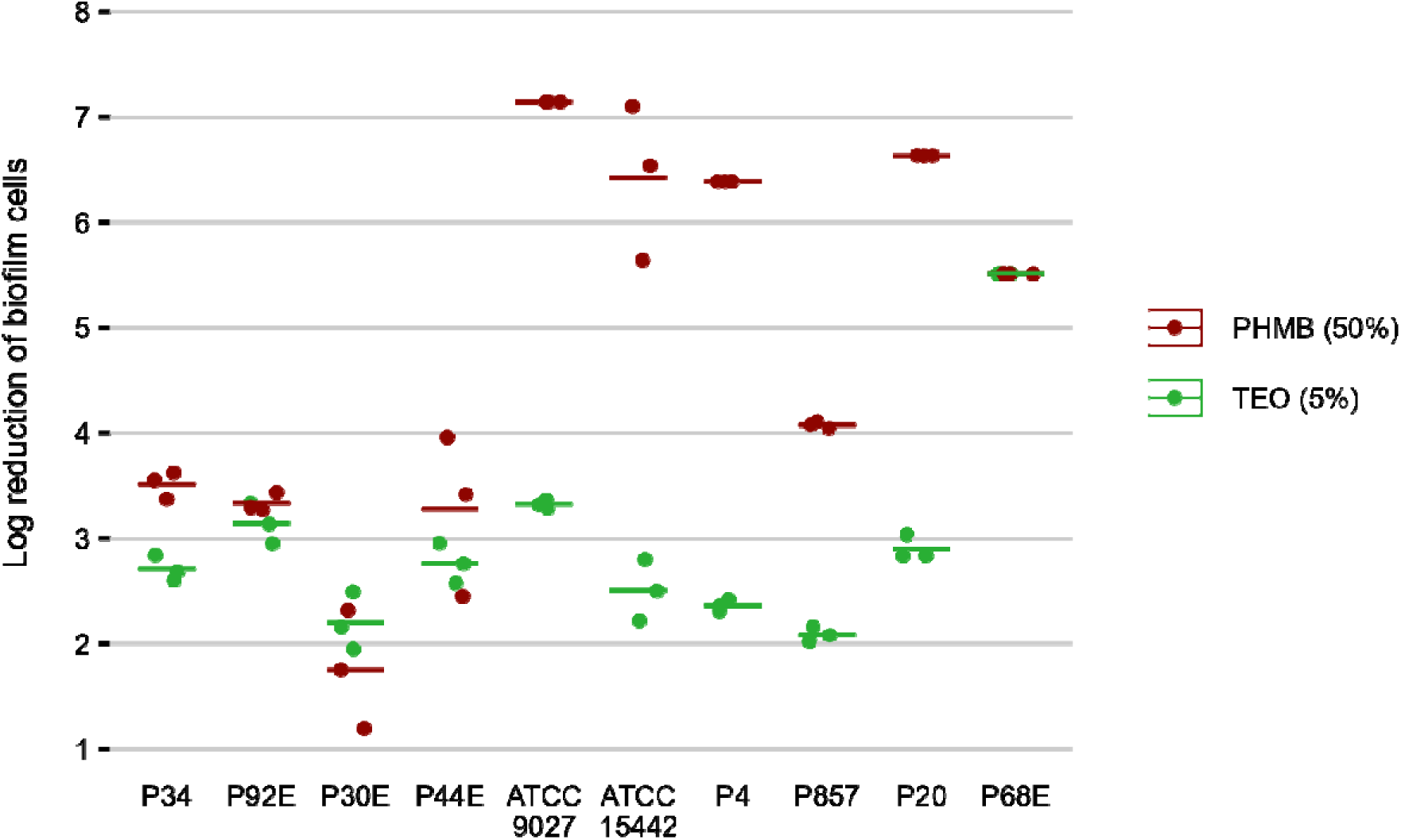
Antibiofilm activity of tested compounds against *P. aeruginosa* strains (n=10) expressed as a reduction (log10) of biofilm cells. TEO-Thyme Essential Oil, PHMB-polyhexanide. The lines represent a mean value.

**Table 5:**
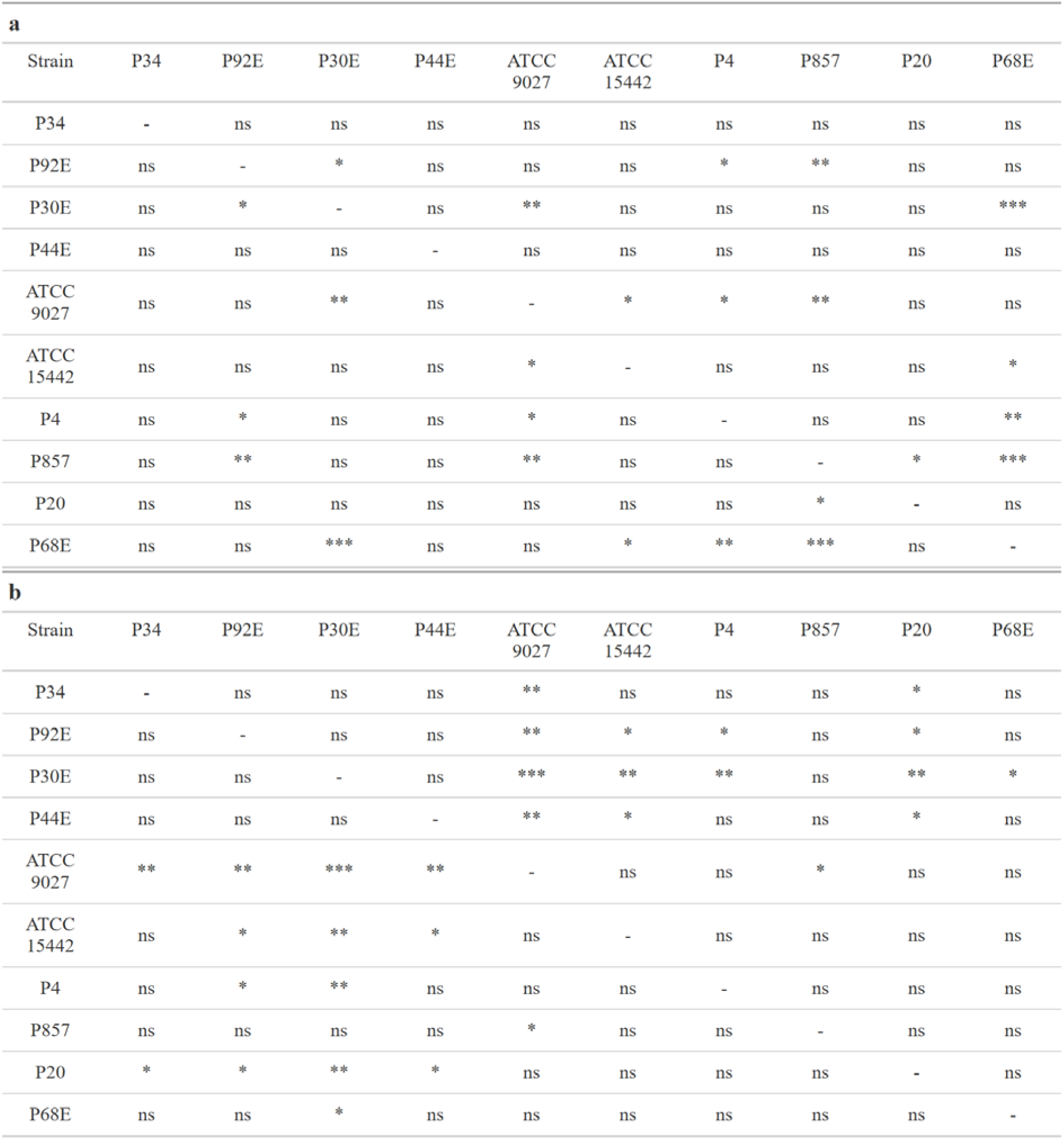
Summarized statistical differences in susceptibility to the tested compounds between particular *P. aeruginosa* strains expressed as biofilm reduction (%). **a.** Thyme Essential Oil (TEO). **b.** Polyhexanide (PHMB). Dunn’s test, followed by the Kruskal-Wallis test, was performed. Values of p<0.05 were considered significant, p<=0.05 was marked with one asterisk, p<=0.01 was marked with two asterisk, and p<=0.001 was marked with three asterisks. Ns-no significant differences.

| **a** |  |  |  |  |  |  |  |  |  |  |
| --- | --- | --- | --- | --- | --- | --- | --- | --- | --- | --- |
| Strain | P34 | P92E | P30E | P44E | ATCC 9027 | ATCC 15442 | P4 | P857 | P20 | P68E |
| P34 | - | ns | ns | ns | ns | ns | ns | ns | ns | ns |
| P92E | ns | - | * | ns | ns | ns | * | ** | ns | ns |
| P30E | ns | * | - | ns | ** | ns | ns | ns | ns | *** |
| P44E | ns | ns | ns | - | ns | ns | ns | ns | ns | ns |
| ATCC 9027 | ns | ns | ** | ns | - | * | * | ** | ns | ns |
| ATCC 15442 | ns | ns | ns | ns | * | - | ns | ns | ns | * |
| P4 | ns | * | ns | ns | * | ns | - | ns | ns | ** |
| P857 | ns | ** | ns | ns | ** | ns | ns | - | * | *** |
| P20 | ns | ns | ns | ns | ns | ns | ns | * | - | ns |
| P68E | ns | ns | *** | ns | ns | * | ** | *** | ns | - |
| **b** |  |  |  |  |  |  |  |  |  |  |
| Strain | P34 | P92E | P30E | P44E | ATCC 9027 | ATCC 15442 | P4 | P857 | P20 | P68E |
| P34 | - | ns | ns | ns | ** | ns | ns | ns | * | ns |
| P92E | ns | - | ns | ns | ** | * | * | ns | * | ns |
| P30E | ns | ns | - | ns | *** | ** | ** | ns | ** | * |
| P44E | ns | ns | ns | - | ** | * | ns | ns | * | ns |
| ATCC 9027 | ** | ** | *** | ** | - | ns | ns | * | ns | ns |
| ATCC 15442 | ns | * | ** | * | ns | - | ns | ns | ns | ns |
| P4 | ns | * | ** | ns | ns | ns | - | ns | ns | ns |
| P857 | ns | ns | ns | ns | * | ns | ns | - | ns | ns |
| P20 | * | * | ** | * | ns | ns | ns | ns | - | ns |
| P68E | ns | ns | * | ns | ns | ns | ns | ns | ns | - |

The activity of TEO and PHMB against *P. aeruginosa* ATCC 15442 and P857 biofilms was confirmed using also fluorescence microscope (Fig. 8). A higher number of bacterial cells with altered/damaged cell walls (red/orange color) was observed after the biofilms were exposed to TEO than PHMB.

**Fig. 8:**
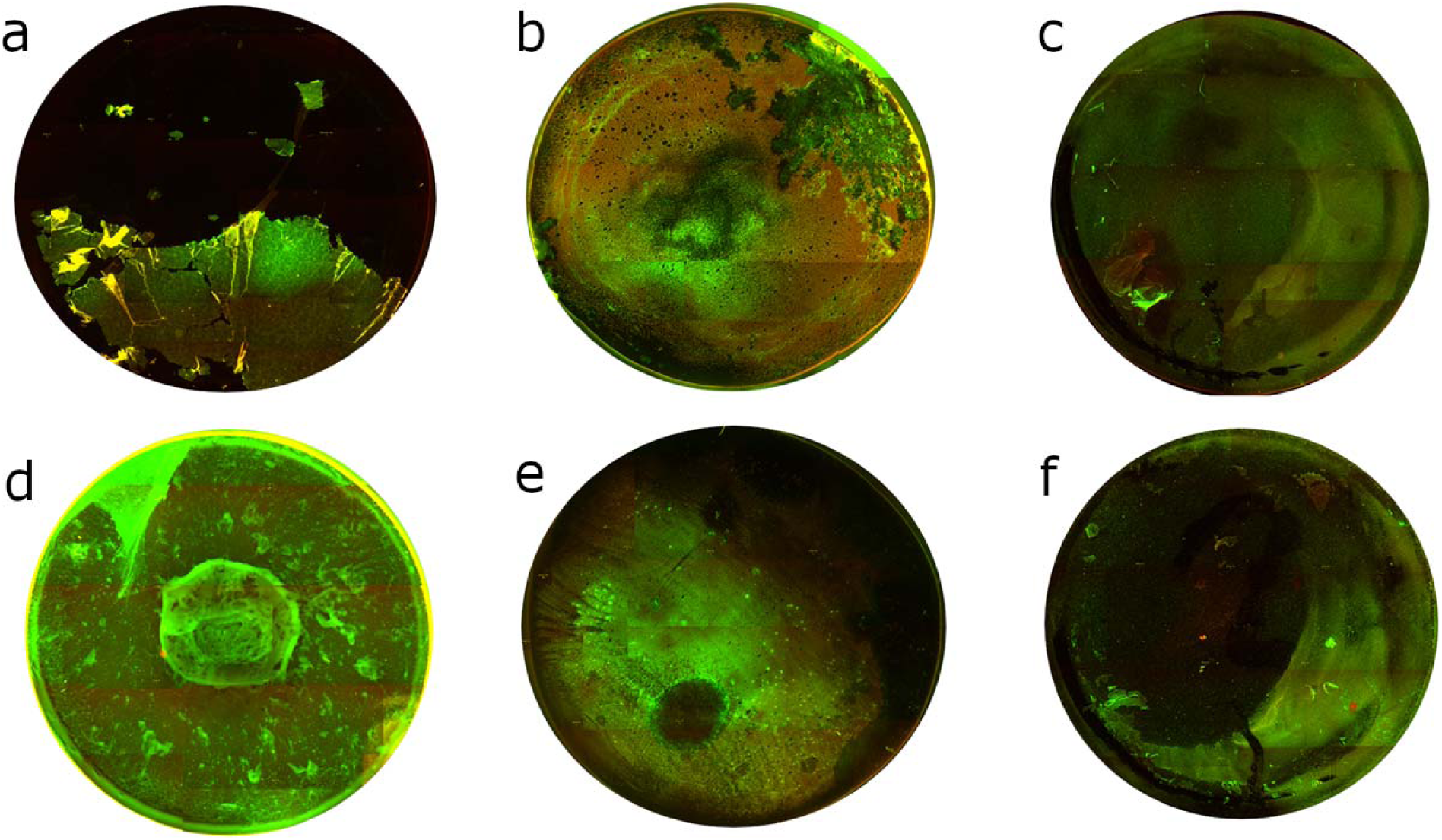
Microscopic visualizations of the *P. aeruginosa* biofilm (covering the surface of a 24-well plate) stained with a LIVE/ DEAD dye. **a.** ATCC 15442 strain’s untreated biofilm. **b.** ATCC 15442 strain’s biofilm treated with Thyme Essential Oil. **c.** ATCC 15442 strain’s biofilm treated with polyhexanide. **d.** P857 strain’s untreated biofilm. **e.** P857 strain’s biofilm treated with Thyme Essential Oil. **f.** P857 strain’s biofilm treated with polyhexanide. The red/orange color shows bacterial cells with altered/damaged cell walls; green color indicates unaltered cell walls. Fluorescence microscope Etaluma 600 (magnification 4×). The well diameter was 15 mm.

Subsequently, all parameters characterizing biofilm features and *P. aeruginosa* tolerance to the tested compounds were analyzed across the groups (Figs. 9-12, Supplementary Tables S11-S15). The biofilm mass and metabolic activity of *group3* were significantly lower in comparison to *group1* and *group2* (p<0.05, Dunn’s test, Supplementary Table S14). No statistically significant differences in biofilm-forming cell numbers between groups were observed. However, the parameter was at the lowest level in *group3*.

**Fig. 9:**
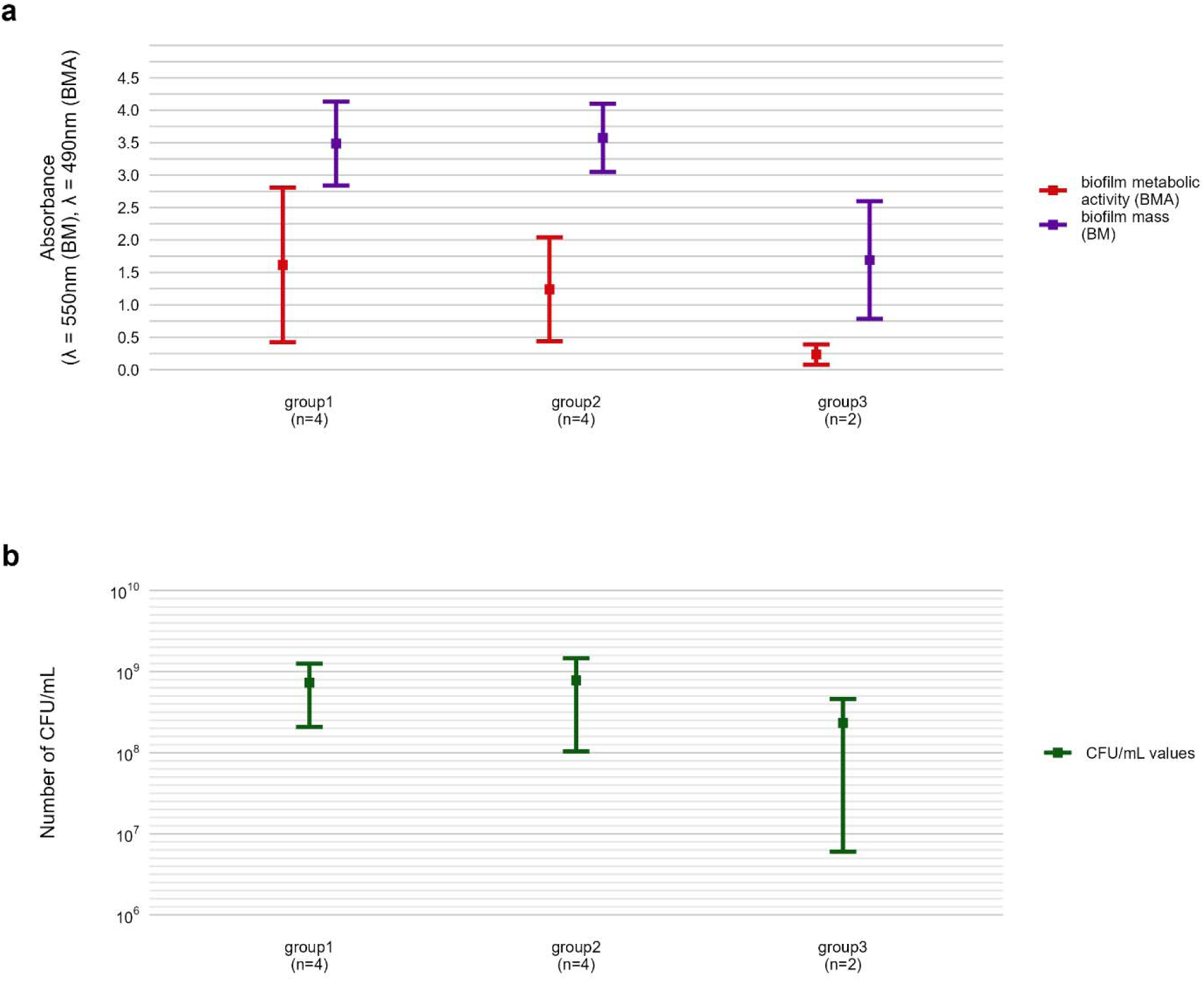
Comparison of biofilm features between genetically distinct groups of *P. aeruginosa* strains. **a.** The mean biofilm biomass and metabolic activity. **b.** The mean number of biofilm CFU/mL (Colony-Forming Unit). The error lines represent the standard deviation.

**Fig. 10:**
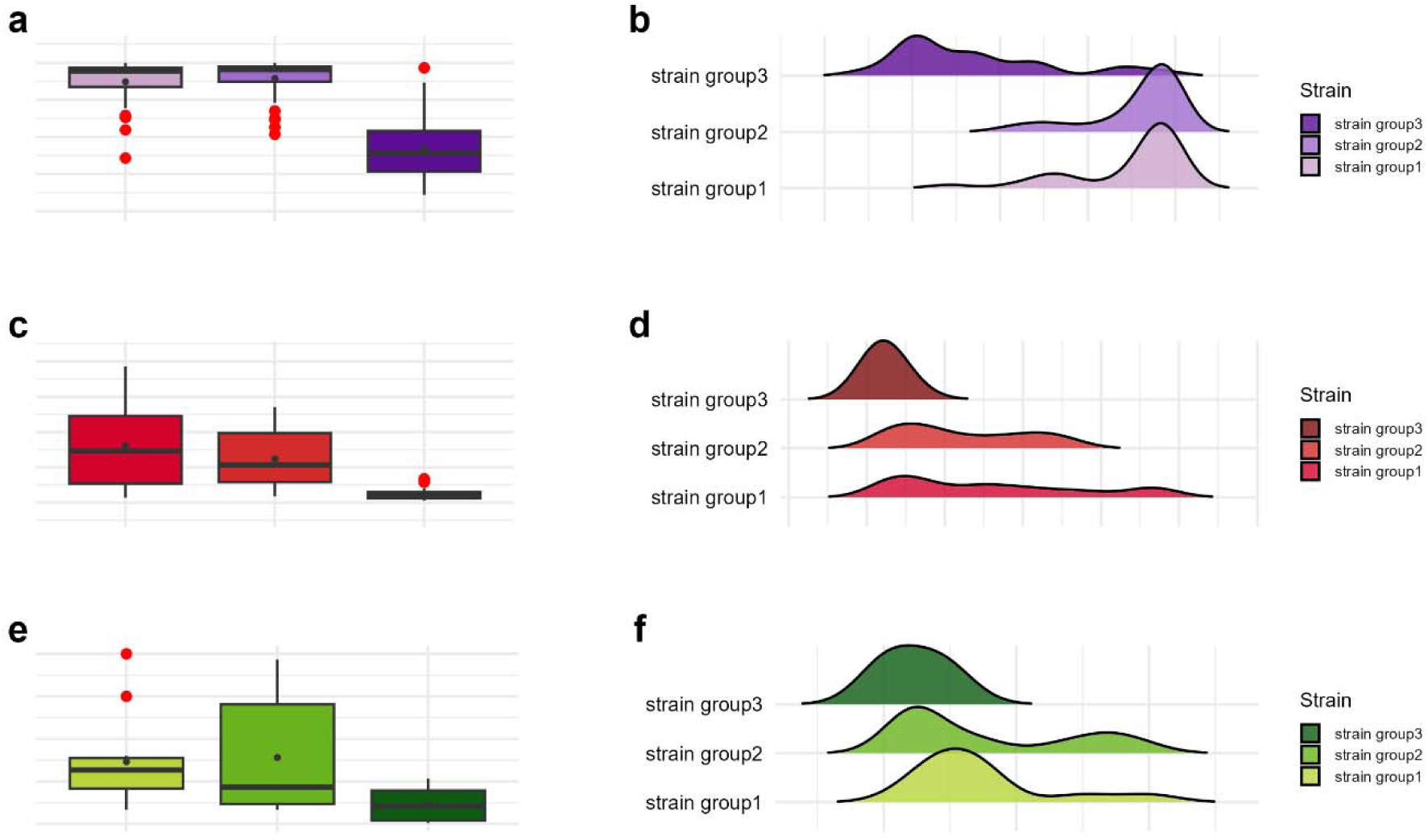
Visual representation of the data distribution for biofilm features across genetically distinct groups of *P. aeruginosa* strains. **a-b**. Distribution of biofilm mass. **c-d.** Distribution of biofilm metabolic activity. **e-f.** Distribution of biofilm Colony-Forming Unit number (CFU/mL). The horizontal lines indicate a median value, the points indicate a mean value, the error lines indicate a standard deviation, the box indicates interquartile range.

**Fig 11:**
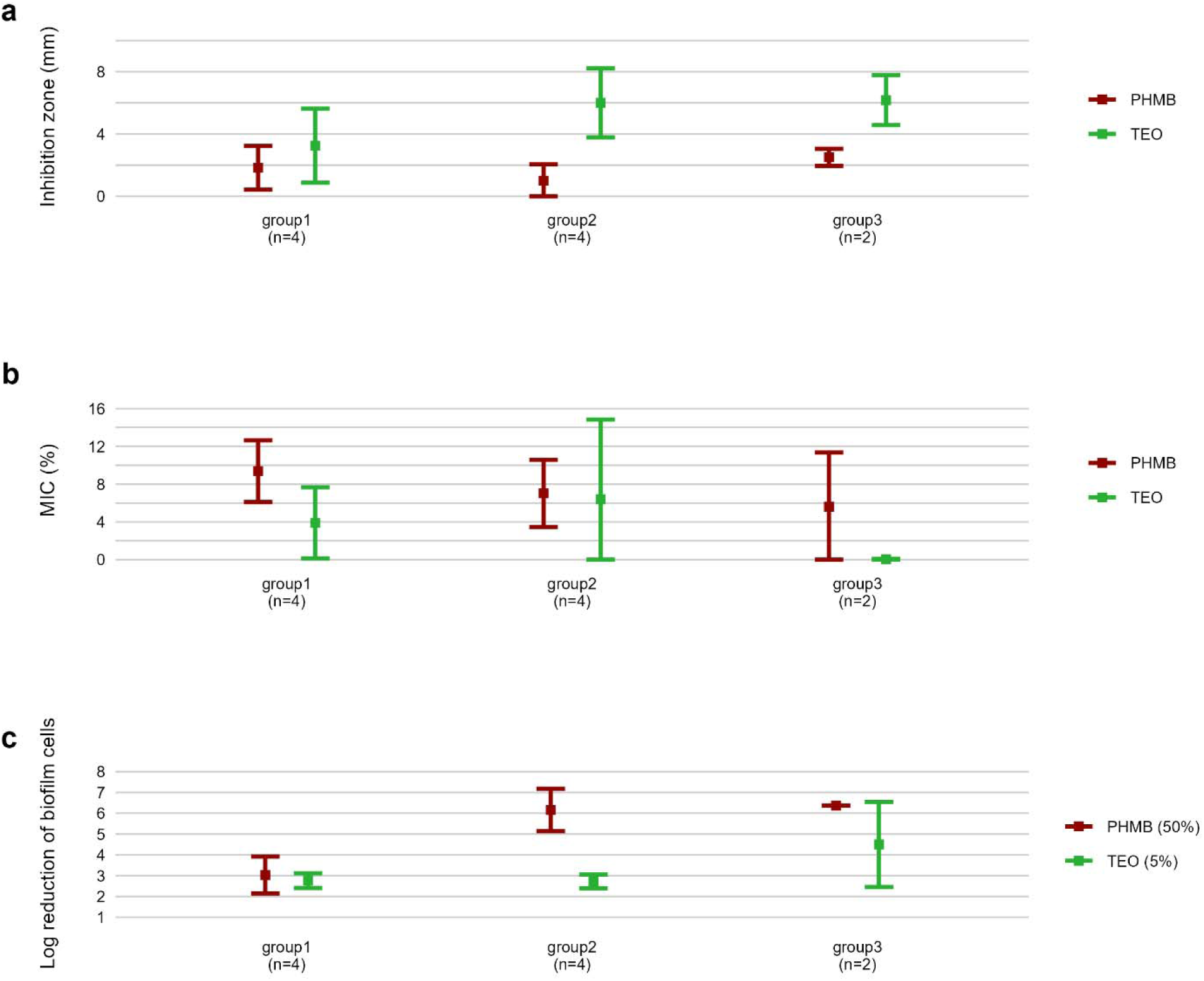
Comparison of antimicrobial and antibiofilm activity of tested compounds against genetically distinct groups of *P. aeruginosa* strains. **a**. The mean diameters of growth inhibition zones (mm). **b.** The mean values of Minimal Inhibitory Concentration (%, v/v)(MIC). **c.** The mean reduction (log10) of biofilm cells. TEO-Thyme Essential Oil, PHMB-polyhexanide. The error lines represent a standard deviation.

**Fig. 12:**
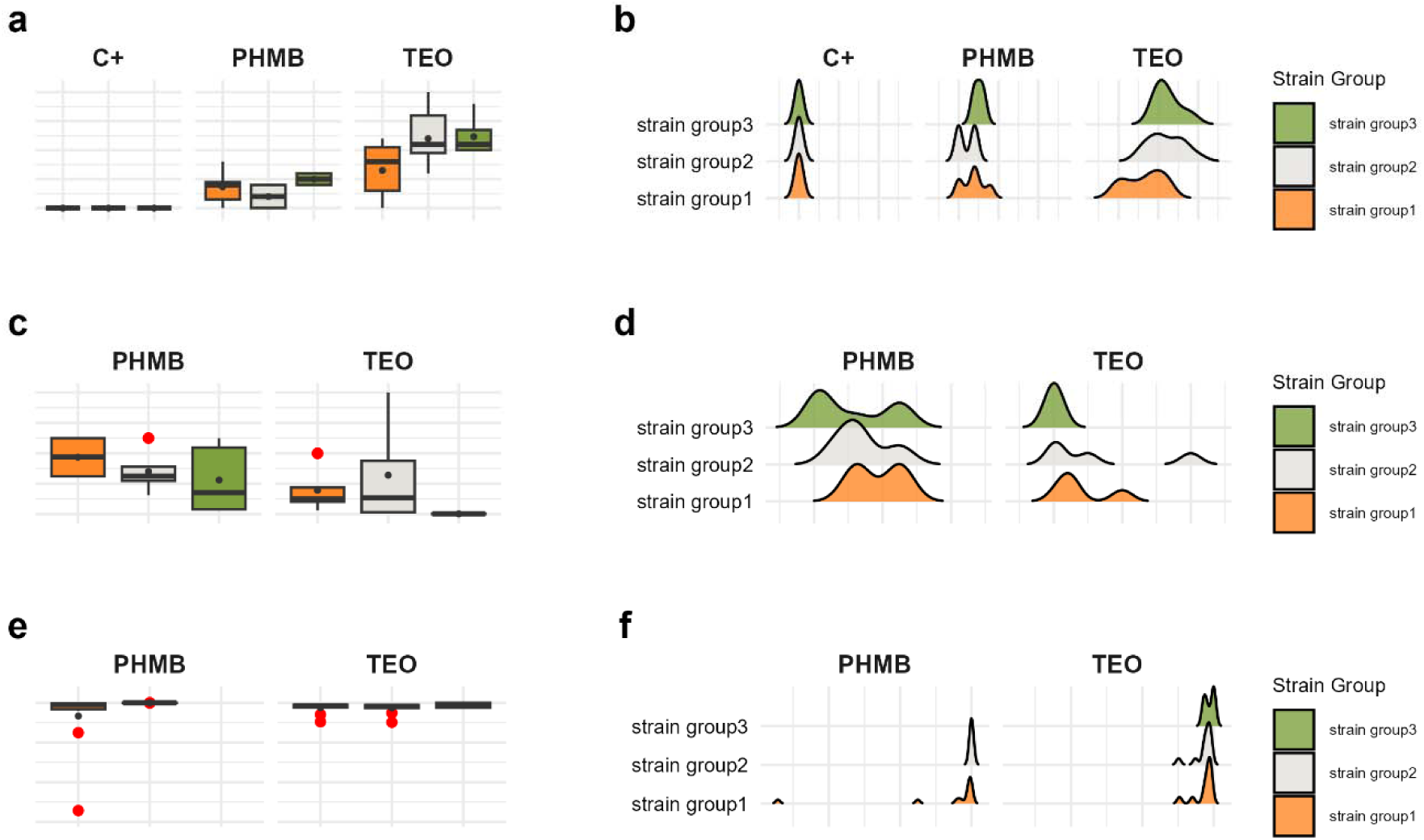
Visual representation of the data distribution for antimicrobial and antibiofilm activity of tested compounds against genetically distinct groups of *P. aeruginosa* strains. **a-b.** Distribution of growth inhibition zones values (mm). **c-d.** Distribution of Minimal Inhibitory Concentration (MIC) (%, v/v). **e-f**. Distribution of biofilm cells reduction (%). The horizontal lines indicate a median value, the spots indicate a mean value, the error lines indicate a standard deviation, the box indicate interquartile range. TEO-Thyme Essential Oil, PHMB-polyhexanide.

Strains in *group3* exhibited significantly lower tolerance (except for no significant difference with *group2* in the growth inhibition zones) to TEO than group1 and 2 in their planktonic forms and biofilms (p<0.05, Dunn’s test), Fig. 11 and Supplementary Table S15). However, in the case of *group3*, tolerance to PHMB was not as significantly different from other groups than in TEO.

The summarization of results is presented in heat-map (Fig. 13).

**Fig. 13:**
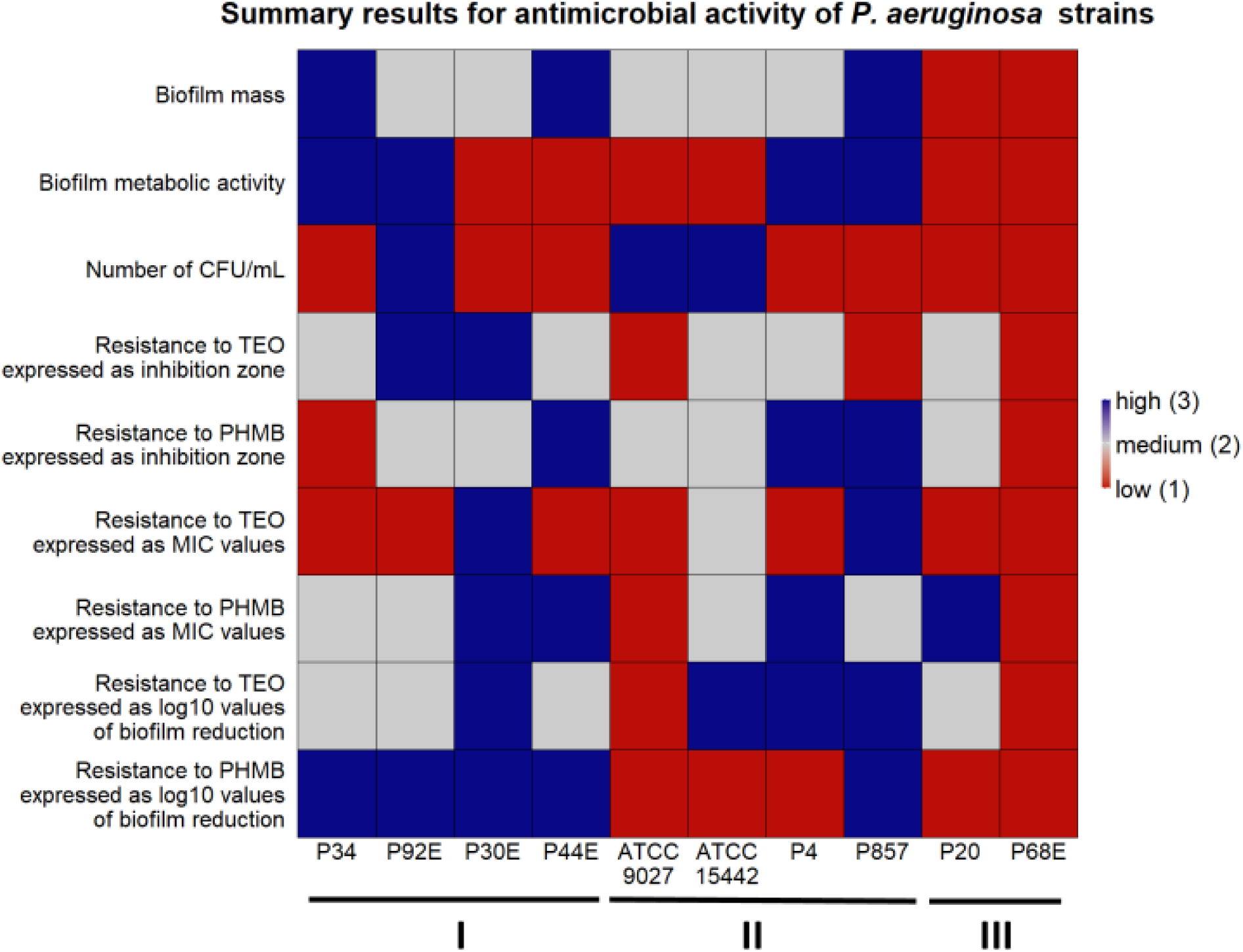
Summary of *P. aeruginosa* strains’ biofilm features and pseudomonal resistance to Thyme Essential Oil (TEO) or PHMB (polyhexanide) in genetically distinct groups. Heatmap was generated by setting a mean of all strains for each parameter ± 25% as high when the value was above the range (blue), low when the value was below the range (red) and medium when the value was equal to the range (grey). CFU/mL-Colony-Forming Unit/mL.

The integrative comparison of biofilm phenotypes and antimicrobial susceptibility across genetically defined groups revealed consistent trends supporting the biological relevance of ERIC-PCR-based clustering. Strains belonging to *group3* demonstrated overall attenuated biofilm-forming capacity, reflected in significantly lower biomass, metabolic activity, and viable cell counts compared to *group1* and *group2*. In antimicrobial assays, the modified disk diffusion (MDD) method identified 30% of strains as exhibiting low resistance to TEO, and 20% as showing low resistance to PHMB. Notably, half of the strains in groups 2 and 3 were classified as low TEO-resistant, whereas half of *group1* strains exhibited high resistance to this compound. Consistent with this, the microdilution method revealed that 75% of *group1* strains displayed low resistance to TEO. Across the entire panel, 7 out of 10 strains demonstrated low resistance to TEO, while only 2 strains exhibited low resistance to PHMB. When assessing biofilm-associated tolerance, a greater proportion of strains showed reduced susceptibility to PHMB than to TEO. Only 20% of isolates displayed low biofilm-level resistance to TEO. Remarkably, all *group1* strains were highly resistant to PHMB in the biofilm model, while 75% of *group2* strains showed the inverse pattern—high resistance to TEO and low resistance to PHMB. *Group3* strains exhibited uniformly low or intermediate resistance to both compounds. Individual strain-level profiles further emphasized this heterogeneity: P34, P44E, and P857, which formed dense biofilms, were markedly resistant to PHMB in the biofilm state; ATCC 9027 displayed low-to-moderate resistance to both agents across all models; strain 30E was consistently less susceptible to TEO regardless of the method; and P68E combined weak biofilm formation with low resistance to both compounds, making it the most responsive strain in the panel.

## Discussion

The rationale for this study arose from the growing recognition that antimicrobial efficacy cannot be reliably inferred from single-strain or single-method experiments—particularly when both the pathogen (*Pseudomonas aeruginosa*) and the antimicrobial agent (Thyme Essential Oil) exhibit inherent biological and chemical heterogeneity, respectively. Our findings confirmed extensive intra-species variability in biofilm-forming capacity and susceptibility to both Thyme Essential Oil (TEO) and chemically synthesized polyhexanide antiseptic (PHMB), affecting planktonic and biofilm-associated phenotypes alike. Importantly, this variability was not random but followed – to a certain extent - genetic stratification patterns, suggesting that distinct lineages may harbor intrinsic differences in tolerance mechanisms. Moreover, susceptibility profiles were strongly influenced by the assay system used, underscoring the importance of model selection in evaluating complex antimicrobials such as essential oils.

These observations resonate with a broader shift in the field toward antimicrobial precision and stewardship. For antibiotics, susceptibility testing is already governed by international standards, with clearly defined breakpoints and reference strains that enable reproducible, clinically meaningful assessments (32). In the case of antiseptics, the concept of “antiseptic stewardship” has recently gained traction, driven by reports of existing or emerging tolerance to compounds such as chlorhexidine, octenidine, and PHMB (33). Yet essential oils—despite their high potential in topical therapy—remain largely exempt from such critical scrutiny. This is particularly concerning given that they are often marketed and applied under the assumption of universal efficacy and low resistance potential (34,35).

Our data challenges this assumption. We demonstrate that even a pharmacopoeia-standardized essential oil, dominated by thymol, exhibits highly variable activity depending on the genetic background of the strain and the biological context of testing. In this light, we advocate for a new conceptual framework: essential oil stewardship. This would entail systematic evaluation of chemically defined oils against diverse and clinically relevant microbial panels, supported by multi-model testing strategies (36). Such an approach is urgently needed to move beyond anecdotal efficacy claims and establish a rational, evidence-based foundation for phytocompound deployment in infection control (37).

In this study, we controlled for chemical composition by using a pharmacopoeia-grade TEO batch characterized via GC–MS (**Fig. 1**), thereby eliminating compound heterogeneity as a confounder. This allowed us to isolate and examine three primary variables driving the observed differences in antimicrobial outcomes: (i) the genetic identity of the *P. aeruginosa strain*, (ii) the physiological state of the bacteria—planktonic versus biofilm-associated, and (iii) the specific *in vitro* model employed. These basic dimensions, yet they can be further developed, shaped both the magnitude and pattern of susceptibility, underscoring the need for multi-layered evaluation frameworks when assessing complex antimicrobial agents.

As mentioned, *P. aeruginosa* strains were stratified into three distinct genetic groups (**Fig. 2**); the phenotypic profiling of biofilm-forming capacity (**Fig. 3–4**) revealed marked inter-strain differences in both biomass and metabolic activity of all of them, despite identical growth conditions. While all ten *P. aeruginosa* strains formed detectable biofilms, the magnitude and functional characteristics varied significantly. Notably, metabolic activity showed a broader dynamic range than biomass, and no consistent correlation was observed between the two parameters—suggesting divergent regulatory pathways underpinning biofilm quantity and viability. *Group3* strains (P20, P68) exhibited consistently attenuated biofilm traits, while others (e.g., P34) showed disproportionately high metabolic output. Viable cell counts (**Tab. 2**), although less discriminatory, reinforced this heterogeneity. These divergent biofilm phenotypes may have critical implications for susceptibility to TEO, whose antimicrobial activity depends on both direct contact and vapor-phase diffusion (38,39). Strains producing high biofilm biomass—with abundant extracellular matrix—may experience reduced diffusion of TEO components, particularly hydrophobic monoterpenes like thymol, thereby conferring a physical barrier to penetration (40). However, if metabolic activity within the biofilm remains high, increased uptake of volatile or partially soluble fractions may paradoxically sensitize certain subpopulations (41). Conversely, low-biomass, low-activity biofilms, such as those formed by *group3* strains, likely lack such structural protection and may be more uniformly exposed to the active compounds—potentially explaining their higher susceptibility in subsequent assays (42). In turn, strains like P34, which produced metabolically hyperactive but structurally less complex biofilms, might remain vulnerable due to elevated uptake or metabolic engagement with the oil components. These nuanced relationships suggest that resistance to EOs is not solely a function of matrix density or cell number, but rather an emergent property of the interplay between biofilm architecture, physiological state, and compound delivery mode (**Fig. 5-7**, **Tab. 3-5**).

In turn, PHMB, obtained via chemical synthesis, is a cationic polymer that exerts its antimicrobial effect through strong electrostatic interactions with negatively charged bacterial surfaces, leading to membrane disruption, increased permeability, and, ultimately, intracellular interference—particularly with DNA (43). Notably, PHMB does not require active metabolism for uptake and retains efficacy irrespective of bacterial physiological state. This may explain why strains forming dense or metabolically quiescent biofilms (e.g., P44E, P857) remained susceptible to PHMB, despite showing reduced sensitivity to TEO. It is important to note, however, that in our study PHMB was applied as a commercial wound irrigation solution, also containing the amphiphilic surfactant undecylenamidopropyl betaine. This additive likely enhanced biofilm penetration by reducing surface tension and improving compound dispersion, which may have contributed to the overall uniformity of PHMB’s antibiofilm activity across strains (44). This observation offers a relevant translational insight: the inclusion of surfactants or other delivery-enhancing excipients may represent a promising strategy for improving the performance of EOs, not only by facilitating their penetration into biofilm structures but also by reducing the variability of outcomes. The mechanistic divergence between PHMB and TEO thus underscores not only their differential interaction with biofilm architectures but also the importance of formulation in optimizing and standardizing antimicrobial efficacy. Our findings emphasize that compound-specific delivery profiles must be carefully considered when interpreting biofilm susceptibility data, especially for complex, multicomponent agents such as EOs.

Our results also highlight critical methodological limitations associated with plate-based biofilm assays. Repeated washing steps, often necessary in classical protocols, can disrupt or incompletely remove loosely attached biomass, leading to an underestimation of actual biofilm architecture (45). This effect was clearly visualized in our fluorescence microscopy panel (**Fig. 8**), where peripheral and central regions of the same well displayed markedly different biofilm structures—even under uniform culture conditions. Such heterogeneity implies that selective imaging of “representative” areas, a common practice in low-throughput settings, may introduce substantial observational bias. In contrast, our approach—mapping entire wells and correlating imaging data with quantitative metrics—revealed spatial variation that would likely be missed by partial sampling. This well-to-well and intra-well structural variability adds yet another layer to the complex landscape of antimicrobial response diversity. In future work, we aim to further dissect this spatial dimension of biofilm behavior to refine interpretation of biofilm susceptibility results and reduce methodological bias.

The integrative heat map (**Fig. 9**) reflects the cumulative outcome of all prior phenotypic, susceptibility, and structural analyses—serving as a functional convergence point of the study. Patterns observed in biofilm mass, metabolic activity, and viable cell counts (**Figs 3–4**, **Tab. 2**) translate directly into biofilm-specific resistance profiles. Strains from *group3* exhibited a unique signature: high overall biofilm biomass, yet low cell density and metabolic activity, implying a disproportionately high extracellular matrix content. This matrix-dominant architecture likely hindered the diffusion of the hydrophobic components of TEO, resulting in moderate resistance, but paradoxically did not protect as effectively against PHMB—whose cationic, amphiphilic nature may enable adsorption and deeper matrix penetration (46,47). In contrast, *group1* strains—particularly P34 and P30E—demonstrated high resistance to TEO, correlating with elevated metabolic activity and denser, more viable biofilms, which may support active detoxification or efflux (48). The heat map also exposes strain-specific nuances within genetically related groups, as exemplified by ATCC 9027 or P857 (*group2*), which displayed an atypically high tolerance to PHMB. Collectively, the heat map confirms that biofilm-mediated tolerance is not determined by any single parameter but arises from the interplay of genotypic background, matrix-to-cell ratio, and biofilm vitality—further modulated by the molecular and formulation properties of the tested compounds.

In summary, our findings underscore the complexity of assessing antimicrobial efficacy against *P. aeruginosa*, a species marked by substantial intra-species heterogeneity in both planktonic and biofilm states. By integrating genotypic grouping, phenotypic profiling, and two complementary *in vitro* models, we demonstrate that resistance to antiseptic agents—even those of pharmacopeial grade—depends not solely on compound potency, but on the interplay between microbial architecture and compound formulation. PHMB exhibited relatively uniform antibiofilm activity across strains, likely supported by its physicochemical profile and formulation; however, exceptions were observed, highlighting the limitations of any single agent. In contrast, the activity of TEO, while more variable, was in part predictable based on biofilm phenotype, offering the possibility of targeted or optimized applications. This variability calls for the development of rationalized strategies for plant-derived antimicrobials in clinical or biotechnological contexts. Considering the growing interest in essential oils as complementary or alternative antimicrobials, we propose extending the concept of antiseptic stewardship to include essential oil stewardship—a framework that emphasizes compositional standardization, model-informed evaluation, and formulation optimization to ensure both reproducibility and relevance in real-world applications.

## Author Contributions

MB, ZS, AJ designed the research. MB, BD, NT, KK, and AJ performed the experiments. ZS performed the statistical analysis. MB and AJ wrote the first draft of the manuscript. MB, ZS, and AJ analyzed the data and prepared graphics. ZS and BD wrote sections of the manuscript. AJ, AM, and YS revised and edited the manuscript. AJ revised the English. AJ and MB supervised the work. All authors contributed to the article and approved the submitted version.

## Acknowledgments

We would like to express our gratitude to dr. Weronika Kozłowska from Division of Pharmaceutical Biotechnology, Department of Pharmaceutical Biology and Biotechnology, Wroclaw Medical University, Wroclaw, Poland for the help in the evaluation of essential oils composition.

