## Supplementary Material for "Toward Essential Oil Stewardship: Strain-Resolved Evaluation of Thyme Oil Activity Against *Pseudomonas aeruginosa*"

**Supplementary Table S1:** Statistical analysis of the data distribution for biofilm mass (a) or metabolic activity (b) of separate *P. aeruginosa* strains (n=10). Normal distribution was considered for values of p>0.05 (Shapiro-Wilk test). SD- standard deviation, IQR- interquartile range (IQR), N- data points.

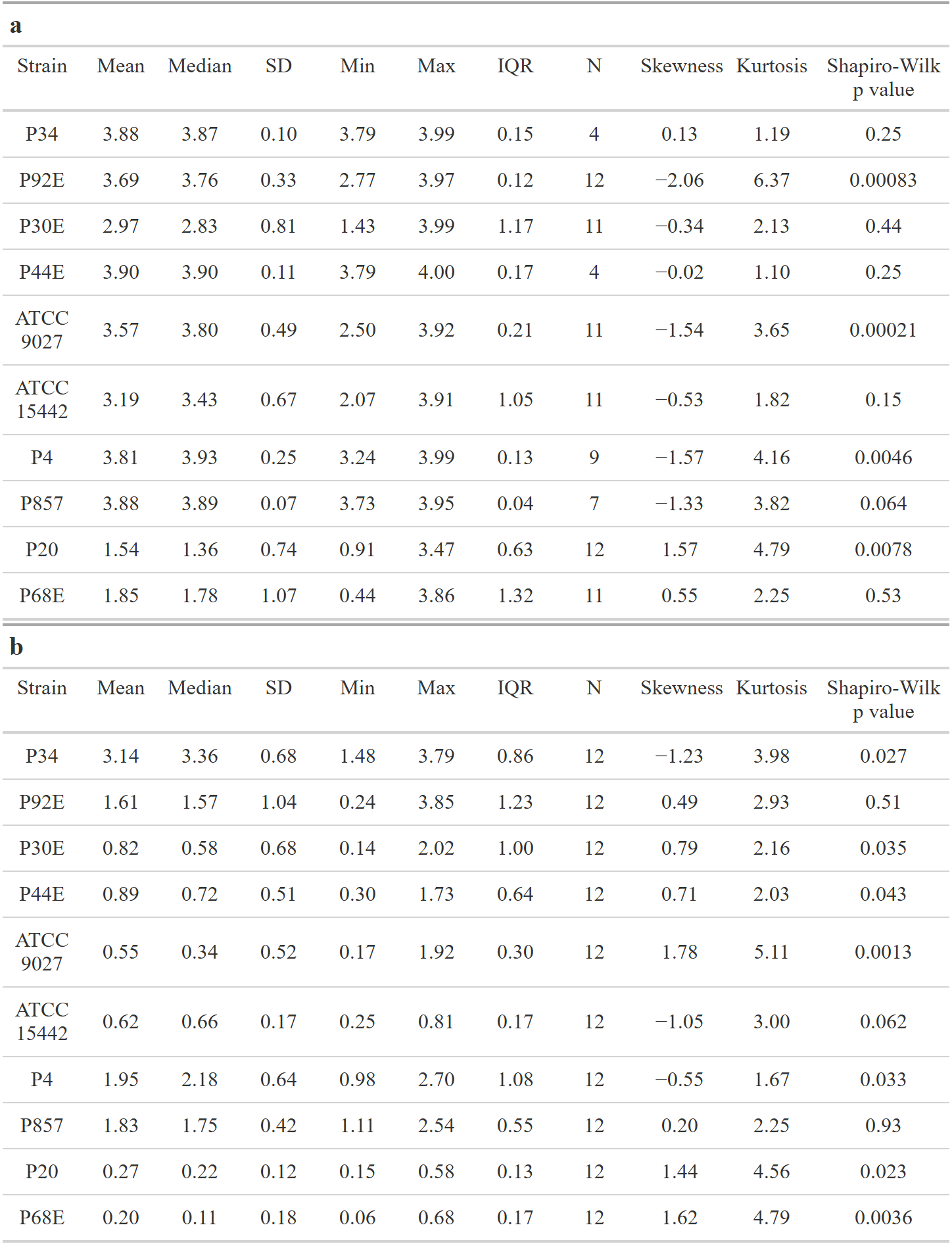

**Supplementary Table S2:** Parameters of the tested statistic of the differences in biological mass between particular *P. aeruginosa* strains. Welch’s ANOVA, followed by the Games-Howell test, was performed. Values of p<0.05 were considered significant, p<=0.05 was marked with one asterisk, p<=0.01 was marked with two asterisk, p<=0.001 was marked with three asterisk, and p<=0.0001 was marked with four asterisk. Ns- no significant differences, upper-lower Cl- confidence interval 95.

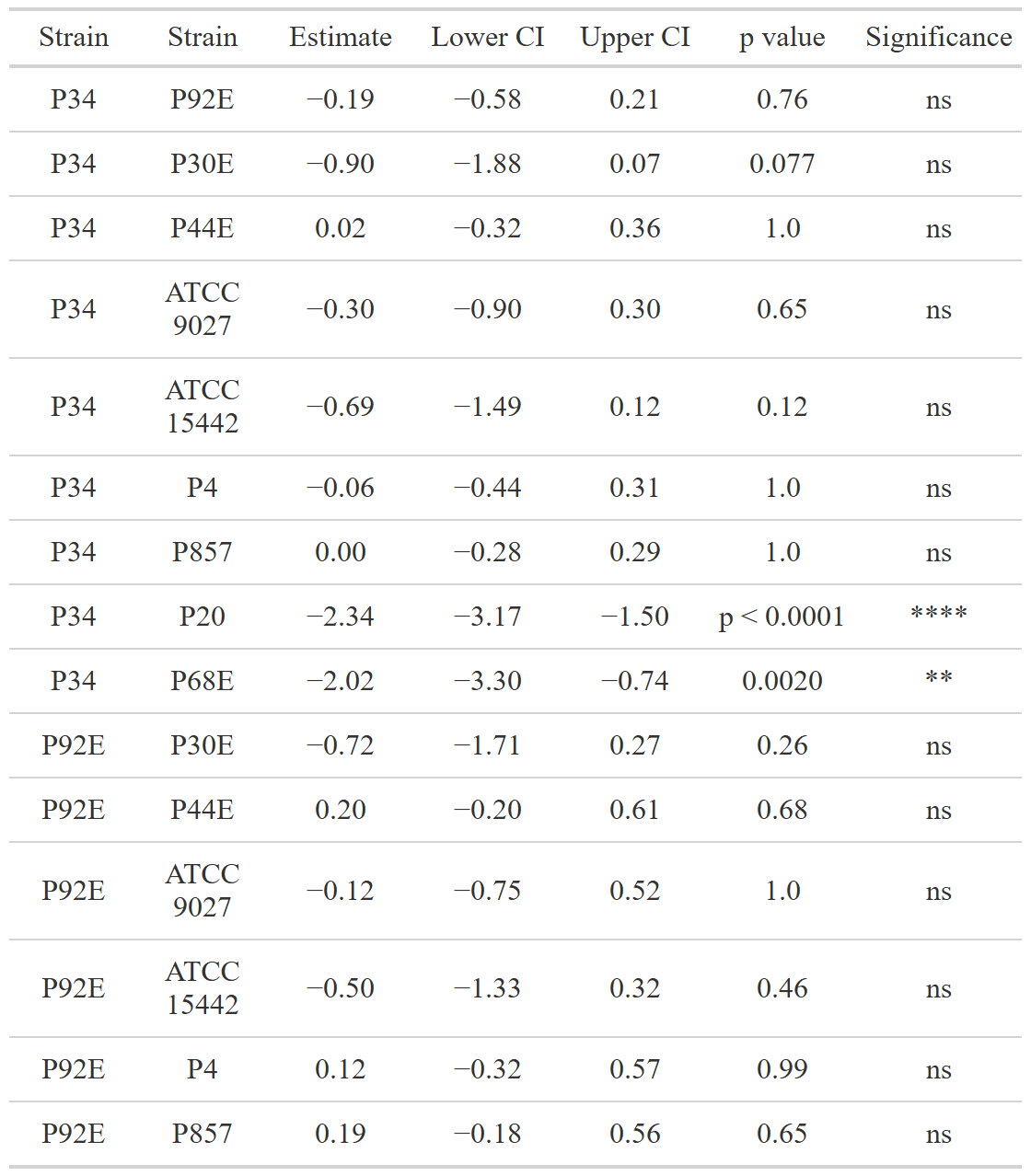

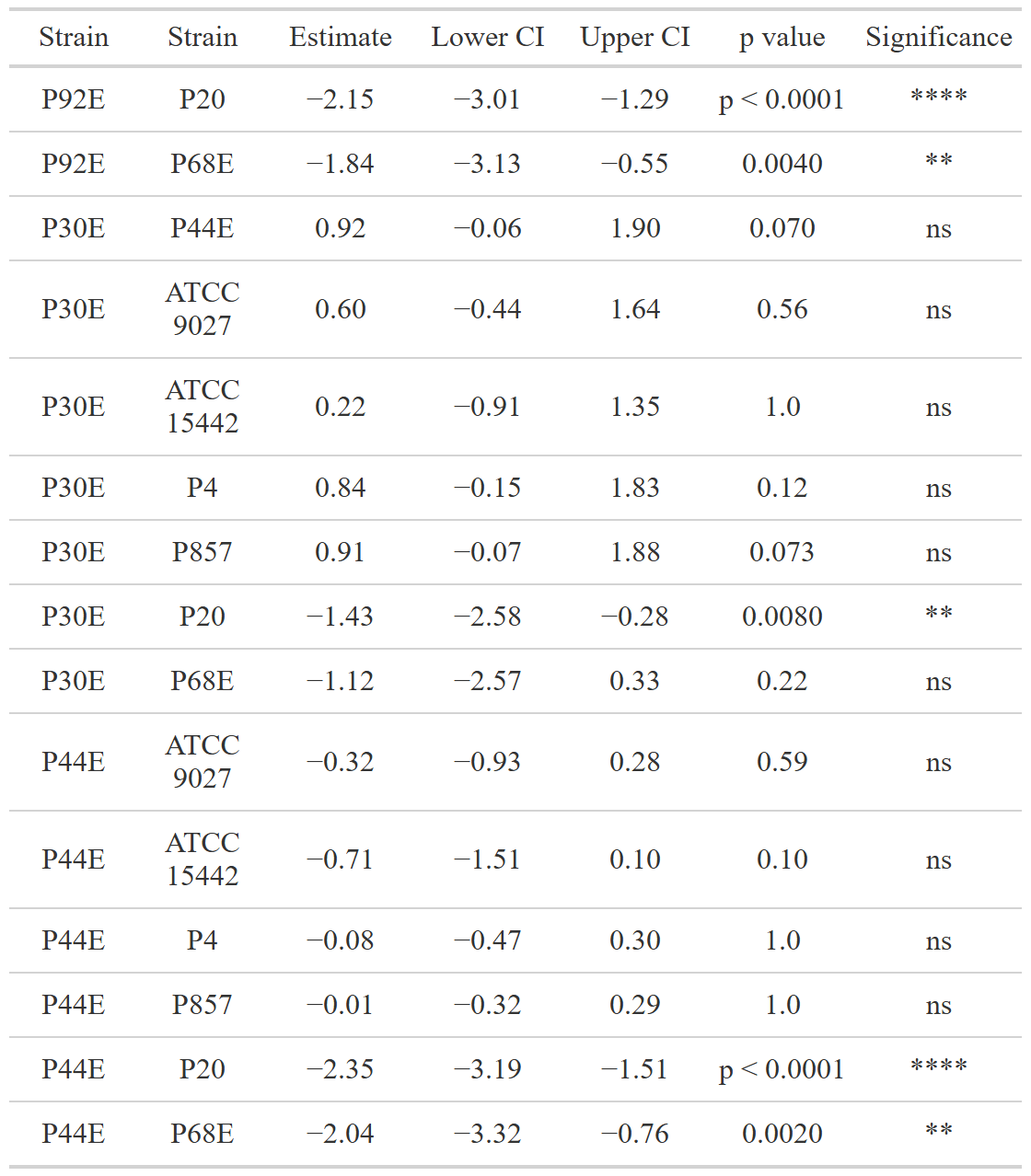

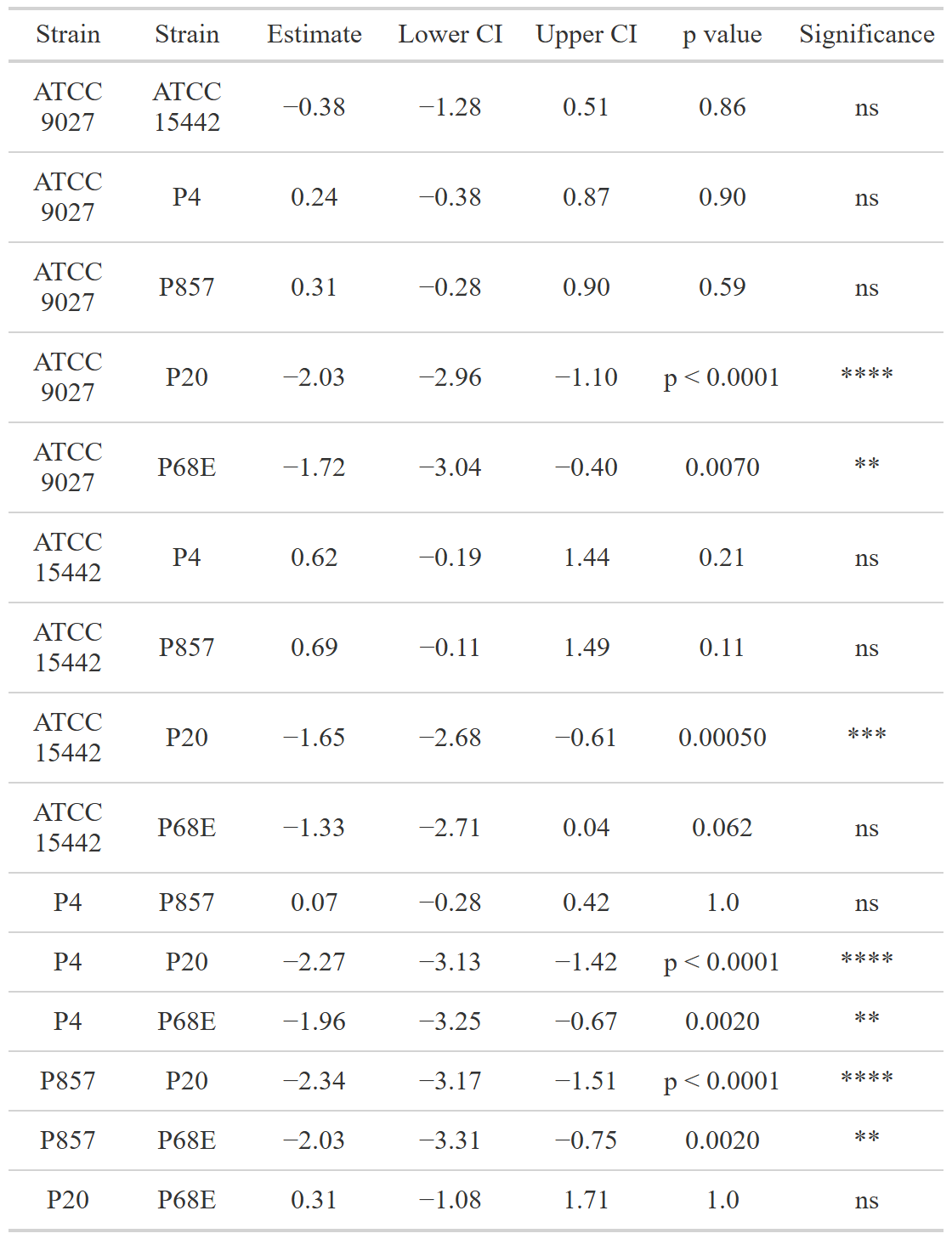

**Supplementary Table S3:** Parameters of the tested statistic of the differences in biofilm metabolic activity between particular *P. aeruginosa* strains. Welch’s ANOVA, followed by the Games-Howell test, was performed. Values of p<0.05 were considered significant, p<=0.05 was marked with one asterisk, p<=0.01 was marked with two asterisk, p<=0.001 was marked with three asterisk, and p<=0.0001 was marked with four asterisk. Ns- no significant differences, upper-lower Cl- confidence interval 95%.

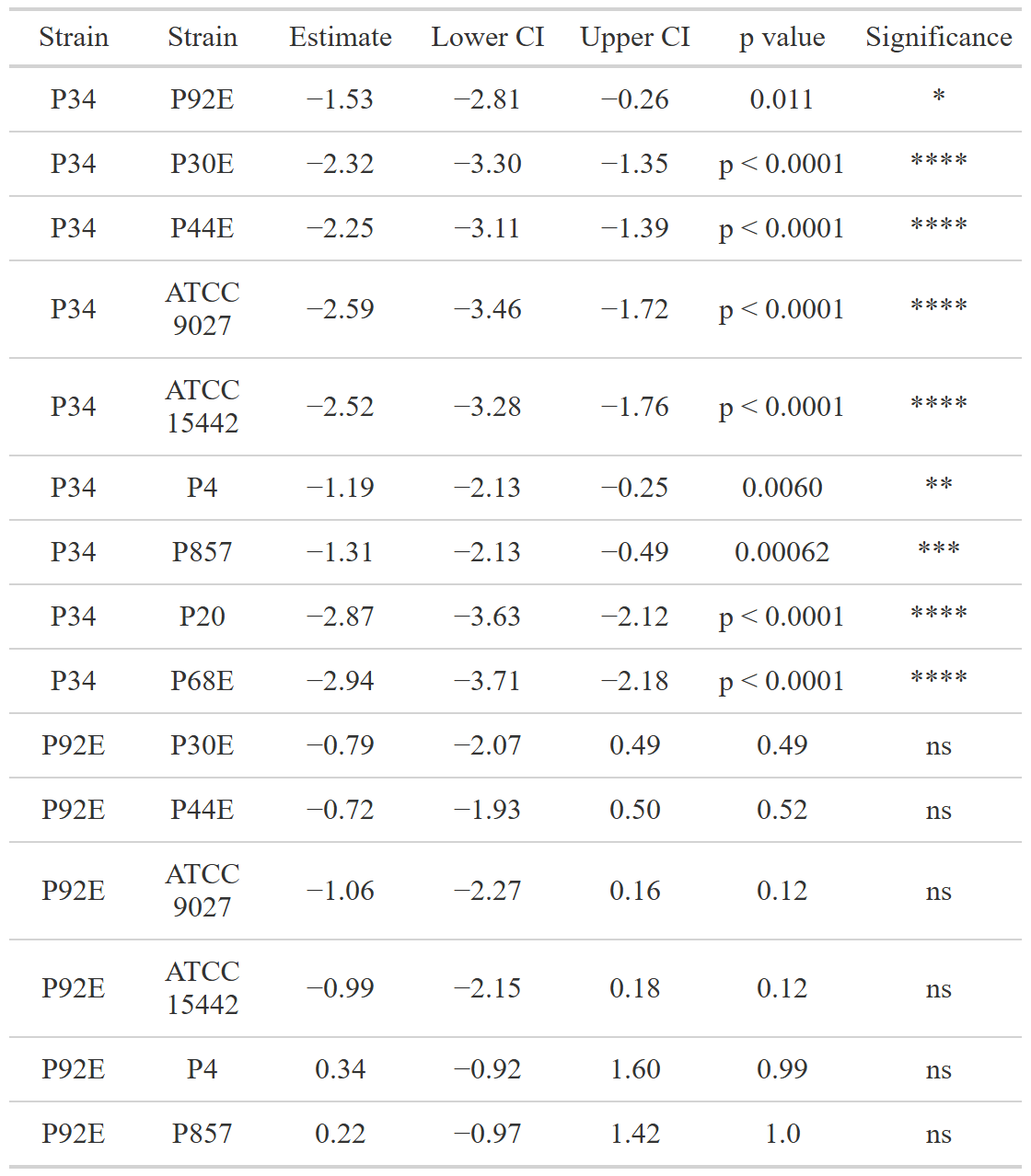

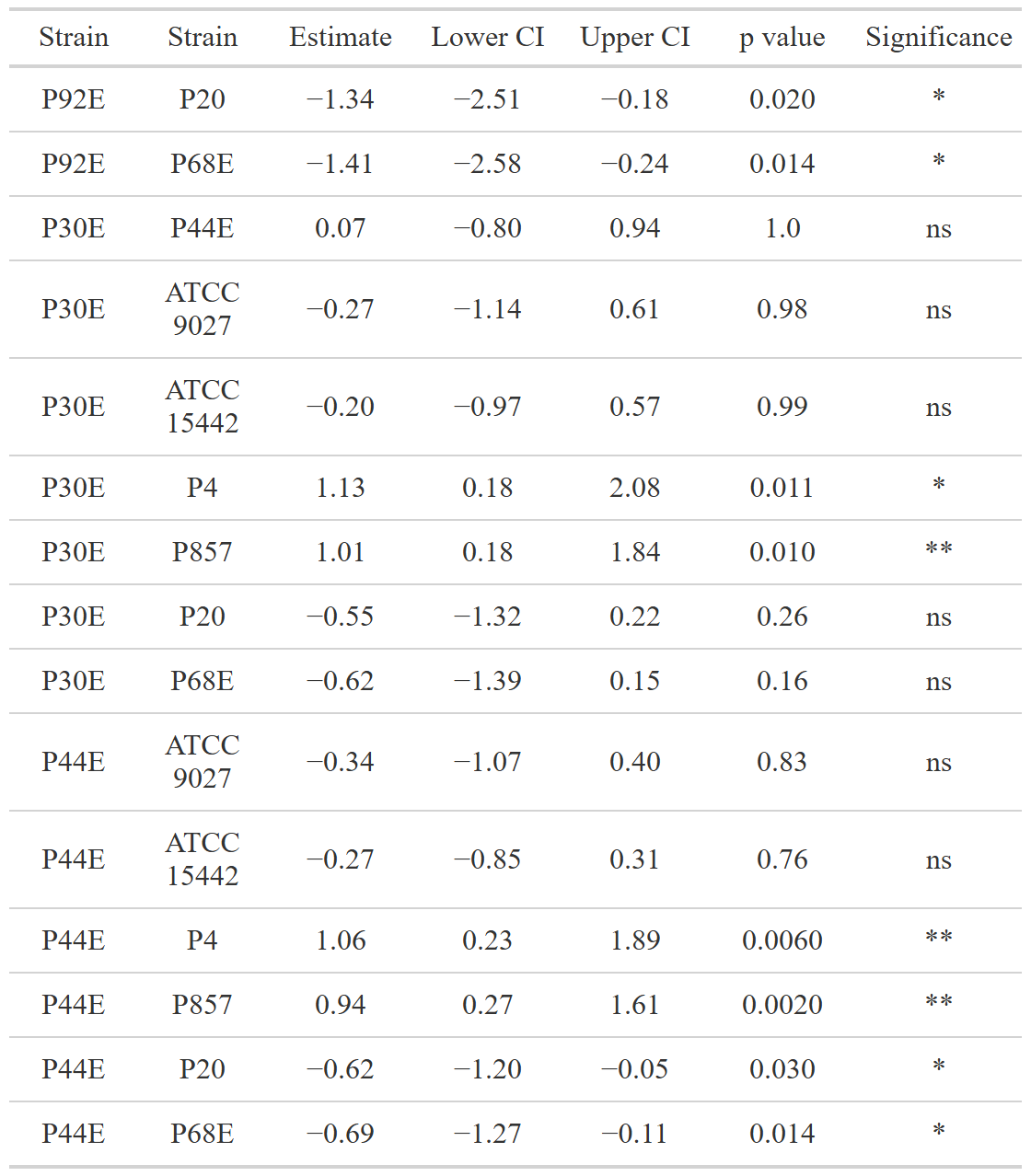

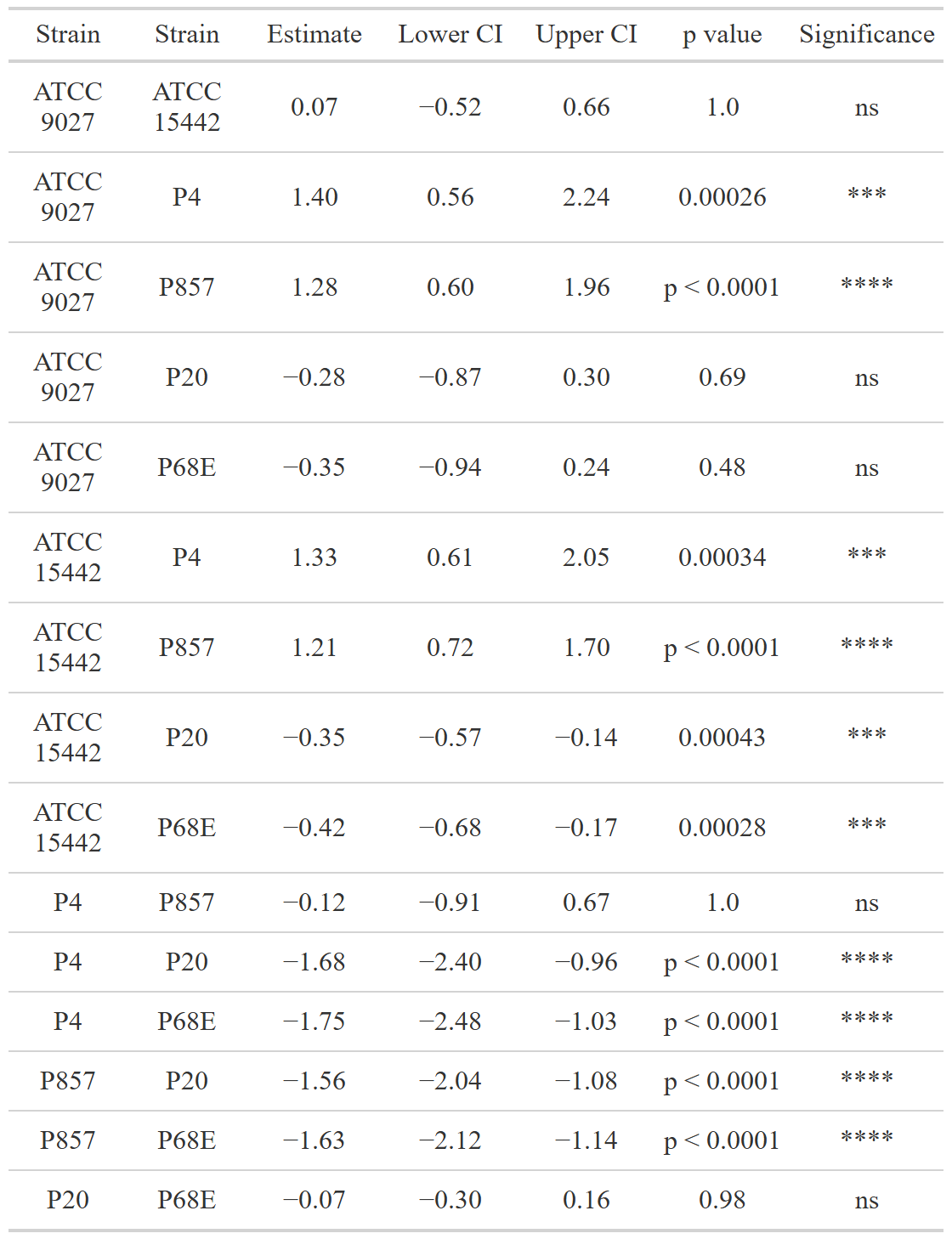

**Supplementary Table S4:** Parameters of the tested statistic of the differences in biofilm Colony-Forming-Unit (CFU/mL) number between particular *P. aeruginosa* strains. Dunn’s test, followed by the Kruskal-Wallis test, was performed. Values of p<0.05 were considered significant,  p<=0.05 was marked with one asterisk, and p<=0.01 was marked with two asterisk. Ns- no significant differences, N-data points.

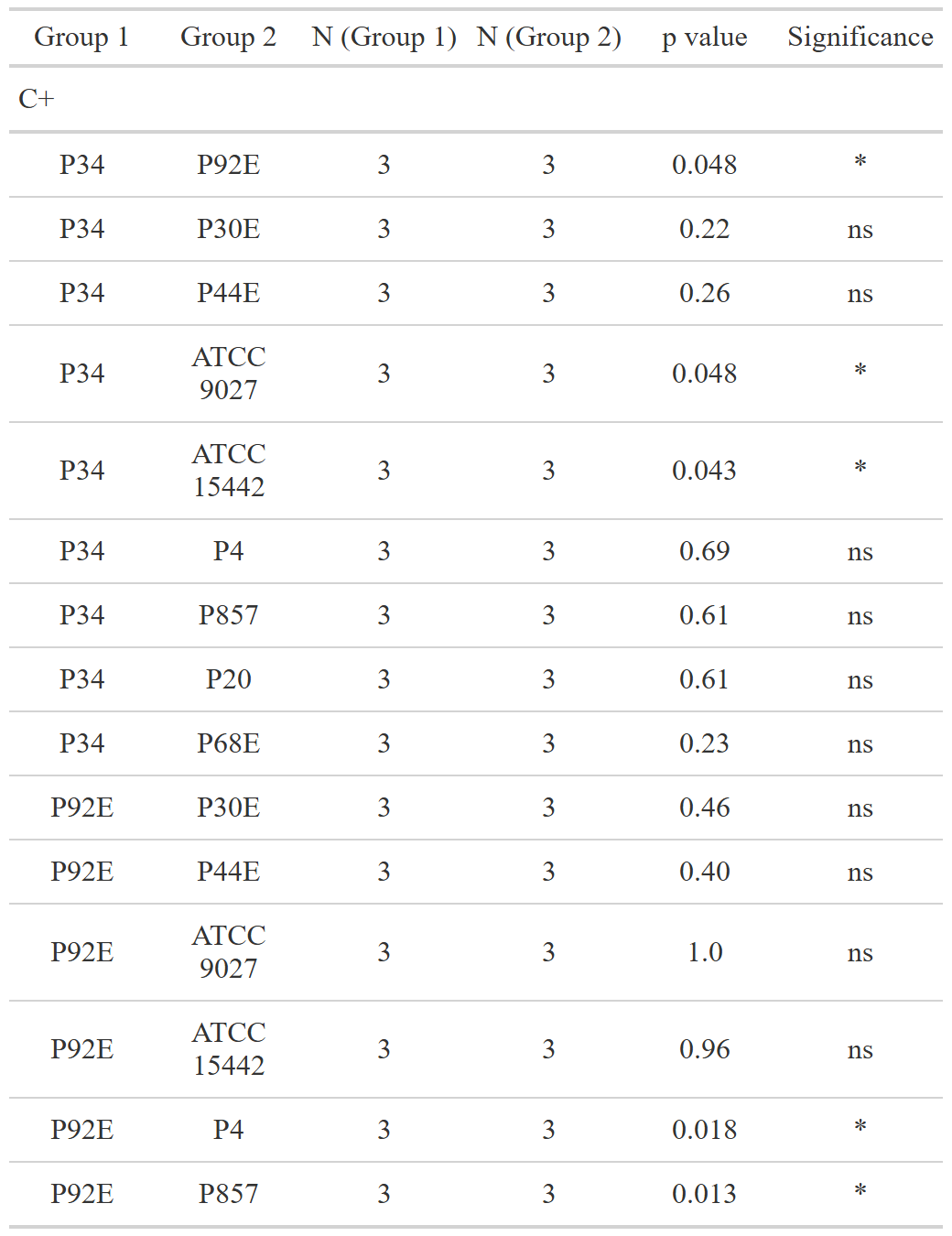

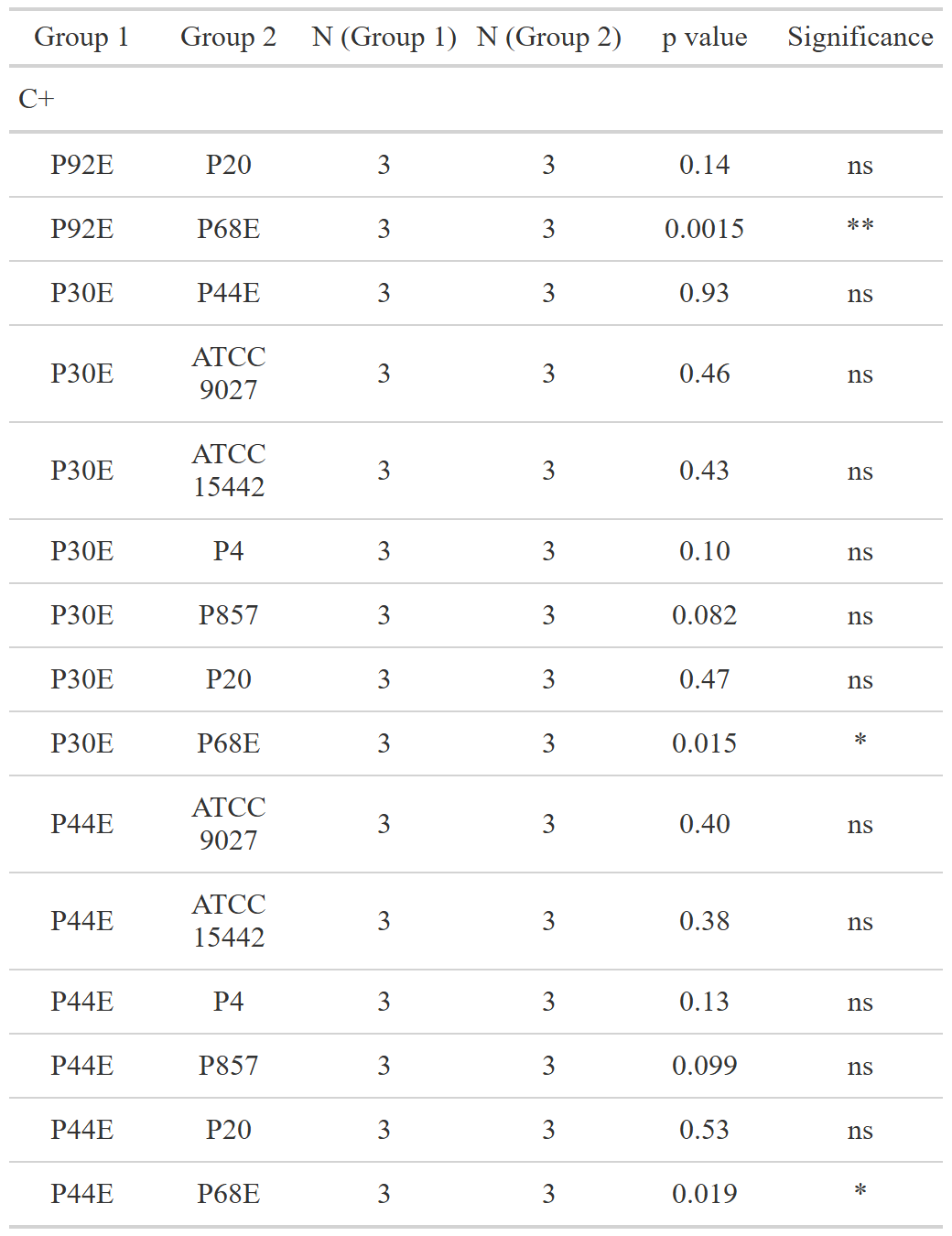

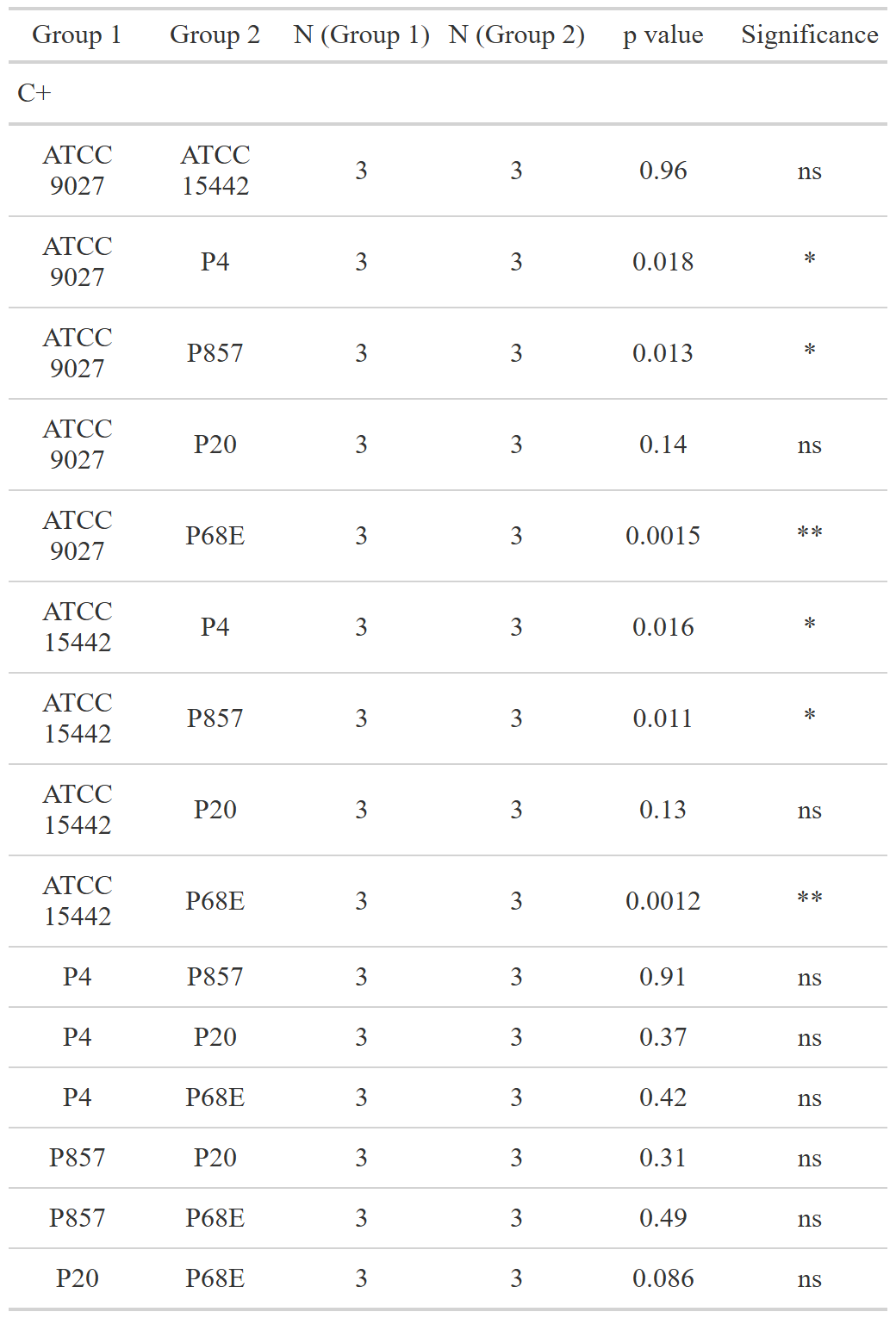

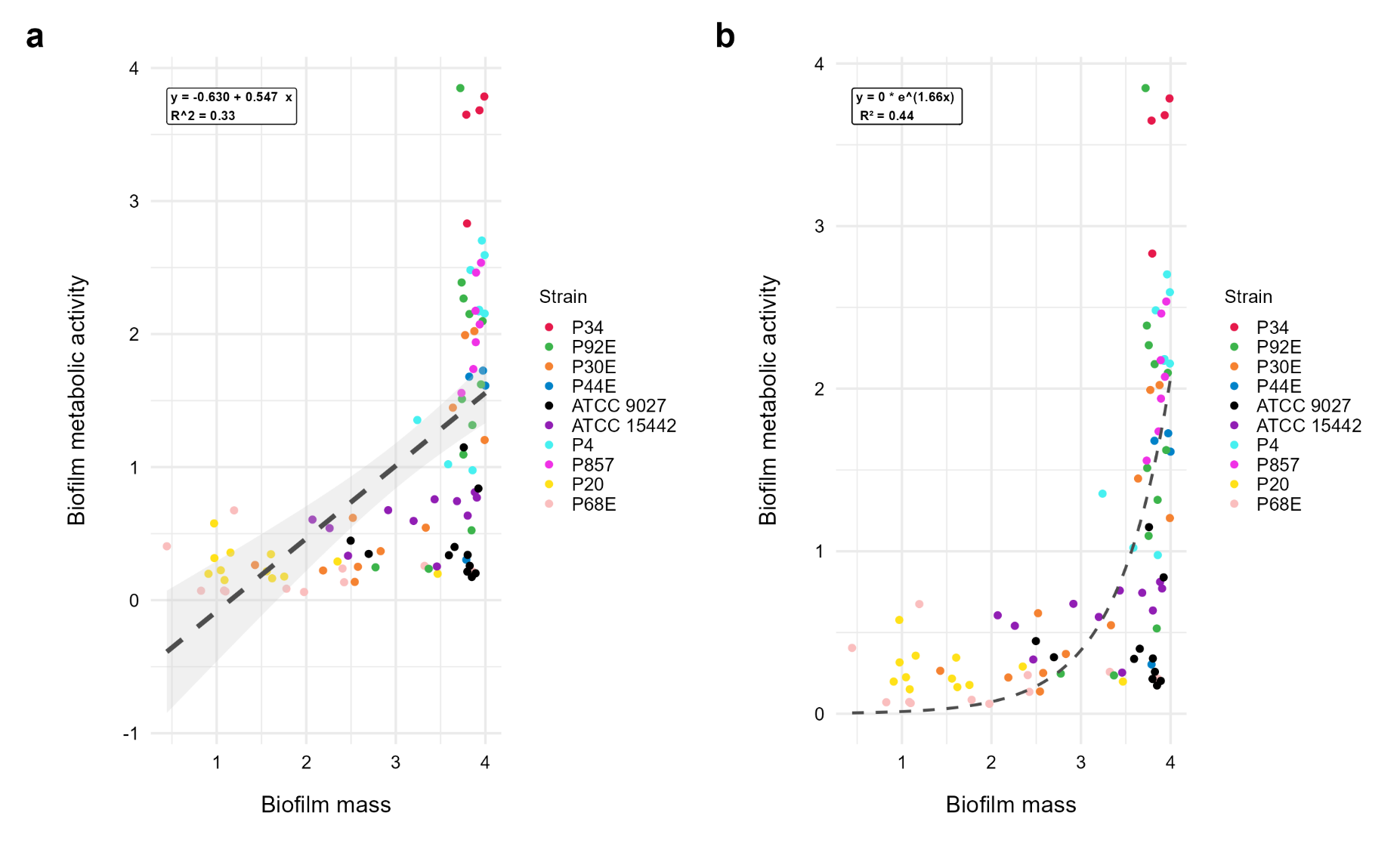

**Supplementary Fig S1:** Scatter plots of correlations of *P. aeruginosa* strains’ biofilm mass and metabolic activity. Data was fitted on (a) linear trend line, and (b) exponential trend line. The equation for the line of best fit, and R^2^- coefficient of determination are present on the top left corners of the respective panels.

**Supplementary Table S5:** Parameters of the tested statistic of the differences in susceptibility to tested compounds between particular *P. aeruginosa* strains expressed as growth inhibition zones (mm). Dunn’s test, followed by the Kruskal-Wallis test, was performed. Values of p<0.05 were considered significant,  p<=0.05 was marked with one asterisk, and p<=0.01 was marked with two asterisk. Ns- no significant differences, N- data points, TEO- Thyme Essential Oil, PHMB- polyhexanide.

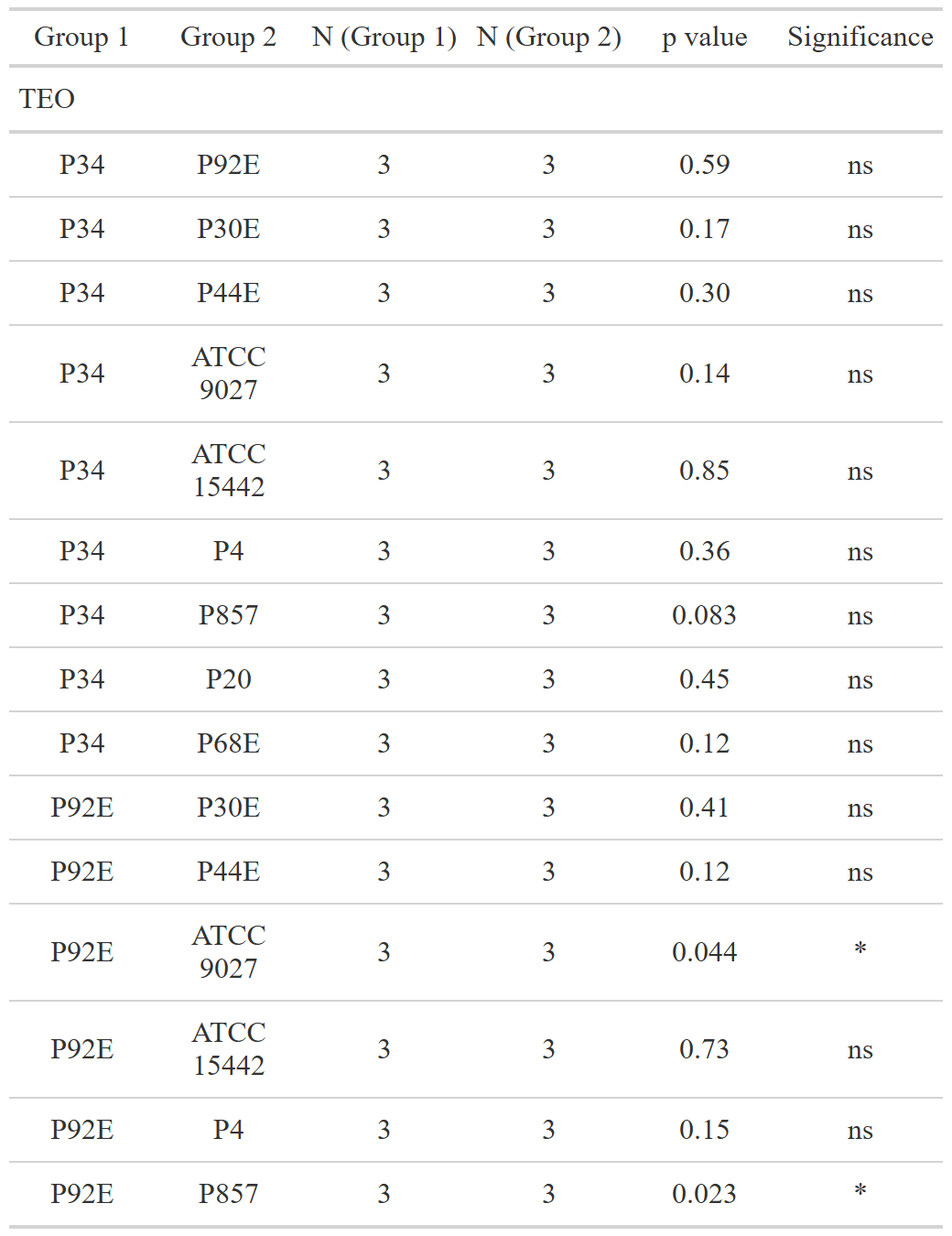

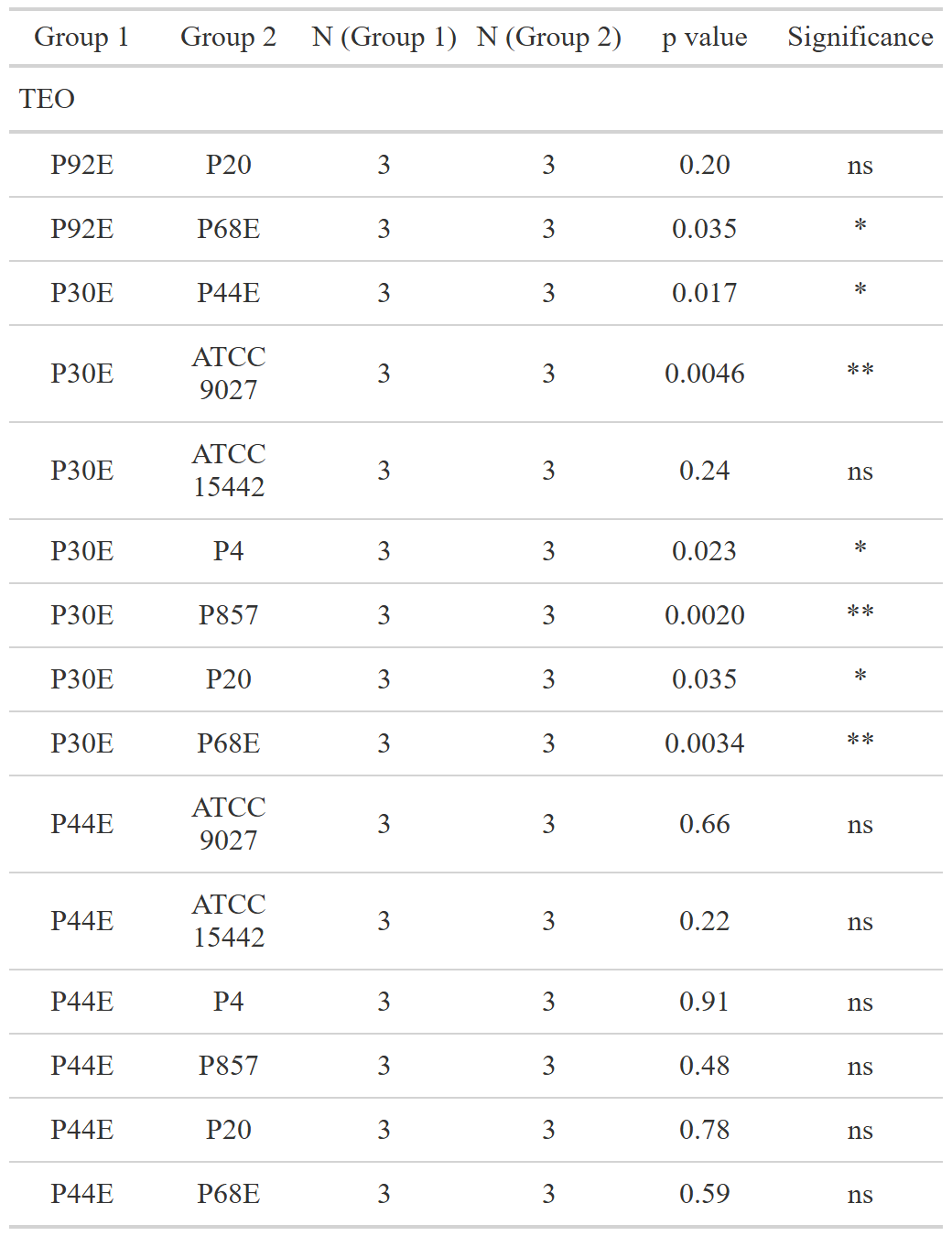

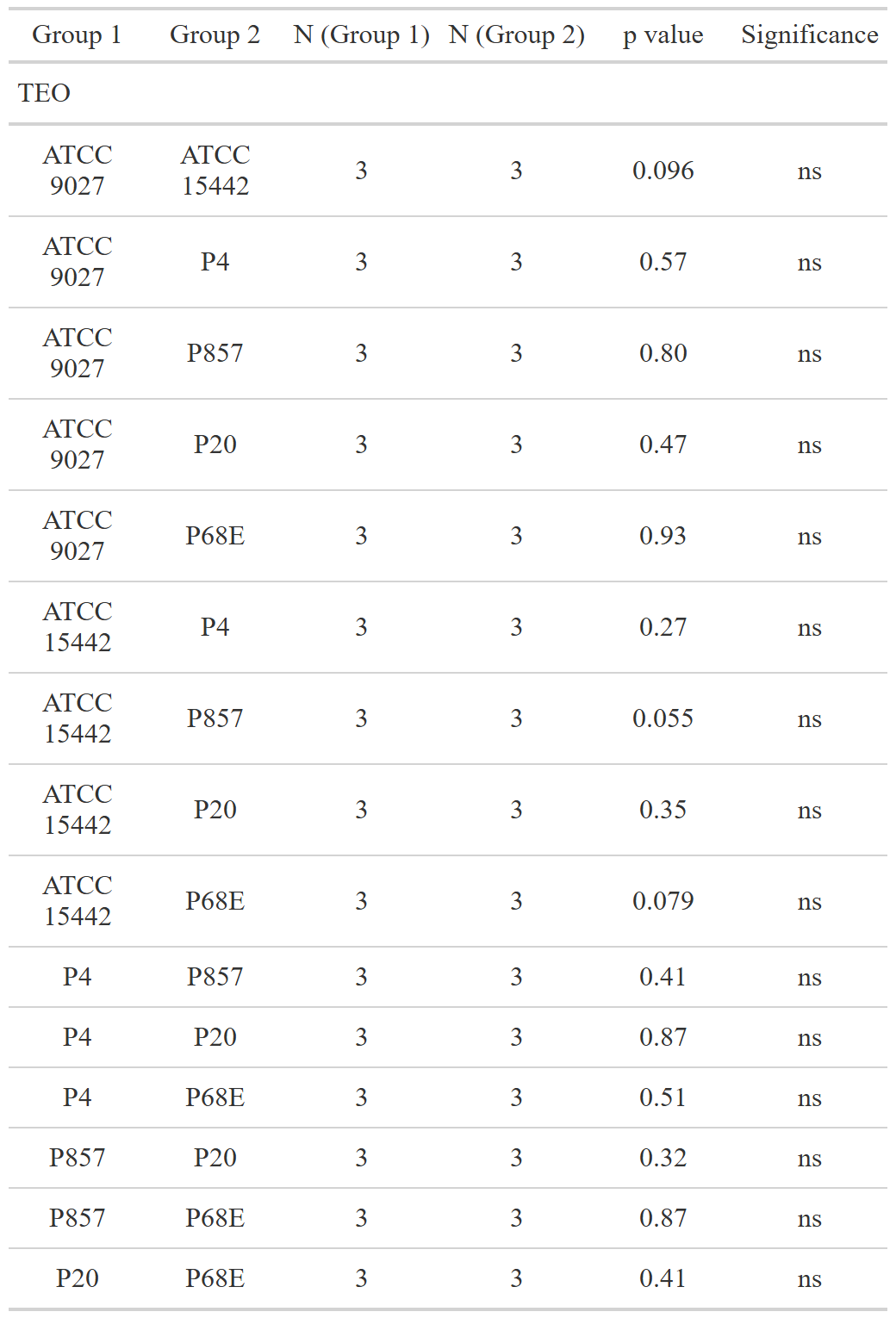

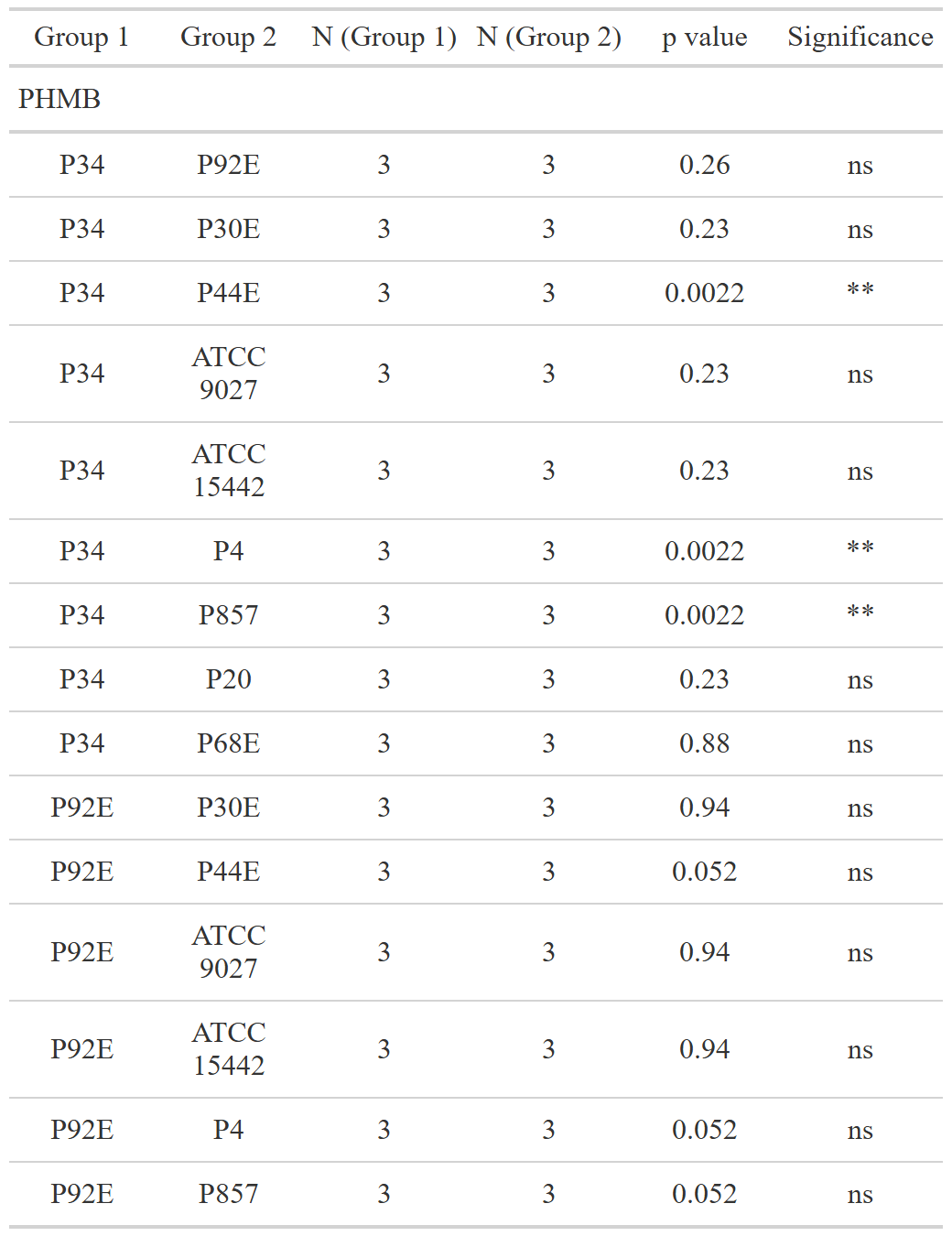

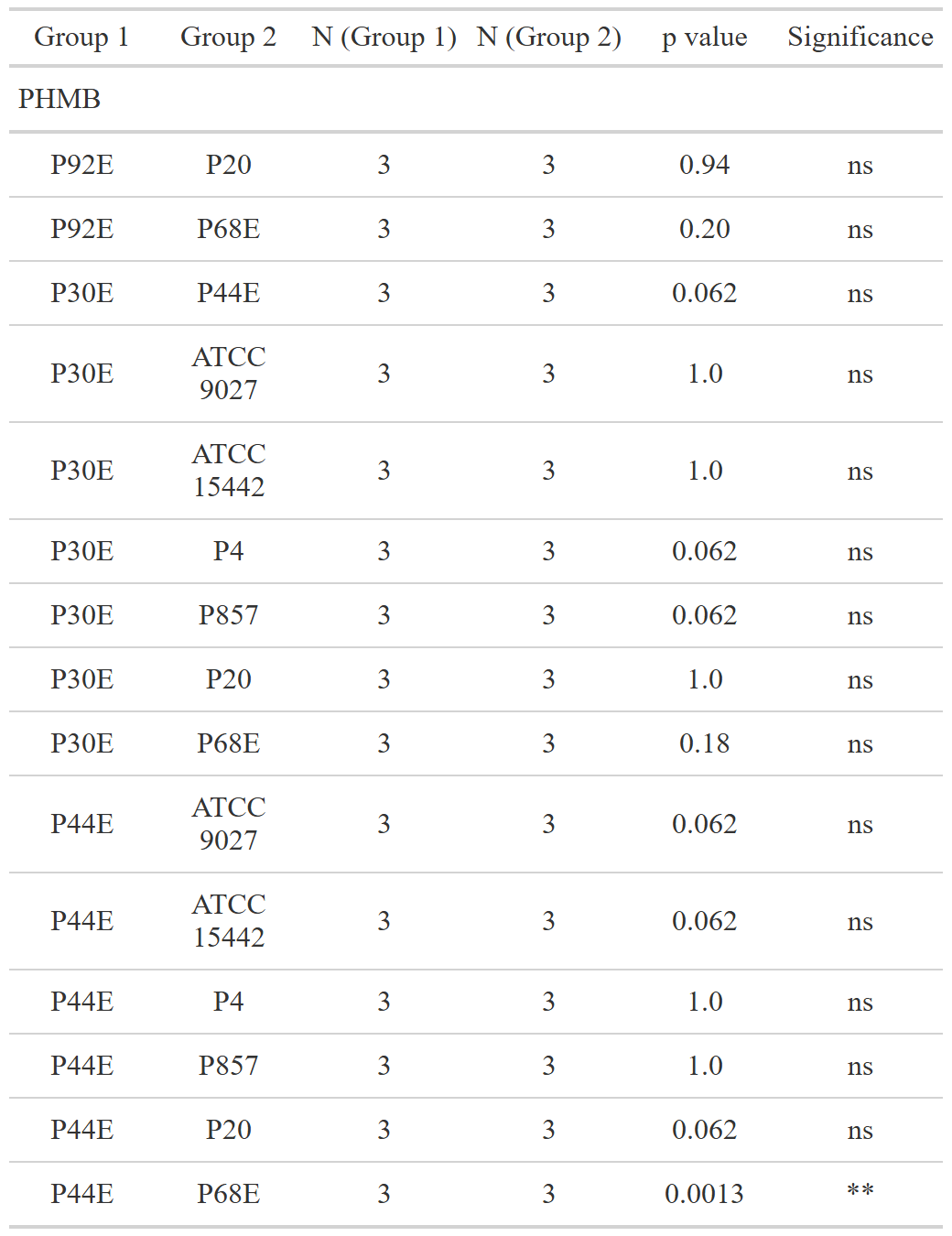

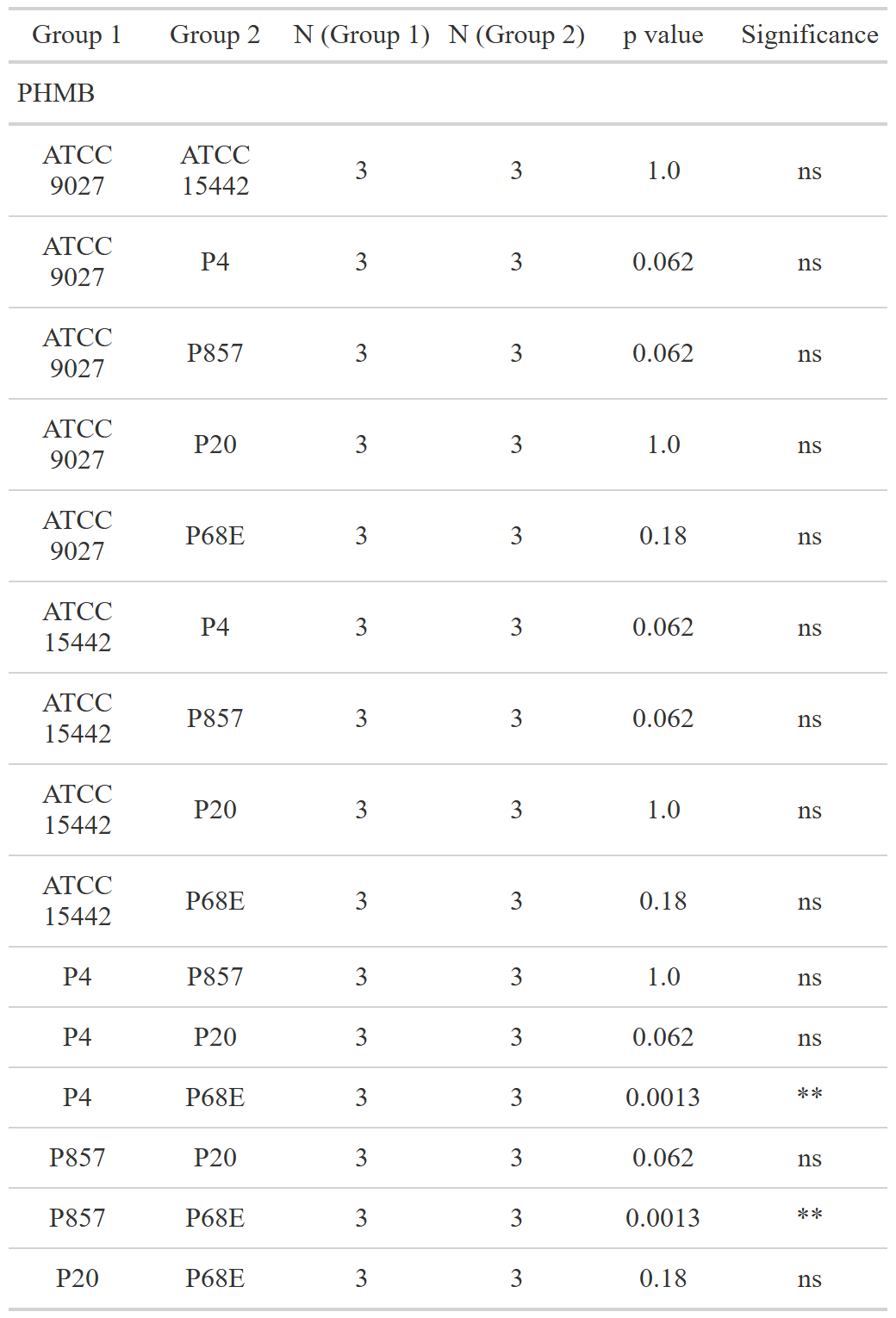

**Supplementary Table S6:** Parameters of the tested statistic of the differences in susceptibility to tested compounds between particular *P. aeruginosa* strains expressed as MIC (%, v/v) values (Minimal Inhibitory Concentration). Dunn’s test, followed by the Kruskal-Wallis test, was performed. Values of p<0.05 were considered significant,  p<=0.05 was marked with one asterisk, p<=0.01 was marked with two asterisk, and p<=0.001 was marked with three asterisk. Ns- no significant differences, N- data points, TEO- Thyme Essential Oil, PHMB- polyhexanide.

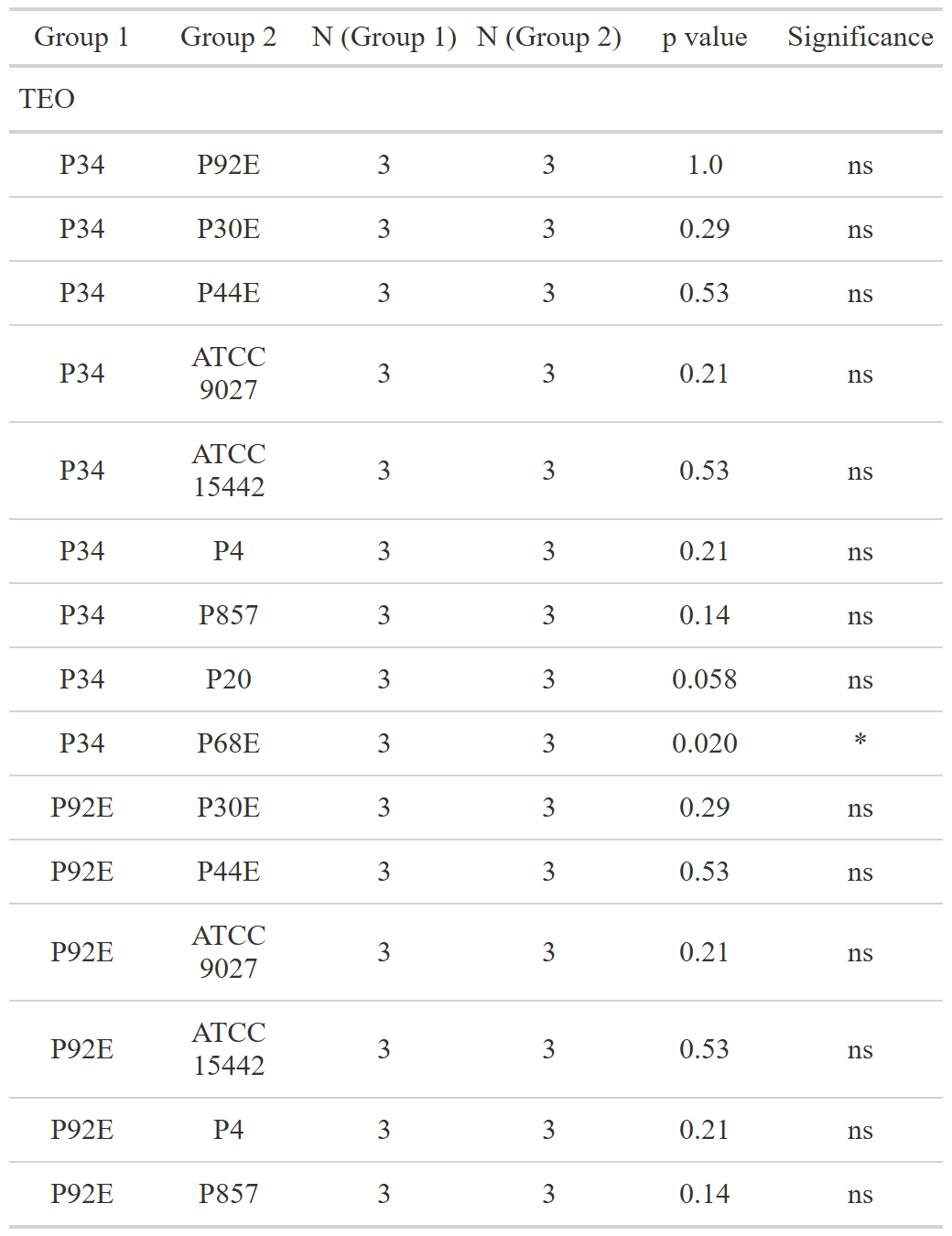

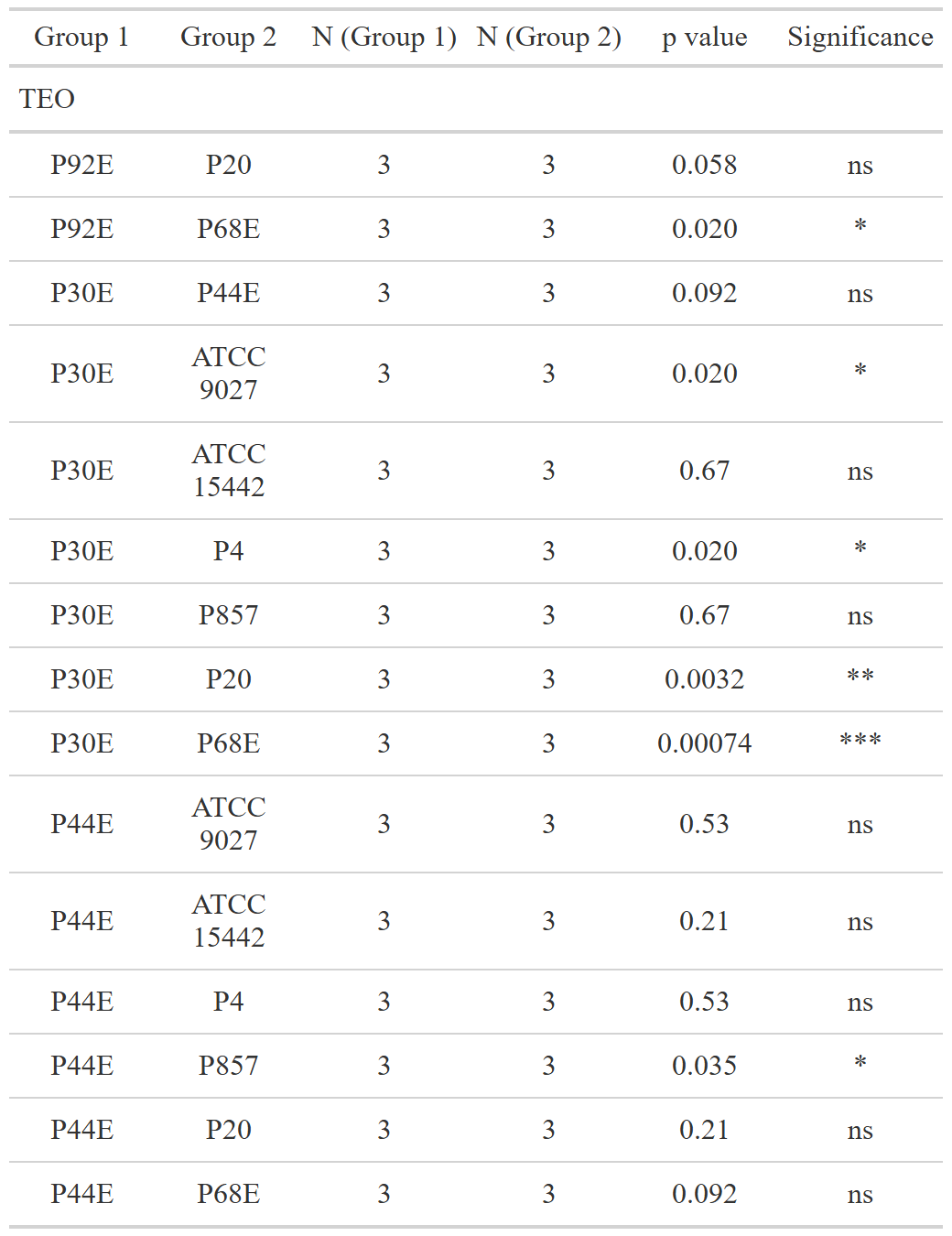

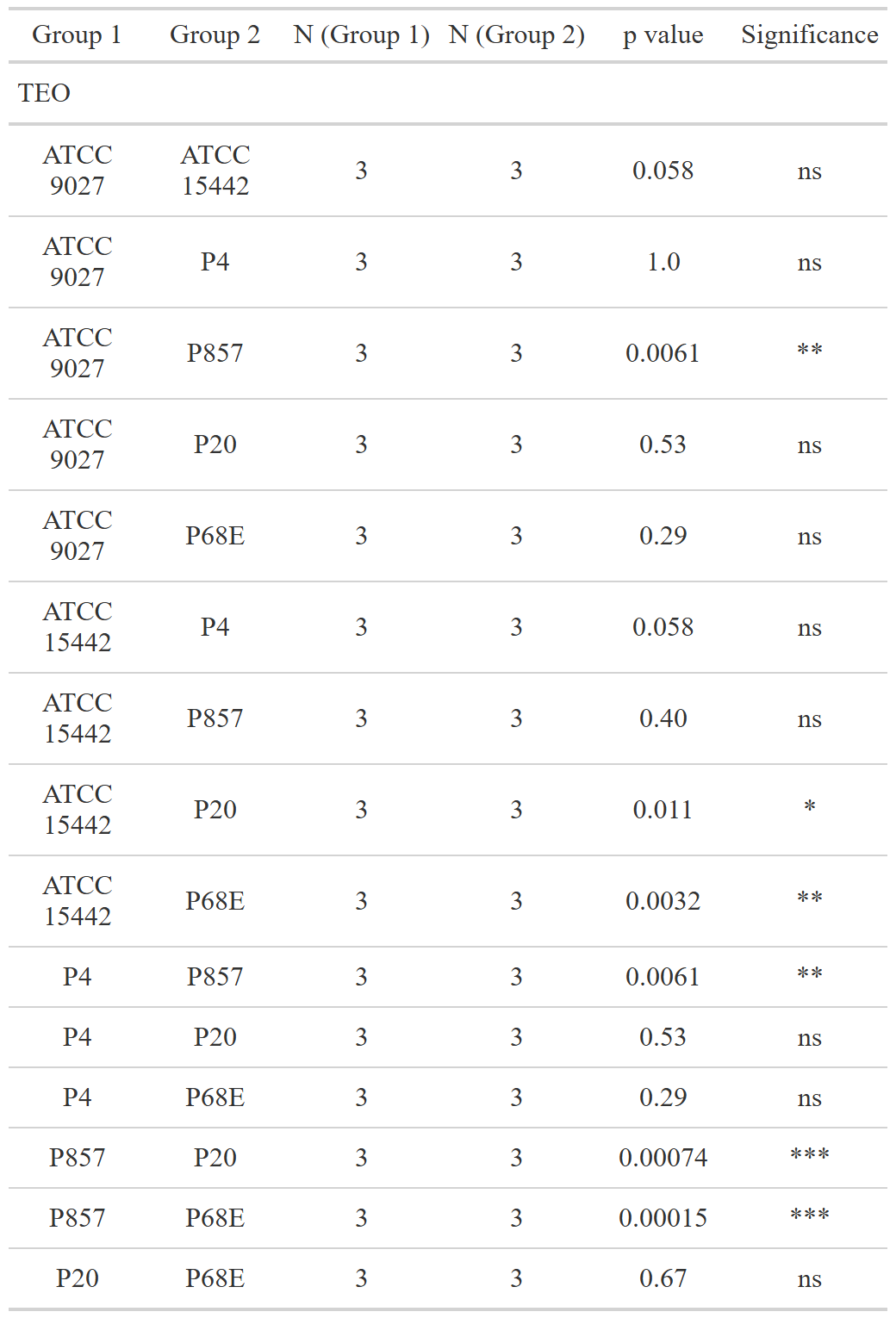

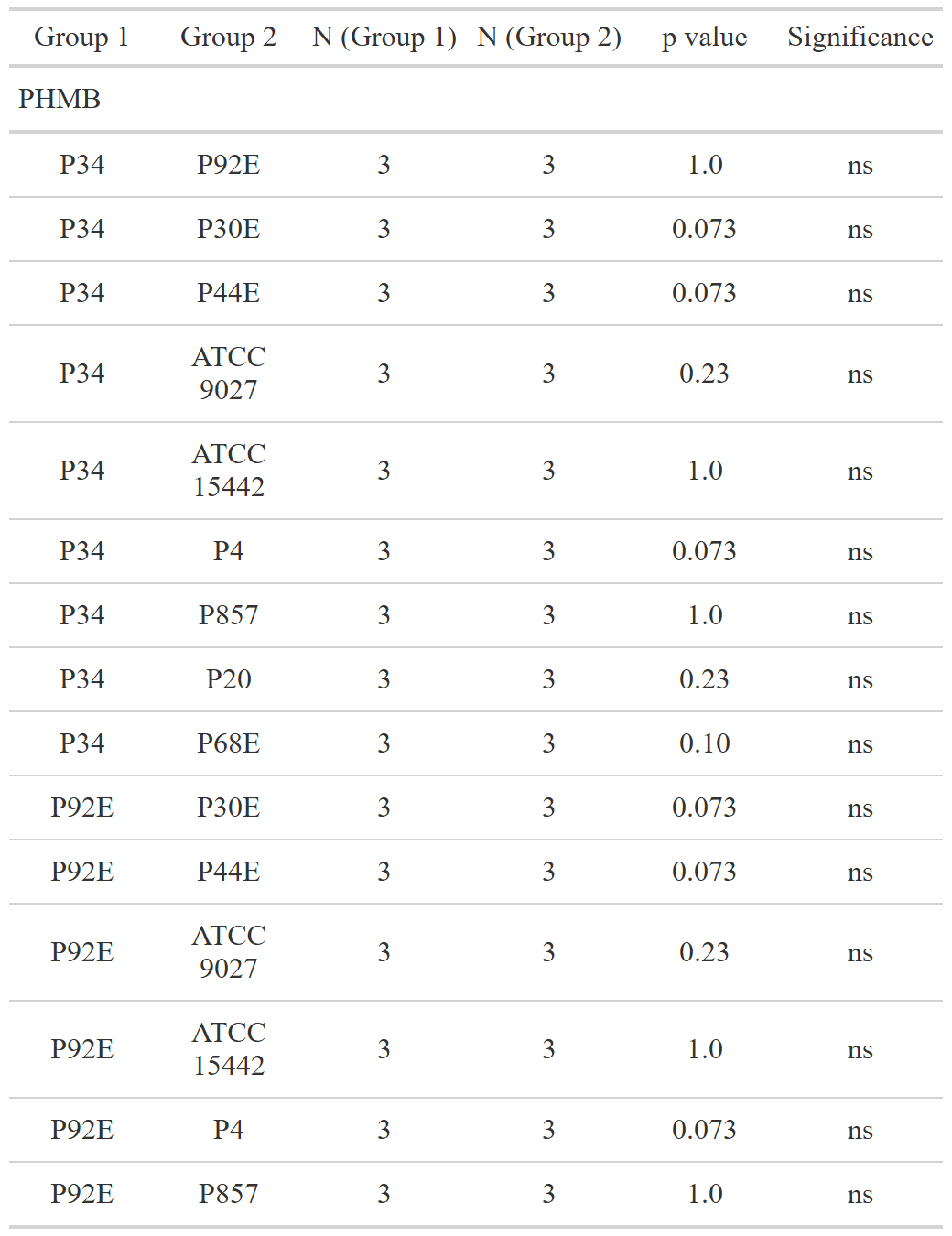

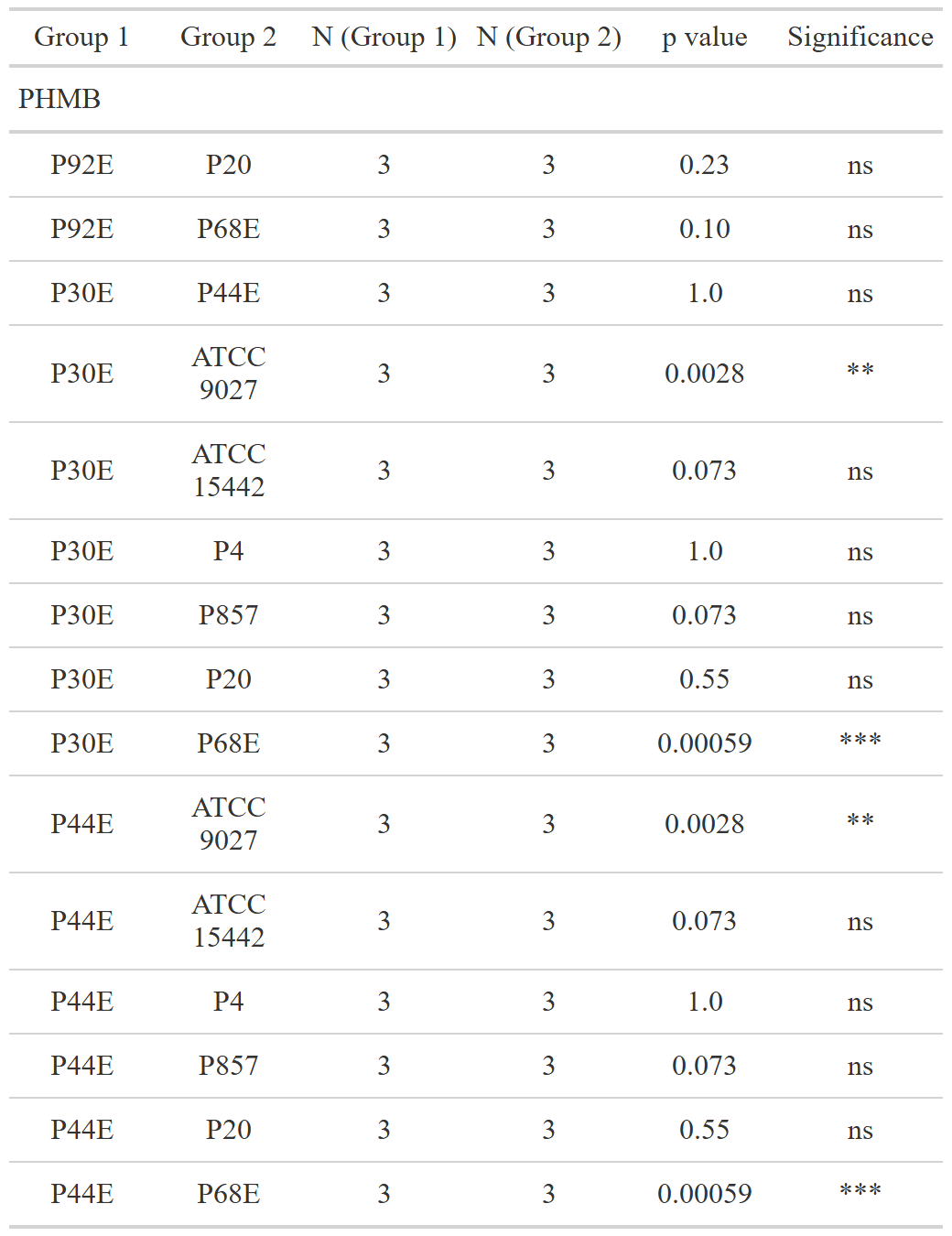

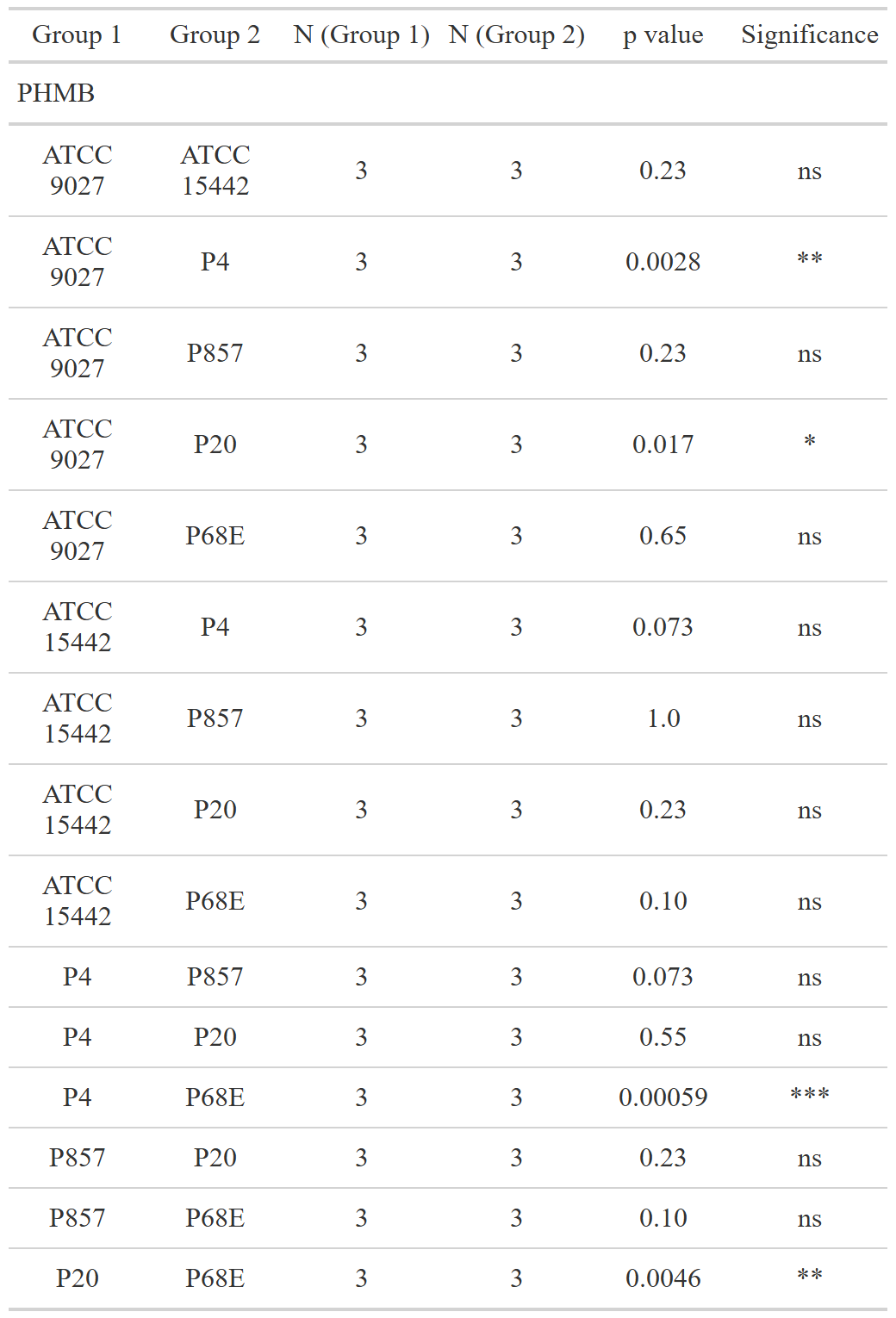

**Supplementary Table S7:** Biofilm CFU/mL values reduction across strains and conditions. TEO- Thyme Essential Oil, PHMB- polyhexanide, CFU- Colony-Forming Unit.

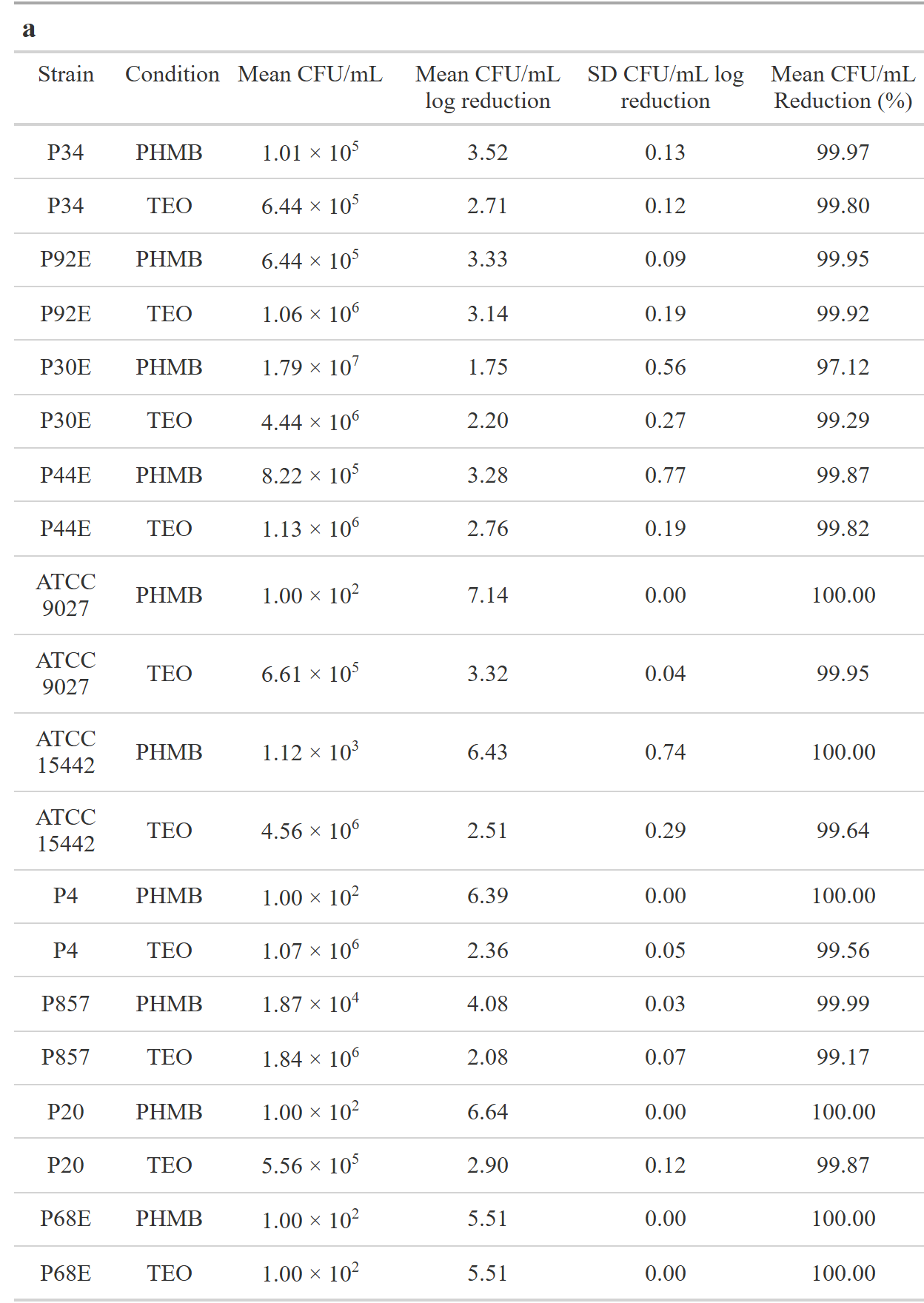

**Supplementary Table S8:** Parameters of the tested statistic of the differences in susceptibility to tested compounds between particular *P. aeruginosa* strains expressed as biofilm reduction (%). Dunn’s test, followed by the Kruskal-Wallis test, was performed. Values of p<0.05 were considered significant,  p<=0.05 was marked with one asterisk, p<=0.01 was marked with two asterisk, and p<=0.001 was marked with three asterisk. Ns- no significant differences, N- data points, TEO- Thyme Essential Oil, PHMB- polyhexanide.

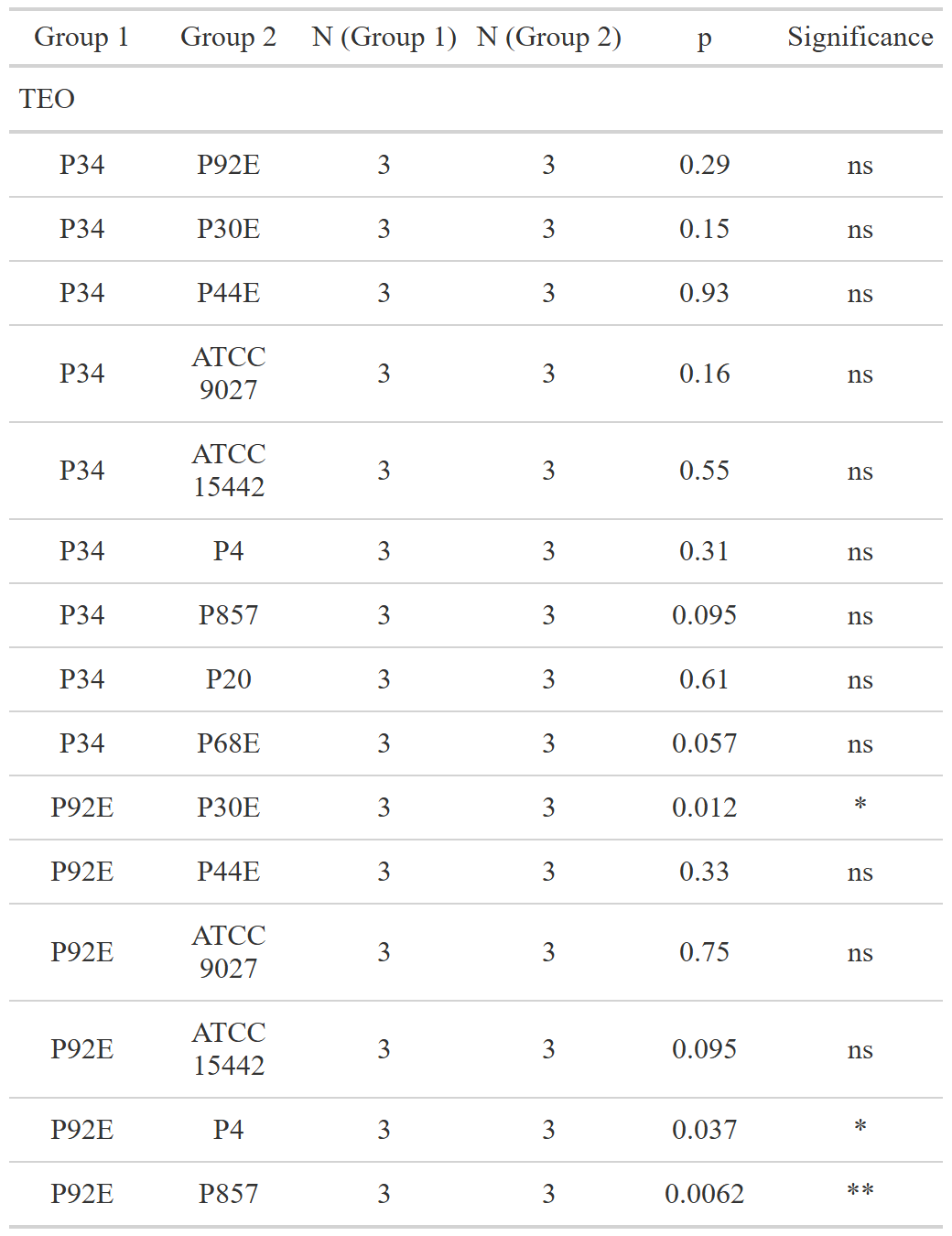

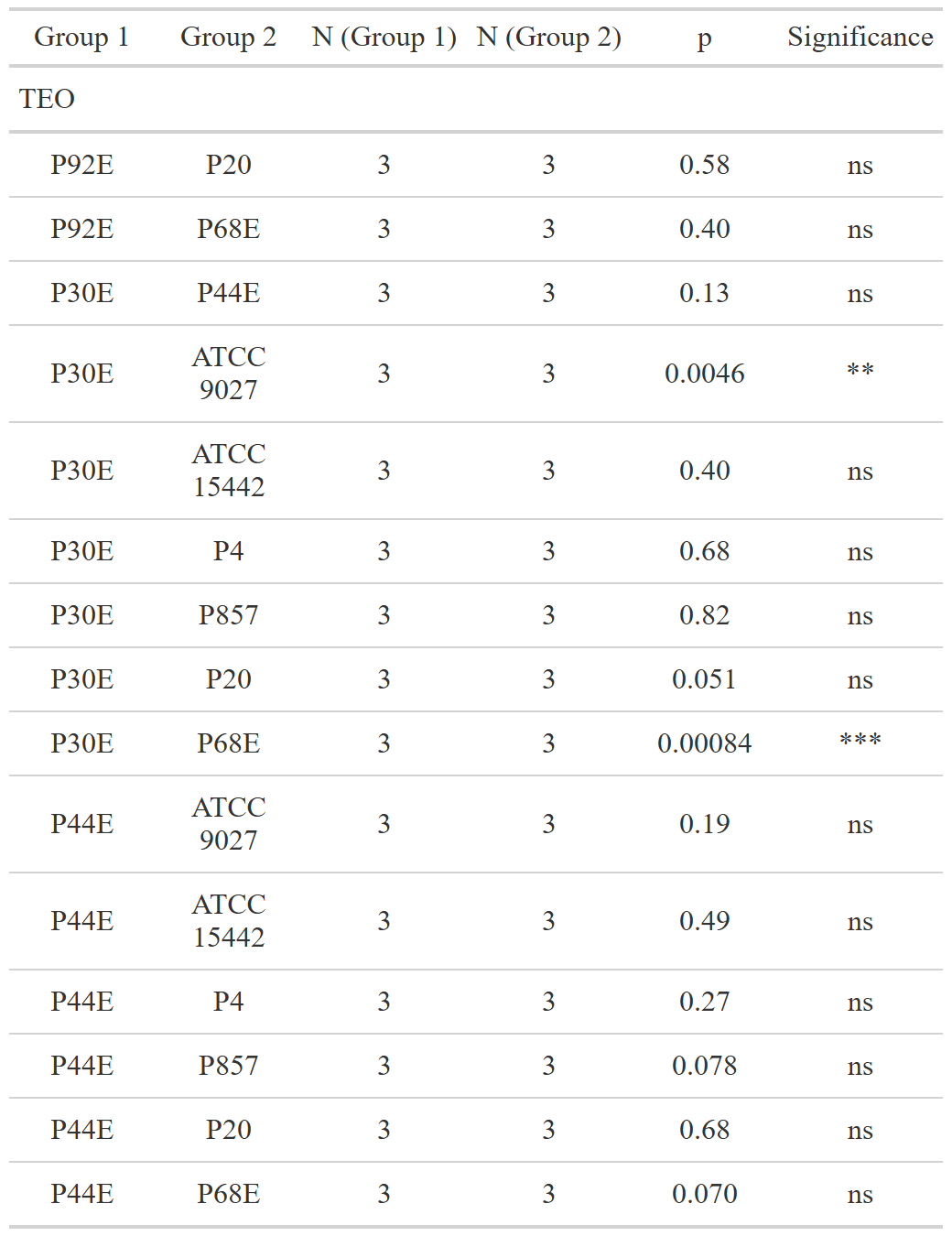

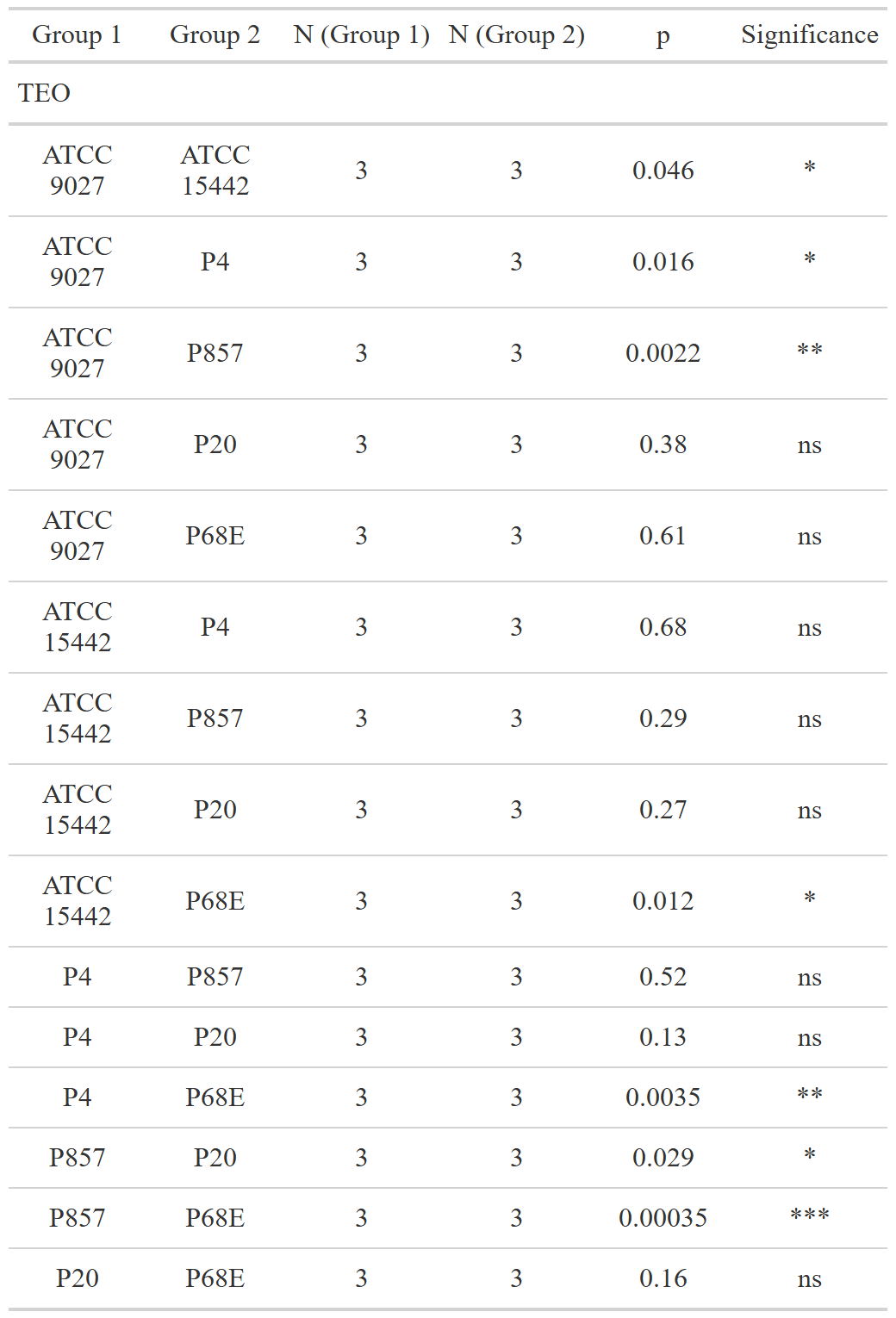

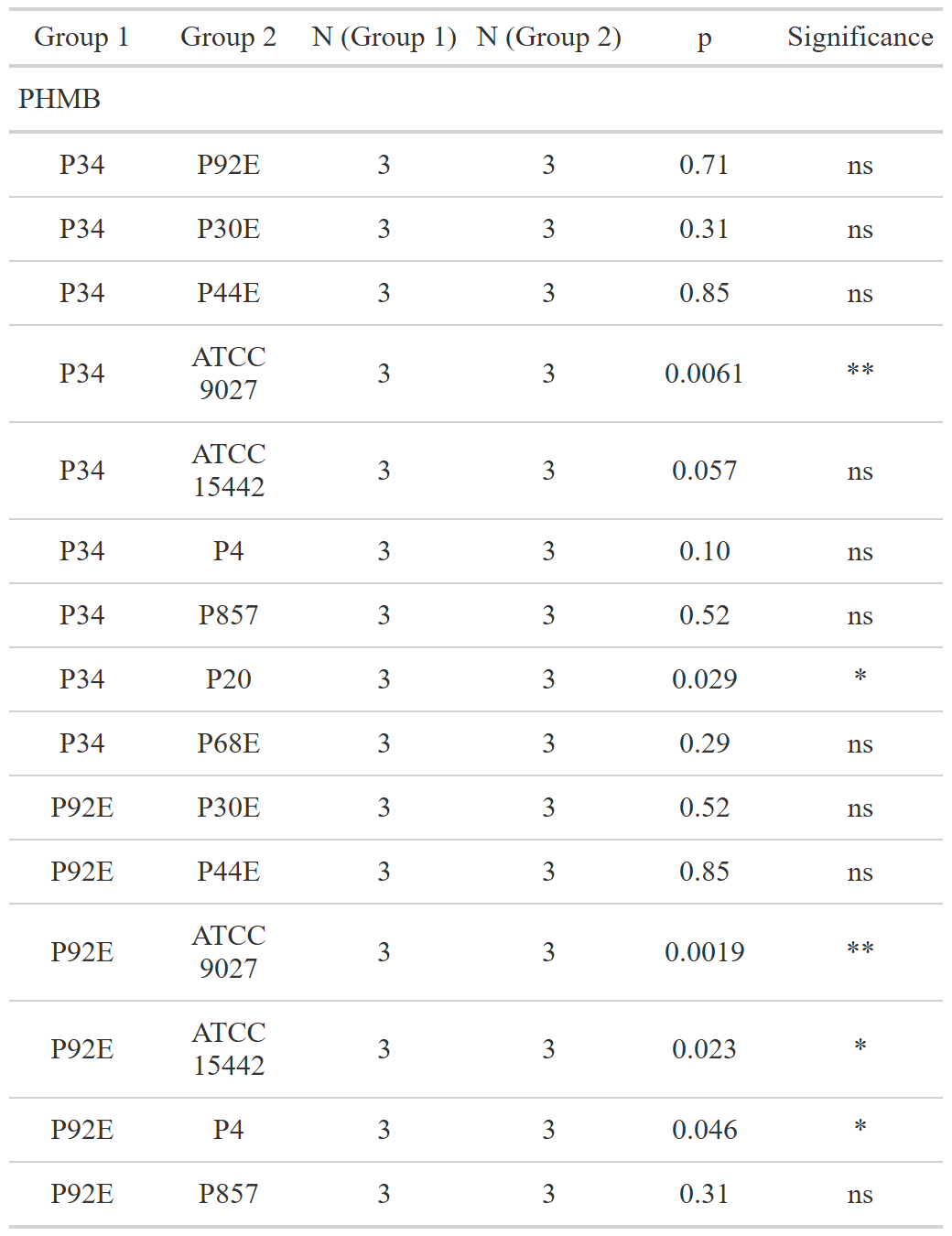

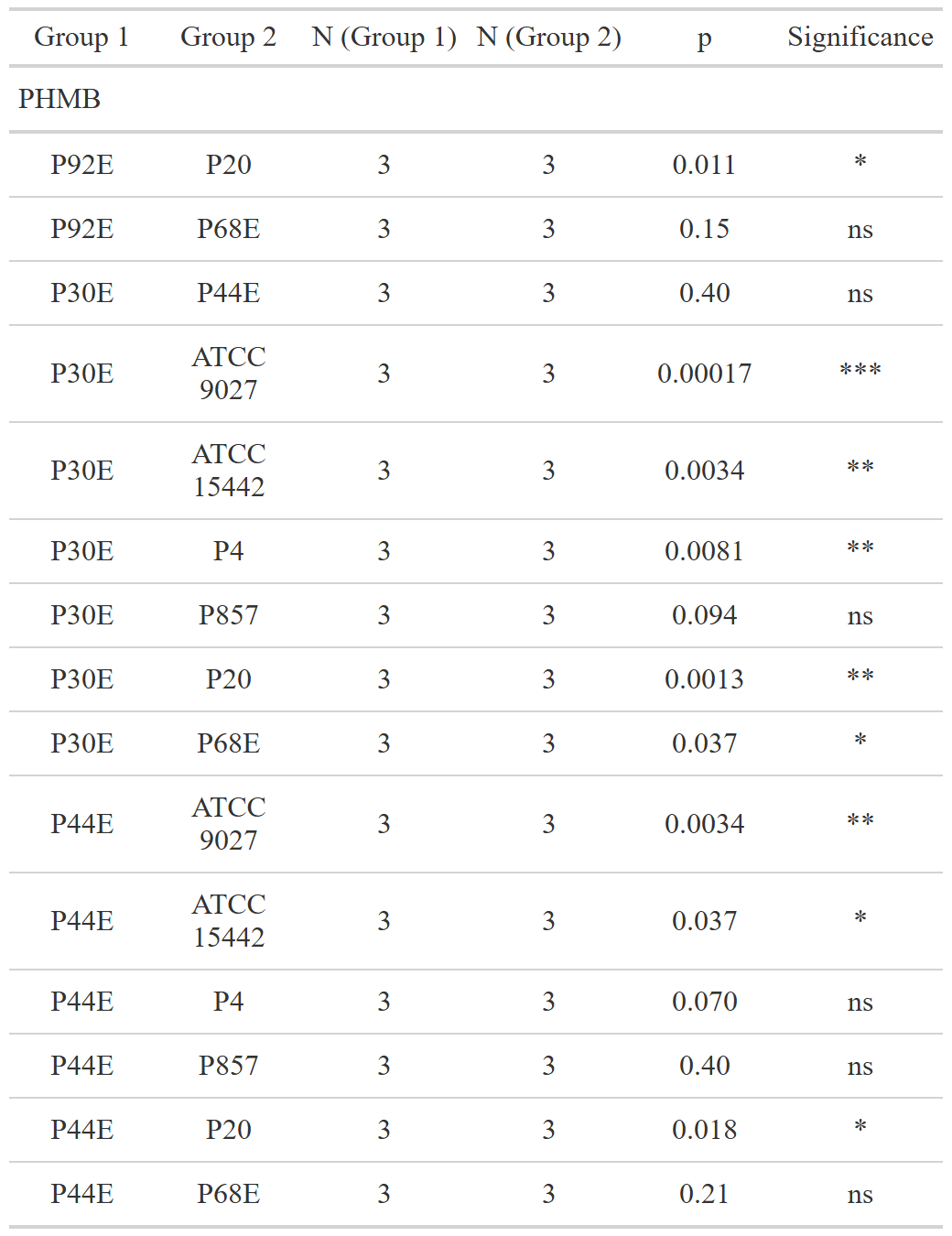

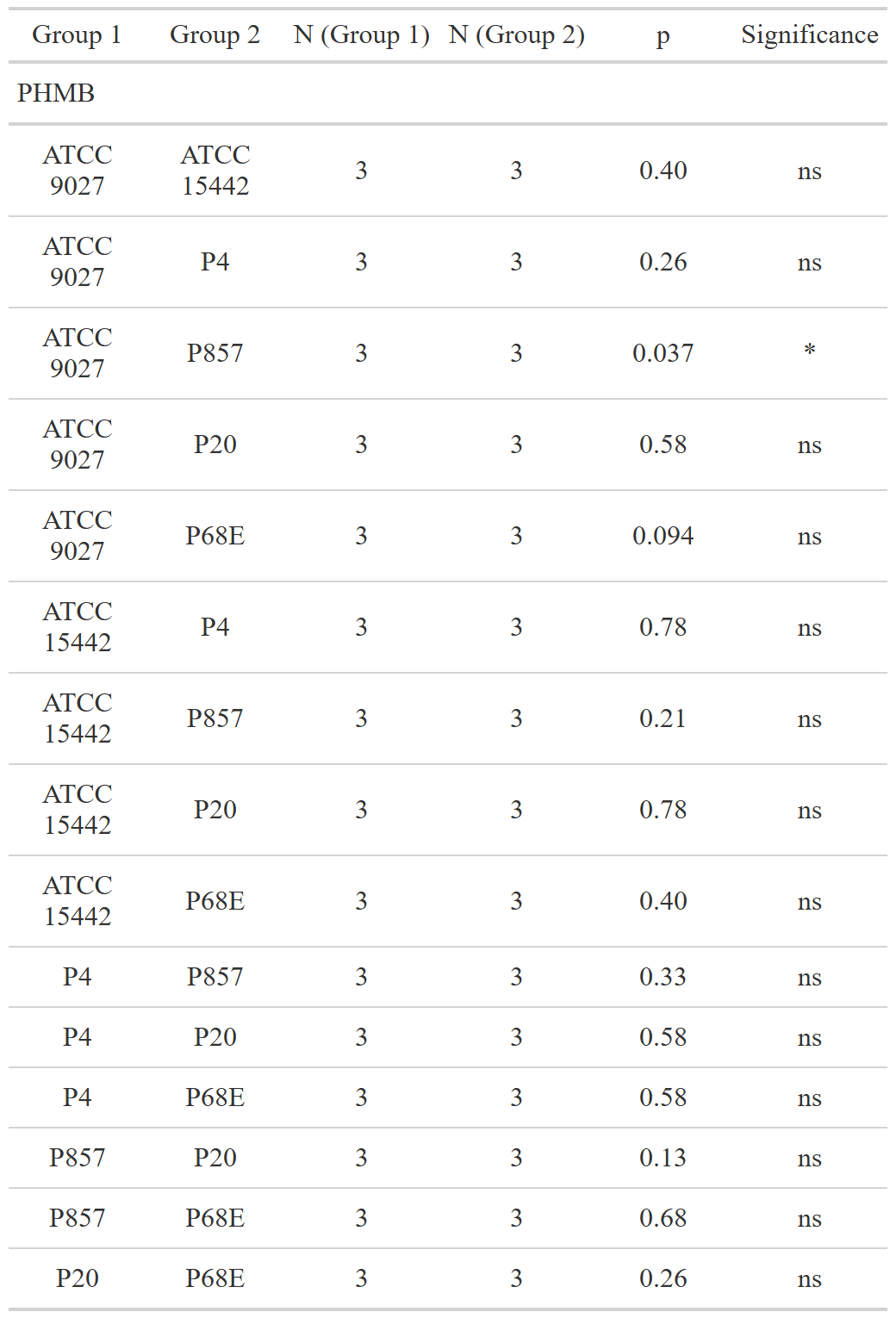

**Supplementary Table S9:** Parameters of the tested statistic of the differences between zones of growth inhibition (mm) of tested compounds against *P. aeruginosa* strains. Dunn’s test, followed by the Kruskal-Wallis test, was performed. Values of p<0.05 were considered significant. Only significant differences were included, p<=0.05 was marked with one asterisk. N- data points, TEO- Thyme Essential Oil, PHMB- polyhexanide.

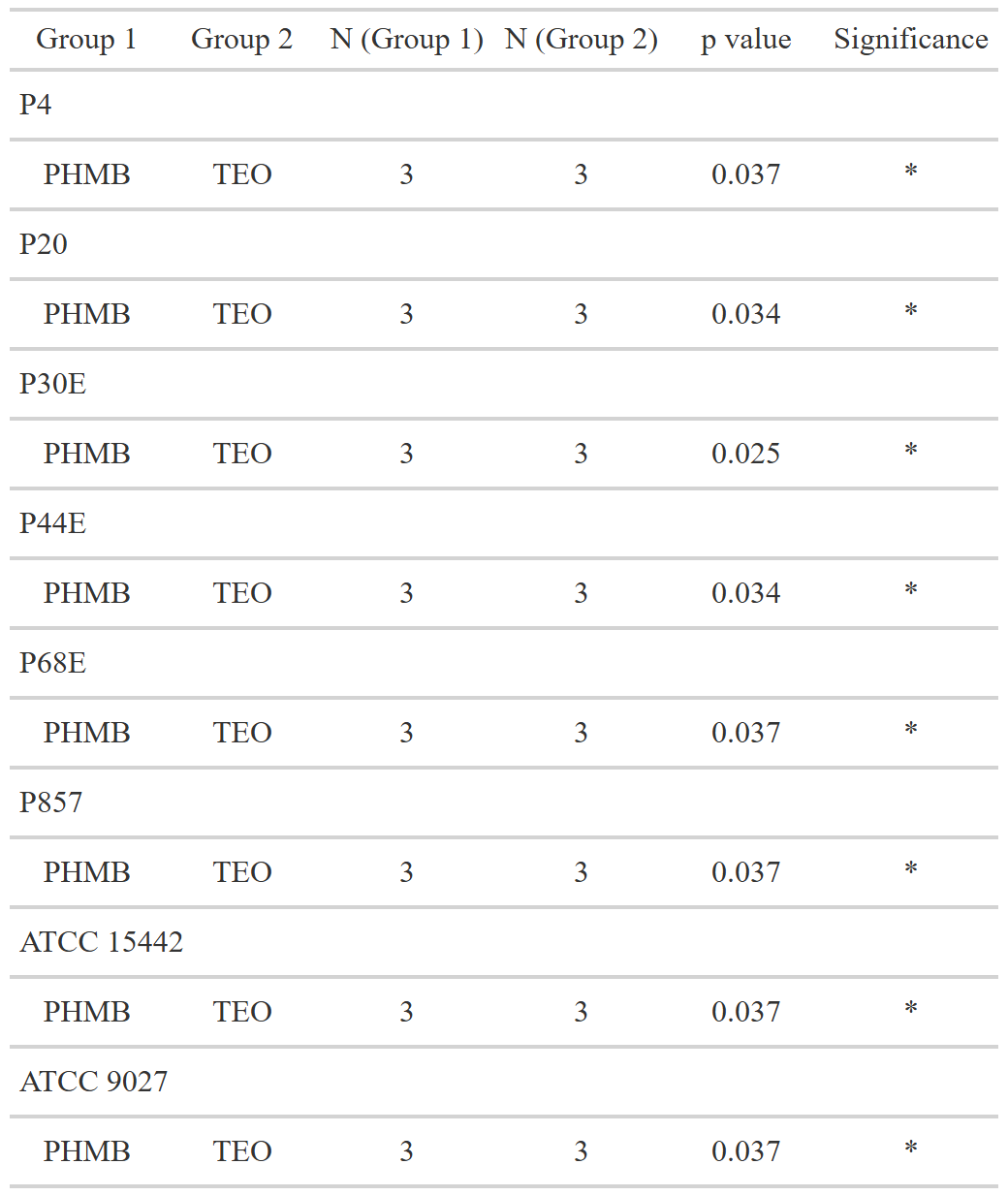

**Supplementary Table S10:** Parameters of the tested statistic of the differences between Minimal Inhibitory Concentration (MIC) (%, v/v) values of tested compounds against *P. aeruginosa* strains. Dunn’s test, followed by the Kruskal-Wallis test, was performed. Values of p<0.05 were considered significant.  Only significant differences were included, p<=0.05 was marked with one asterisk. N- data points, TEO- Thyme Essential Oil, PHMB- polyhexanide.

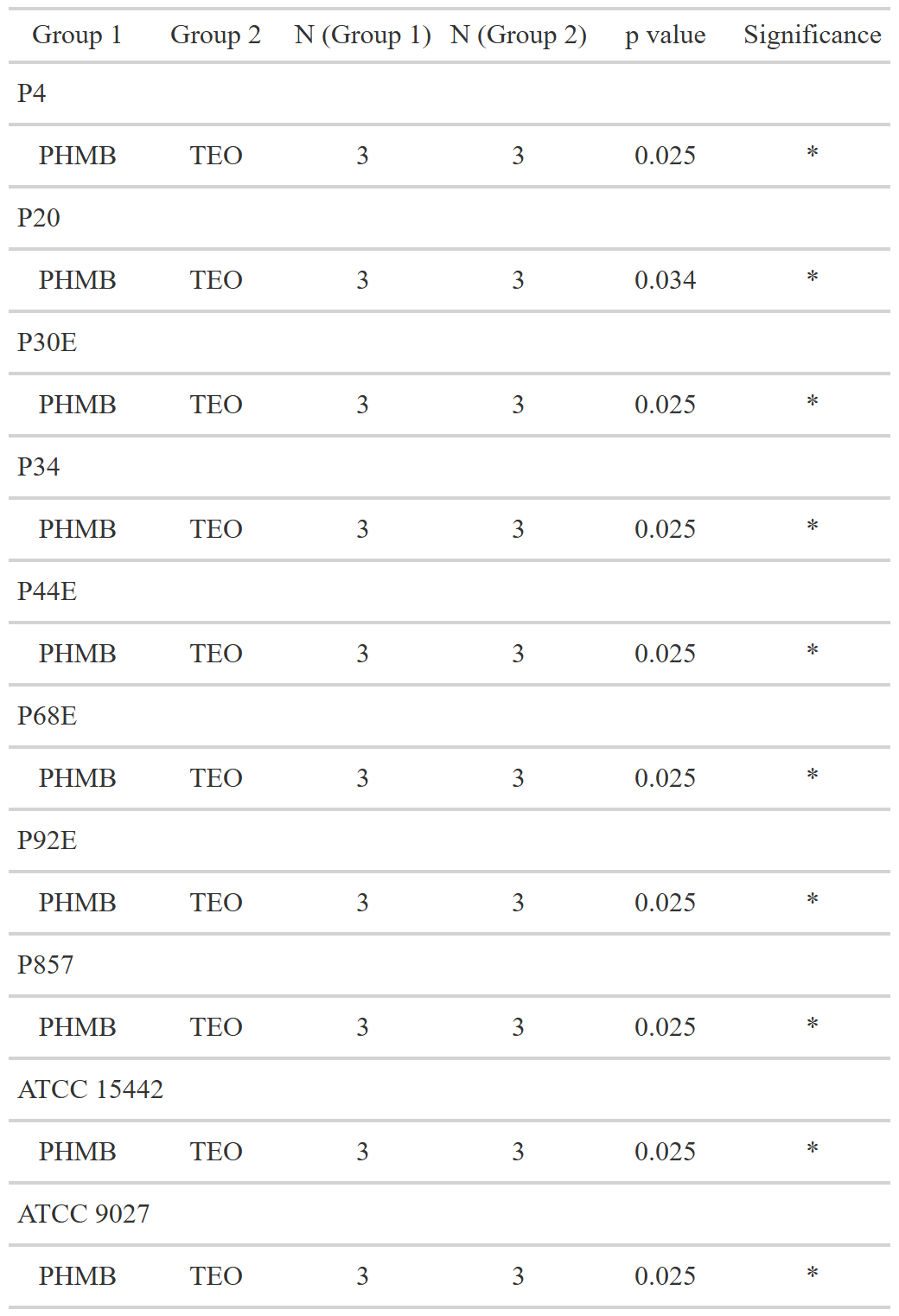

**Supplementary Table S11:** Parameters of the tested statistic of the differences between antibiofilm activity of tested compounds against *P. aeruginosa* strains. Dunn’s test, followed by the Kruskal-Wallis test, was performed. Values of p<0.05 were considered significant.  Only significant differences were included, p<=0.05 was marked with one asterisk. N- data points, TEO- Thyme Essential Oil, PHMB- polyhexanide.

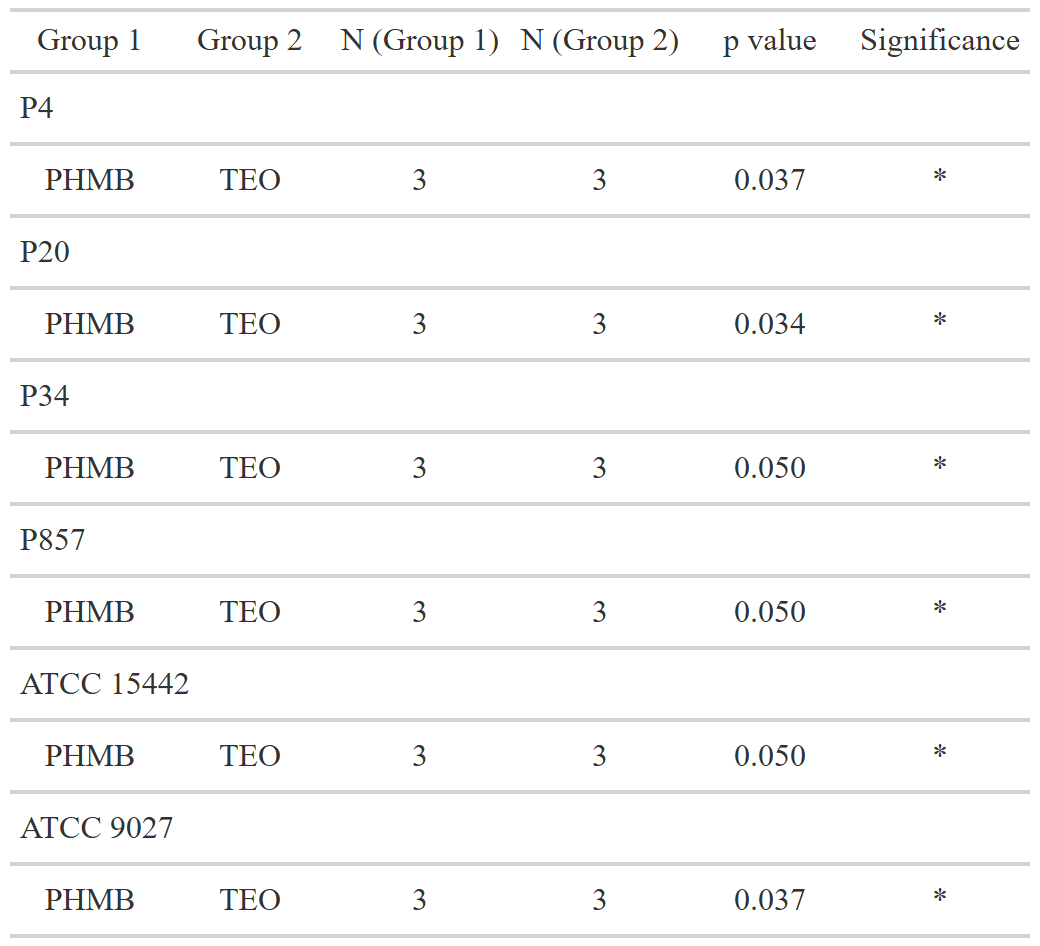

**Supplementary Table S12:** Statistical analysis of the data distribution for biofilm mass (a) or metabolic activity (b) or Colony-Forming Unit number (CFU/mL) (c) across genetically distinct groups of *P. aeruginosa* strains. Normal distribution was considered for values of p>0.05 (Shapiro-Wilk test). SD- standard deviation, IQR- interquartile range (IQR), N- data points, n- number of strains included in each group.

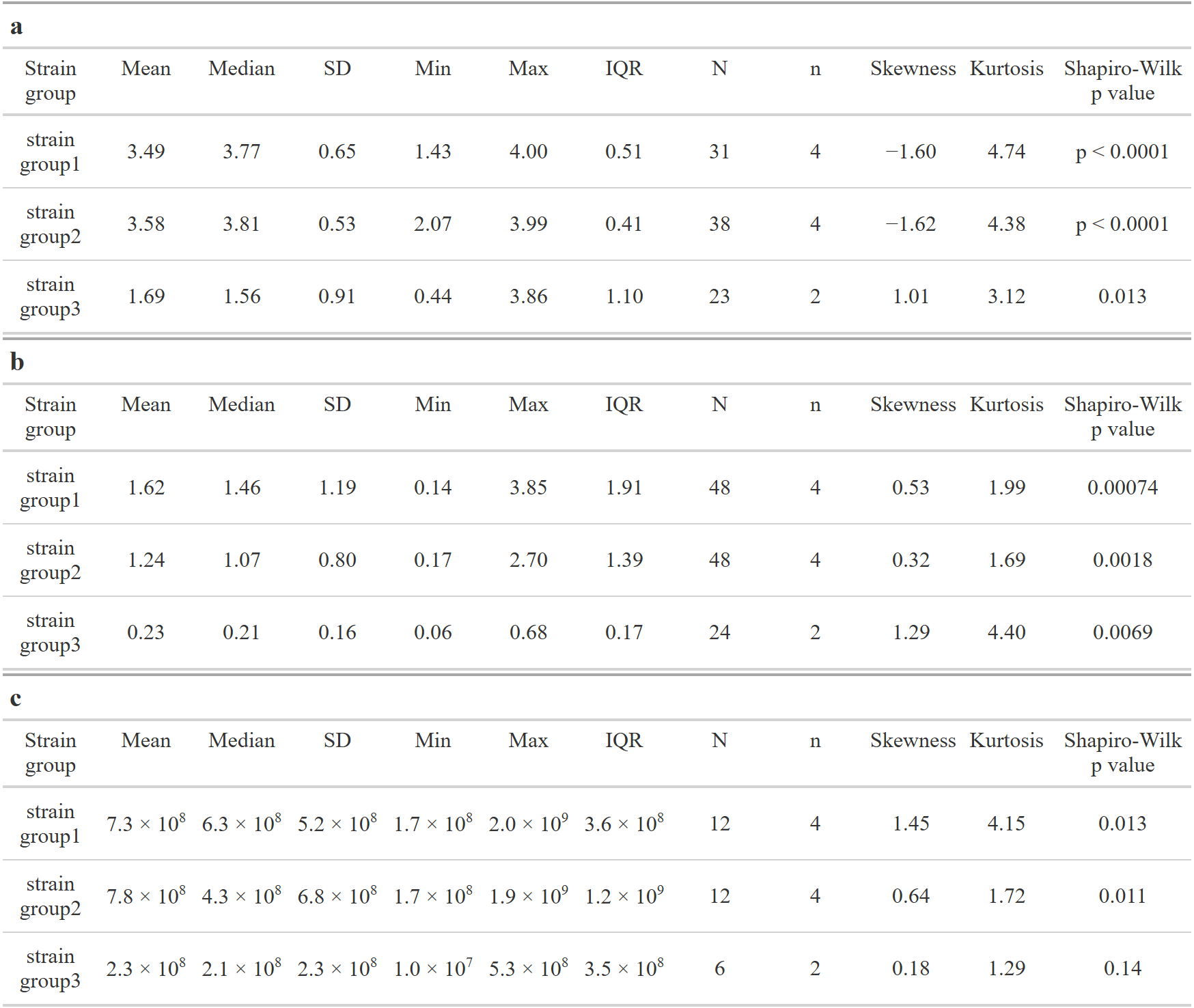

**Supplementary Table S13:** Statistical analysis of the data distribution for antimicrobial and antibiofilm activity of tested compounds against genetically distinct groups of *P. aeruginosa* strains. Growth inhibition zones values (mm) (a); Minimal Inhibitory Concentration (MIC) (%, v/v) (b); Biofilm cells reduction (%) (c). Normal distribution was considered for values of p>0.05 (Shapiro-Wilk test). SD- standard deviation, IQR- interquartile range (IQR), N- data points, n- number of strains included in each group,NaN/ NA- not applicable. TEO- Thyme Essential Oil, PHMB- polyhexanide C+- control with Phosphate-Buffered Saline.

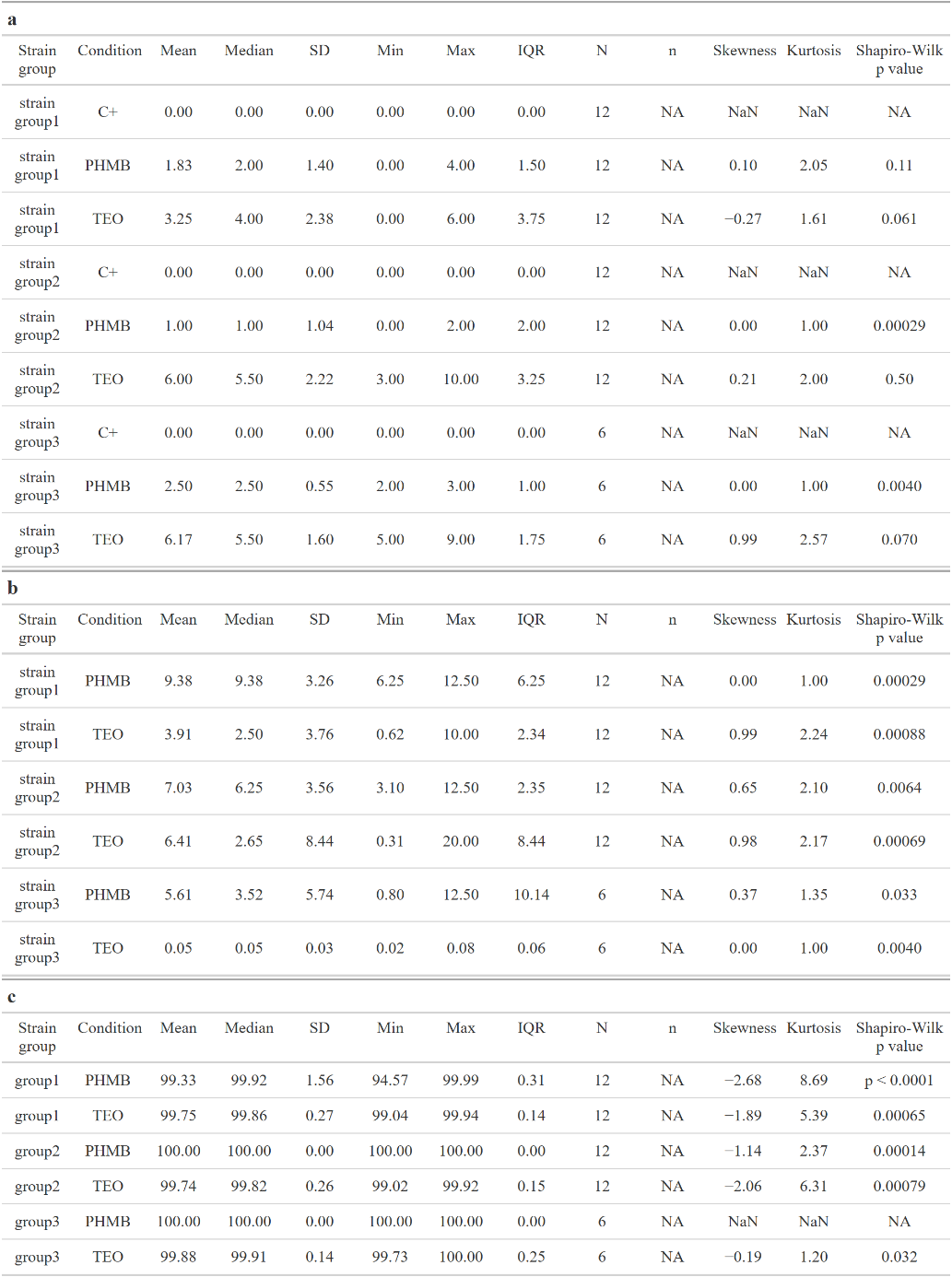

**Supplementary Table S14:** Statistic data of differences in biofilm features between *P. aeruginosa* genetically distinct groups. Biofilm mass (a); biofilm metabolic activity (b). Values of p<0.05 were considered significant (Dunn’s test), p<=0.0001 was marked with four asterisk. Ns- no significant differences, N- data points, n- number of strains included in each group. For biofilm Colony-Forming Unit number (CFU/mL) p values between all groups were no significant.

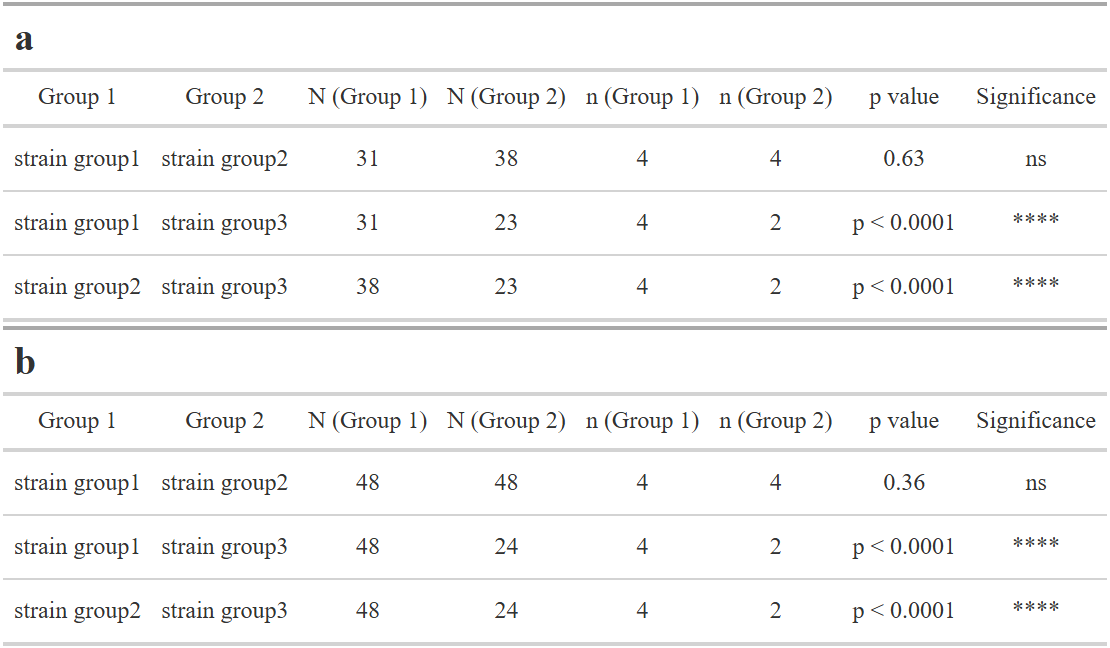

**Supplementary Table S15:** Statistic data of differences for antimicrobial and antibiofilm activity of tested compounds against genetically distinct groups of *P. aeruginosa* strains. Inhibition zone values (mm) (a); Minimal Inhibitory Concentration (MIC) (%, v/v) (b); biofilm cells reduction (%) (c). Values of p<0.05 were considered significant (Dunn’s test), p<=0.05 was marked with one asterisk, p<=0.01 was marked with two asterisk, p<=0.001 was marked with three asterisk, and p<=0.0001 was marked with four asterisk. Ns- no significant differences, N- data points, n- number of strains included in each group. TEO- Thyme Essential Oil, PHMB- polyhexanide.

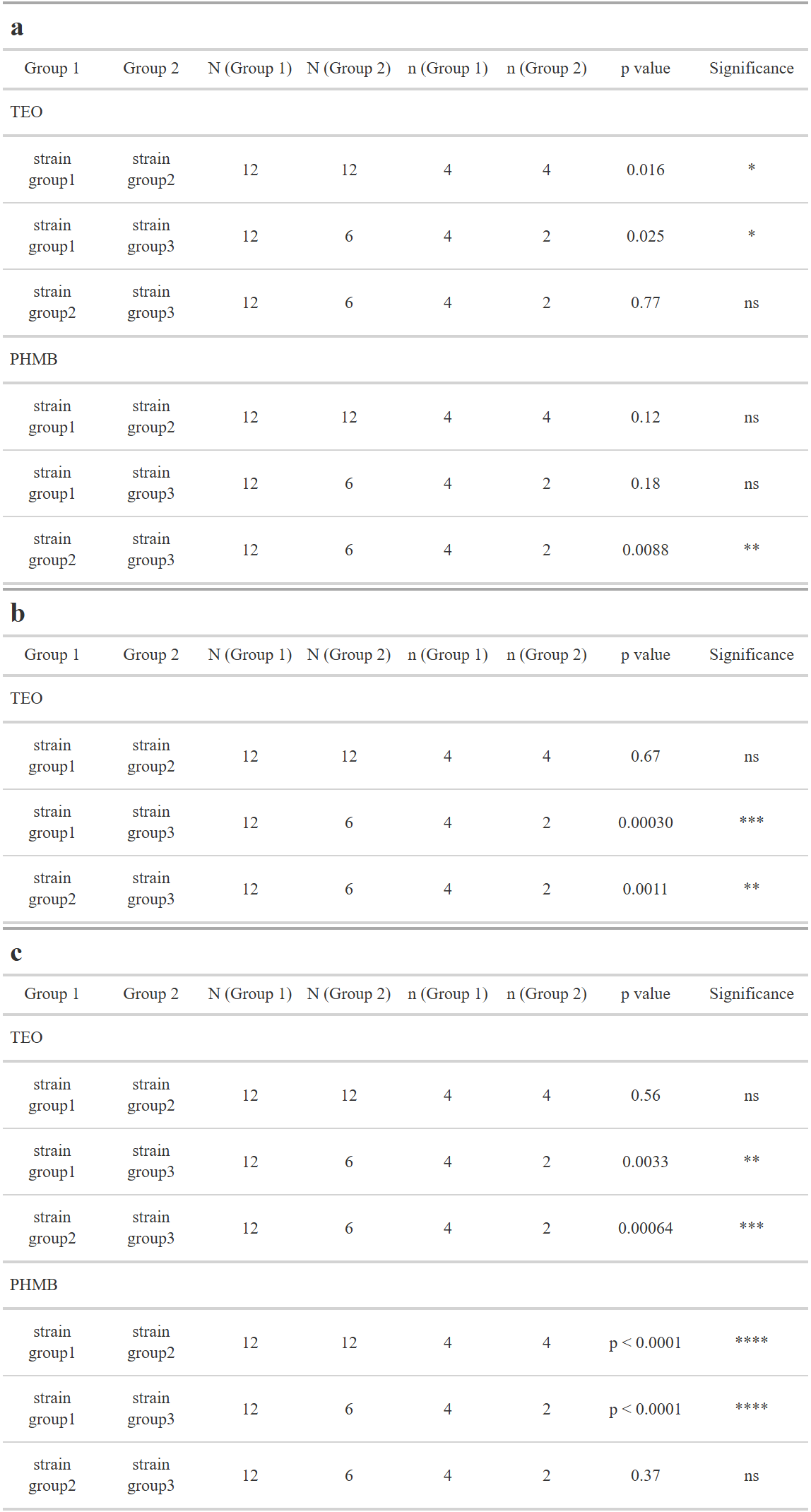
